# Single-cell spatial mapping reveals reproducible cell type organization and spatially-dependent gene expression in gastruloids

**DOI:** 10.1101/2025.07.14.664617

**Authors:** Catherine G. Triandafillou, Pranav Sompalle, Yael Heyman, Arjun Raj

## Abstract

Gastruloids are stem-cell-based models that recapitulate key aspects of mammalian gastrulation, including formation of an anterior-posterior (AP) axis. However, we do not have detailed spatial information about gene expression and cell type organization, particularly at the level of individual gastruloids. Here, we report a spatially resolved, single-cell molecular catalog of the transcriptomes of 26 individual gastruloids. We found that cell type composition and tissue-scale spatial organization were largely consistent across gastruloids, but meso-scale patterning of specific cell types varied between samples. Posterior cell types formed distinct, organized clusters, while anterior cell types were more disorganized. To distinguish progressive differentiation from cell type differences, we developed the L-score, a parameter-free quantification of mutually exclusive gene expression. This analysis revealed spatial organization without explicit encoding, recapitulated known cell type relationships, and identified novel gene expression states and spatial subclusters within cell types. We confirmed that in gastruloids, NMP differentiation occurred through a continuous, spatially-coordinated process. We also showed that endothelial precursors exhibited unique spatial organization and had distinct gene expression profiles dependent on their association with anterior somitic or posterior endodermal tissues. This work enables the rigorous use of gastruloids as models for studying the molecular mechanisms underlying mammalian development and tissue organization, and introduces new computational tools for analyzing spatially-resolved single-cell datasets.

## Introduction

Embryogenesis involves not only the creation but, crucially, the organization of new cells and new cell identities. Gastruloids model an important stage of embryogenesis: gastrulation. Gastruloids break symmetry, elongate along an anterior-posterior (AP) axis, and form many of the cell types found in embryos at a similar developmental stage, as shown by single-cell RNA-seq analyses (Beccari et al., 2018; Turner et al., 2017; van den Brink et al., 2020, 2014; van den Brink and van Oudenaarden, 2021; Veenvliet et al., 2020). These characteristics mean they could hold enormous potential to enable us to deeply characterize, perturb, and test developmental hypotheses, circumventing many practical and ethical constraints imposed by using *in vivo* models. Yet despite extensive single-cell characterization of gastruloids, we still lack detailed information about the patterning of cell types within individual gastruloids.

Sequencing-based gene expression analyses of gastruloids have at most been able to generate coarse-grained maps of the spatial arrangement of the constituent cells via sectioning (Moris et al., 2020; van den Brink et al., 2020). These methods cannot resolve the position of individual cells, nor can they measure how tissues are organized other than along the AP axis. Computational methods that measure gene expression and identity of individual cells can only estimate AP axis location through inference methods and do not measure organization in any other dimension (Harland et al., 2025; Yaman and Ramanathan, 2023). Additionally, detailed comparative spatial analyses of individual gastruloids have not been performed, as the experiments either require pooling gastruloids or are low-throughput, such that there are insufficient samples to make meaningful comparisons. While some studies point to stringent reproducibility of gene expression along the AP axis in gastruloids (Bennabi et al., 2024; Merle et al., 2024), others show that there is great variation, gastruloid-to-gastruloid, in the organization of specific tissues (Farag et al., 2023). Without high-resolution spatial characterization across multiple gastruloids, it is unclear what aspects of tissue development can be effectively modeled with this system.

The cell types present in gastruloids have largely been inferred from single-cell sequencing analyses. As is standard for these experiments, cells are clustered on the basis of their overall gene expression profiles, and types are assigned based on a qualitative assessment of the genes differentially expressed by the cluster (McNamara et al., 2023; Moris et al., 2020; Regalado et al., 2025; Rosen et al., 2022; van den Brink et al., 2020). However, it is unclear whether discrete binning cells into categories is the only, or even the most useful, approach to single-cell studies of gastruloids, a system nearly entirely composed of cells in the process of differentiating and transitioning between identities. A more gene-centric analysis, especially one coupled to a spatial measurement, could more naturally represent these transitions and how they are coupled to the arrangement and physical location of cells. Whether the diversity of cell types observed in sequencing experiments truly maps onto spatially organized collections of cells, and if this organization is consistent between gastruloids, remains an open question.

To address these gaps, and to create a systematic, high resolution dataset of gene expression in gastruloids considered to be morphologically normal, we developed a spatially resolved, single-cell molecular map of the location, identity, and gene expression of cells within 26 individual gastruloids with normal morphologies. We found that despite some morphological variability within the qualitative category of elongated and polarized, “normal” gastruloids had largely reproducible cell type composition. The order of types and expression of individual genes along the AP axis was also quite consistent. Leveraging our spatial single-cell expression data, we found that the posterior of the gastruloids was characteristically patterned with distinct clusters of progenitors and differentiated cell types. The degree of this clustering varied between samples, as did the mesoscale organization of cell types found near the middle of the AP axis. Some, but not all, of these differences were explainable by variation in morphology. We present novel computational techniques useful for identifying genes with mutually exclusive expression patterns, which illuminated the spatial component of a bipotent progenitor differentiation decision and revealed a spatial hierarchy of gene expression patterns. Using evidence generated from this measure, we show evidence of spatial and functional specialization of endothelial cells, suggesting that gastruloids may be used in the future to model early stages of vasculogenesis and organization.

## Results

### Cell types’ locations and relative proportions are consistent across morphologically normal gastruloids

To measure the spatial distribution of gene expression, we prepared gastruloids using mouse E14TG2a cells and a standard protocol (see Methods). We harvested mature gastruloids after 120 hours of growth. The experiment was performed 3 times on different days, so to ensure consistency we checked that the proportion of the gastruloids that formed correctly was the same or greater than the median of all experiments (Figure **S1.1a**). Although there was variation in the length, width, and relative amounts of anterior and posterior tissues in the gastruloids considered, they were within the range of what would qualitatively be considered a ‘morphologically normal’ gastruloid (Merle et al., 2024; van den Brink et al., 2014).

To address potential batch effects due to biological differences between runs, we examined brightfield images of all the gastruloids generated for each experiment (529 total gastruloids across 6 plates on 3 different days), segmented them, and quantified morphological characteristics. When we embedded all 529 gastruloids into PCA space, there was near complete overlap between all groups, with the exception of one plate from 9/1/2024, which was slightly higher in PC1. **Figure S1.1b** shows this embedding, and examples of gastruloids at the extreme ends of PCs 1 and 2. We note that the samples collected on 9/1/2024 were on average smaller than those from the other two experiments, but spanned the same range of elongation (**Figure S1.1c**). Interestingly, the final size as measured by cross sectional area of a brightfield image of the gastruloid did not correlate with the initial seeding number (the experiment on 4/4/2025 used 100 starting cells and the other two experiments used 300). Previous studies have demonstrated that the gene expression differences between gastruloids seeded with 100 and 300 cells is extremely small (Bennabi et al., 2025).

We embedded fixed gastruloids whole-mount in polyacrylamide gel and assayed with seqFISH (Eng et al., 2019). An example of raw images showing spots and nuclear quality, as well as deconvolved spots for an example gene Dll1, is shown in **Figure S1.1d**. To assess the quality of the data, we first assigned an AP axis to each gastruloid using the expression of T, a canonical marker for the posterior (**Figure 1a**). We compared how gene expression varied along the AP axis, and saw good agreement at a coarse-grained level with a previous study that sectioned gastruloids along the axis and analyzed gene expression in each section (van den Brink et al., 2020) (**Figure S1.2a**). The colinearity of the peak expression of Hox genes in our panel was also consistent with this dataset, with a median Pearson correlation of 0.695 (compared to 0.663 for all genes, **Figure S1.2b**).

**Figure 1:**
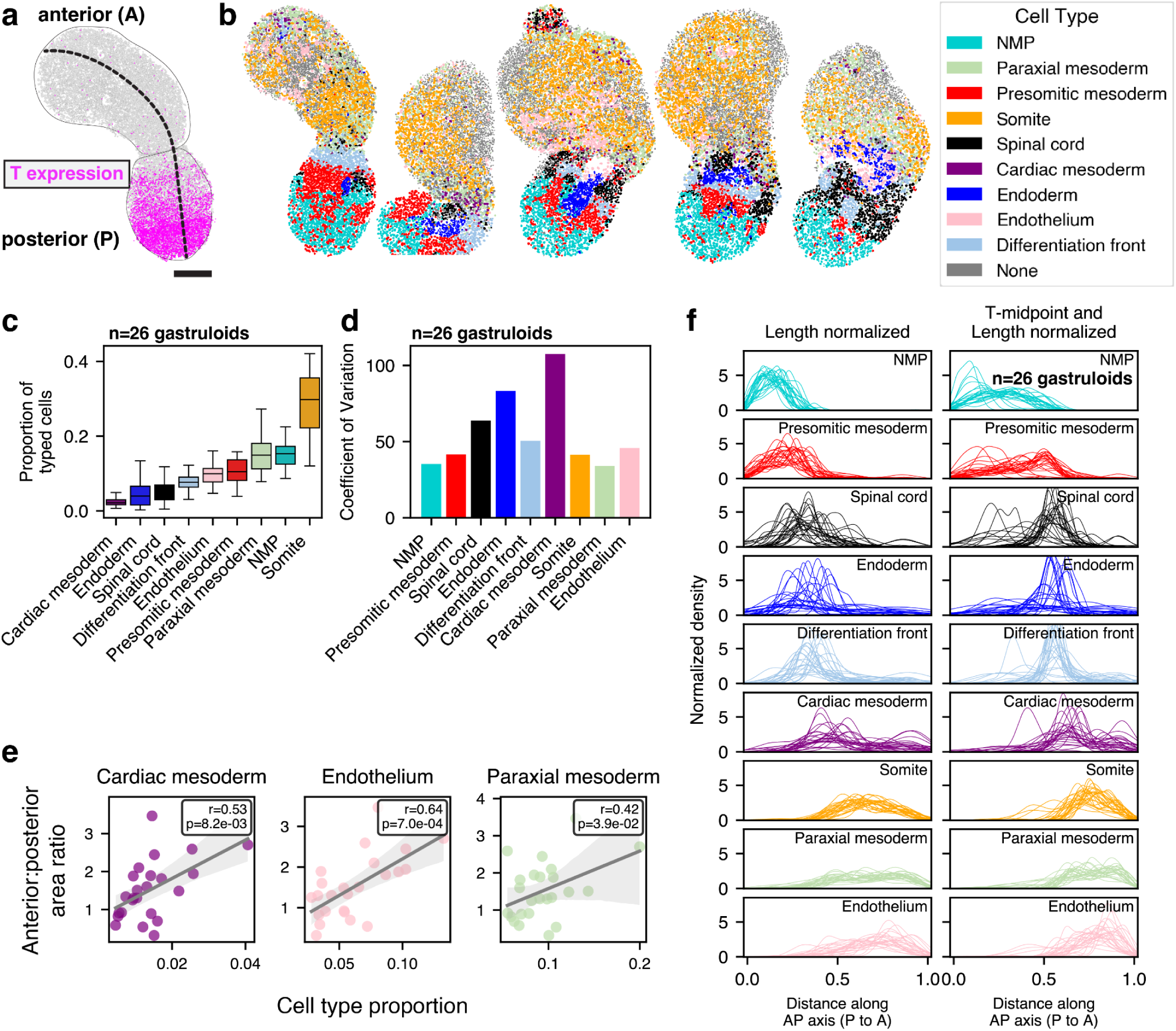
Cell types’ locations and relative proportions are consistent across morphologically normal gastruloids. **a.** Representative gastruloid with the expression of T shown in magenta. Each spot is one transcript (count), nuclei are shown in gray. Thick dashed line denotes the AP axis, thin lines outline the anterior and posterior halves of the gastruloid. Scale bar is 200 μm. **b.** Gallery of representative gastruloids. Scale bar is the same as **a**. **c.** Proportion of each cell type across all gastruloids, ordered by median proportion. **d.** Coefficient of variation for distributions shown in **c**. **e.** Correlation between the ratio of anterior area to posterior area across gastruloids versus the proportion of specific cell types. Cell types shown had a statistically significant (p < 0.05) correlation; all cell types are shown in **Figure S1.4e**. **f.** Smoothed density of each cell type along the AP axis. Nuclei in each gastruloid were projected onto the AP axis, and their position was normalized to the total length (left) or symmetrically around a midpoint defined by T expression (right). Traces from individual gastruloids are shown by individual curves.

To profile the spatial organization of individual gastruloids, we assigned a cell type to each nucleus using a cell type scoring method with known marker genes. We compared these results to those obtained with unsupervised clustering. We found that although clustering did produce clusters, they were not strongly separated and we were not able to individually resolve some cell types we expected to find: specifically, there was a mixed differentiation front and presomitic mesoderm cluster, and NMP and spinal cord cells were also clustered together (**Figure S3.6a**). Therefore, we proceeded with the scoring-based method; see Methods for additional details. A representative gallery of typed gastruloids is shown in **Figure 1b**; the full dataset is in **Figure S1.3**. Because each cell received a cell type score for each type, we could use the entropy of the cell type score probability distribution to assess confidence of our cell type assignment: a cell that received a similar score for multiple cell types would have a high entropy distribution, while one which scored highly for one type and low for the rest would have low entropy. On average, the cell type entropies for most cells in a given cell type were low, with the exception of paraxial mesoderm and cardiac mesoderm cells, which had intermediate values (see additional discussion below). The generally low entropies indicate that most of our cell type assignments were high-confidence (**Figure S1.4a**).

The seqFISH technique is imaging-based, and the fact that we image transcripts in a single plane admits the possibility that due to rotational differences, different parts of the gastruloid are imaged in each sample. We controlled for this to the extent possible within experimental limitations by imaging multiple gastruloids across several experiments and keeping our imaging parameters, particularly the instrument z-depth relative to the coverslip, nearly identical across experiments. We also note that previous analysis of 2D and 3D distances has indicated that in many cases, 2D distances (as we use in this work) are preferable for making comparisons between cells in a sample (Finn et al., 2017).

Once we had the cell type identity and spatial location of each cell in all the gastruloids, we first qualitatively examined where each cell type was found relative to other types and overall morphology. The posterior region, although variable in size (**Figure S1.3a,b,c**), mainly consisted of neuromesodermal precursors (NMP, turquoise), a bipotent cell type that contributes to both neural and mesodermal tissues (Gouti et al., 2014; Henrique et al., 2015; Tzouanacou et al., 2009). This localization is consistent with previous observations of the expression of NMP genes in the gastruloid posterior (Moris et al., 2020; Underhill and Toettcher, 2023; van den Brink et al., 2020). However, unlike these previous studies, which inferred NMPs from the expression of single genes and could not resolve other cell types, we were able to see that NMPs were intermixed in a patchy fashion with mesodermally-fated cells (presomitic mesoderm, PSM, red). In some instances, we saw PSM cells forming symmetric patches on either side of a central neurally-fated region (**Figure 1b**, center gastruloid, **Figure S1.3a v., xi., xii, b iv.**), mimicking the organization found in embryos (Gossler and Tam, 2002; Tam and Behringer, 1997; Xie et al., 2025) and thought to occur in gastruloids (Yamanaka et al., 2023; Yaman and Ramanathan, 2023). Generally, we saw 1-2 distinct clusters of neurally fated cells (spinal cord precursors, black), often adjacent to or opposite the region associated with somitic fate acquisition (“differentiation front”, light blue), although occasionally found in the center of the gastruloid (**Figure S1.2a ii., vi., xvii**). Although distinct organization of neurally fated cells has been observed in other embryo models (Veenvliet et al., 2020), the organization of spinal cord precursors in gastruloids in the context of other cell types has not yet been systematically characterized; staining for spinal cord-associated genes showed compelling patterning, but the patterns varied by gene, complicating interpretation of the organization of the tissue (Beccari et al., 2018). Consistent with (van den Brink et al., 2020), the differentiation front was nearly always found in one distinct cluster marking the boundary between the anterior and posterior (**Figure S1.3a i., iii., vi., b iii., vi**). The anterior regions consisted mainly of somitic cells, presomitic cells, and cardiac mesoderm (yellow, green, and purple) (**Figure S1.3 a iii., v., x., xv., xvi.**). We also detected endoderm cells, although not in all gastruloids, consistent with observations in (Farag et al., 2023). This simultaneous mapping of all cell types is qualitatively consistent with many observations in the literature about the expression of individual genes and integrates them into one consistent picture of cell type organization.

We sought to quantify variability in cell type composition between the 26 morphologically normal gastruloids profiled. Previous single-cell datasets relied on pooling multiple gastruloids, thus obscuring the degree to which the overall cell type distribution was reflected in each individual gastruloid. However, recent single-cell measurements of individual gastruloids have suggested substantial gastruloid-to-gastruloid variation in cell type proportions (Rosen et al., 2022). **Figure 1c** shows distributions of cell type proportions across samples, and **Figure 1d** shows the coefficient of variation of these proportions. Individual gastruloid cell type distributions, including the proportion of cells that had insufficient reads to be confidently assigned a type, are shown in **Figure S1.4b,c**. We found that cardiac mesoderm, endoderm, and spinal cord cells had the greatest coefficient of variation in proportion between gastruloids (**Figure 1d**). To calculate statistical significance, we first performed a centered log-ratio (CLR) transform on the proportions, then looked for covariation between cell types across gastruloids. We found there was a statistically significant inverse correlation between the proportion of endoderm and NMP, presomitic mesoderm, and differentiation front (**Figure S1.4d**). We did not observe gastruloids that were as strongly neurally-biased as those reported in (Rosen et al., 2022), but we did see some gastruloids with a relatively high proportion of spinal cord precursor cells (**Figure S1.3a ii., xv., b vii.**), and overall the proportion of spinal cord had a negative covariation with the mesodermally-derived cell types, consistent with the anticorrelation also reported in (Rosen et al., 2022) (**Figure S1.4**).

The proportion of presomitic mesoderm cells was significantly positively correlated with the proportion of somite cells (covariation = 0.63, **Figure S1.4d**). Presomitic mesoderm cells progressively take on somitic fate at and after the differentiation front; a positive correlation between these two would suggest that the amount of presomitic mesoderm is linked to how much somite is ultimately formed. How much presomitic mesoderm is formed depends on fate decisions made by NMPs: NMP progeny can become either mesodermal or neural, depending on their local signaling environment. This chain of correlation suggests that fate decisions made by NMPs may drive the overall composition of individual gastruloids and that, as in embryos, a balance between the amount of neural and mesodermal tissue is achieved, although likely to a less stringent degree (Garriock et al., 2015). Further experiments could delineate whether, as in embryos, Wnt3a drives this balance (Garriock et al., 2015), and the degree to which growth of differentiated progeny affects compositional variation (Underhill and Toettcher, 2023).

Finally, we quantified correlations between morphology and cell type proportion: we used the segmented images of the gastruloids to generate an anterior to posterior area ratio, and looked for correlation between this ratio and the proportion of individual cell types. We found that only cardiac mesoderm, endothelium, and paraxial mesoderm cells exhibited significant correlation with A:P ratio, and in all cases the correlation was positive (**Figure 1e**, all cell types in **Figure S1.4e**).

### Cell types’ arrangement along the AP axis is highly consistent across gastruloids

Given that the proportions of cell types within each gastruloid were fairly consistent, to what degree did their spatial organization vary? We first projected each cell of each type onto the AP axis and looked at the distribution of AP axis locations across gastruloids (**Figure 1f**, left hand side). The most posterior cell types (NMP and presomitic mesoderm) showed wide distributions from 0-30% of the axis. Centered around 30%, spinal cord precursors and endoderm had distinctive peaks (clusters), the exact location of which varied between gastruloids. Using a threshold of T expression to define the midpoint of each gastruloid and uniformly length-normalizing each half caused these cell types to collapse into a single peak (**Figure 1f**, right hand side). The differentiation front was similarly located in one peak (between 30-40% of the AP axis), and the variation in the location of this peak decreased with T-expression normalization. To quantify this variability, we bootstrapped a null distribution of cell type locations by pooling each cell type together across samples and using the resulting distribution to define a reference mean. When we compared how much peak variation there was among samples randomly drawn from this null to our observed data, we found that although nearly every cell type (excepting paraxial mesoderm) had more variability than expected by chance, the effect size of this variation was small (**Figure S1.5a**). Normalizing location to T-expression decreased variability in most cell types, most notably for differentiation front and spinal cord (**Figure S1.5b**).

We also asked how the order of cell types along the AP axis varied between gastruloids. When we ranked the peaks shown in **Figure 1f** per gastruloid, we found that Kendall’s W, an overall measure of rank coherence across independent samples that spans from 0 (no agreement) to 1 (complete agreement), was 0.834 (**Figure S1.5c**). We found that the cell types most likely to swap rank order were spinal cord and endoderm, and paraxial mesoderm and endothelium (**Figure S1.5d**).

### Overall cell type mixing varies between gastruloids

Several studies of gene expression in gastruloids have used pooled measurements to infer the AP axis-location of genes and cell types (Anlas et al., 2025; Moris et al., 2020; Rosen et al., 2022; Veenvliet et al., 2020; Yamanaka et al., 2023) and our data are largely consistent with these lower-resolution findings. Yet it is obvious from individual gene staining (Beccari et al., 2018; Braccioli et al., 2022; van den Brink et al., 2014; Yaman and Ramanathan, 2023) and our detailed 2D maps of cell identity and location that gastruloid organization is much more complex than the average order of cells along the AP axis. We thus needed an analytical method for quantifying spatial organization beyond distributions along the AP axis. To further characterize spatial organization, we sought to quantify the degree to which cells were mixed in each gastruloid, and how that mixing might vary between gastruloids. For each cell in each gastruloid, we counted the interactions between that cell and all its neighbors within a 16 μm radius (on average 5-6 neighbors per cell), and summarized all these interactions for all cells in the gastruloid in a matrix, normalizing each element by the frequency of the cell type considered to be the ‘neighbor’ in the interaction.

To quantify overall mixing, we calculated the sum (across types) of the frequency of self interactions, normalized to the total interactions and then took the inverse. We normalized this value to the expected cross-type interactions predicted from random mixing (i.e. the proportion of the neighboring cell type). This gave us, for each gastruloid, a single value that we call the mixing index. The mixing index ranged from -1 (totally segregated) to +1 (totally mixed with less frequent self-interactions than expected from chance); a score of 0 indicates a random distribution, i.e. neighbor frequency is exactly what would be predicted by that cell type’s frequency alone. The mixing index values range from -0.50 to -0.22 (**Figure 2a**). While all values were negative, the range was large. This observation led us to conclude that while in all the gastruloids profiled cell types tended to cluster together, there was variation between gastruloids in the degree of coherent clustering between types.

**Figure 2:**
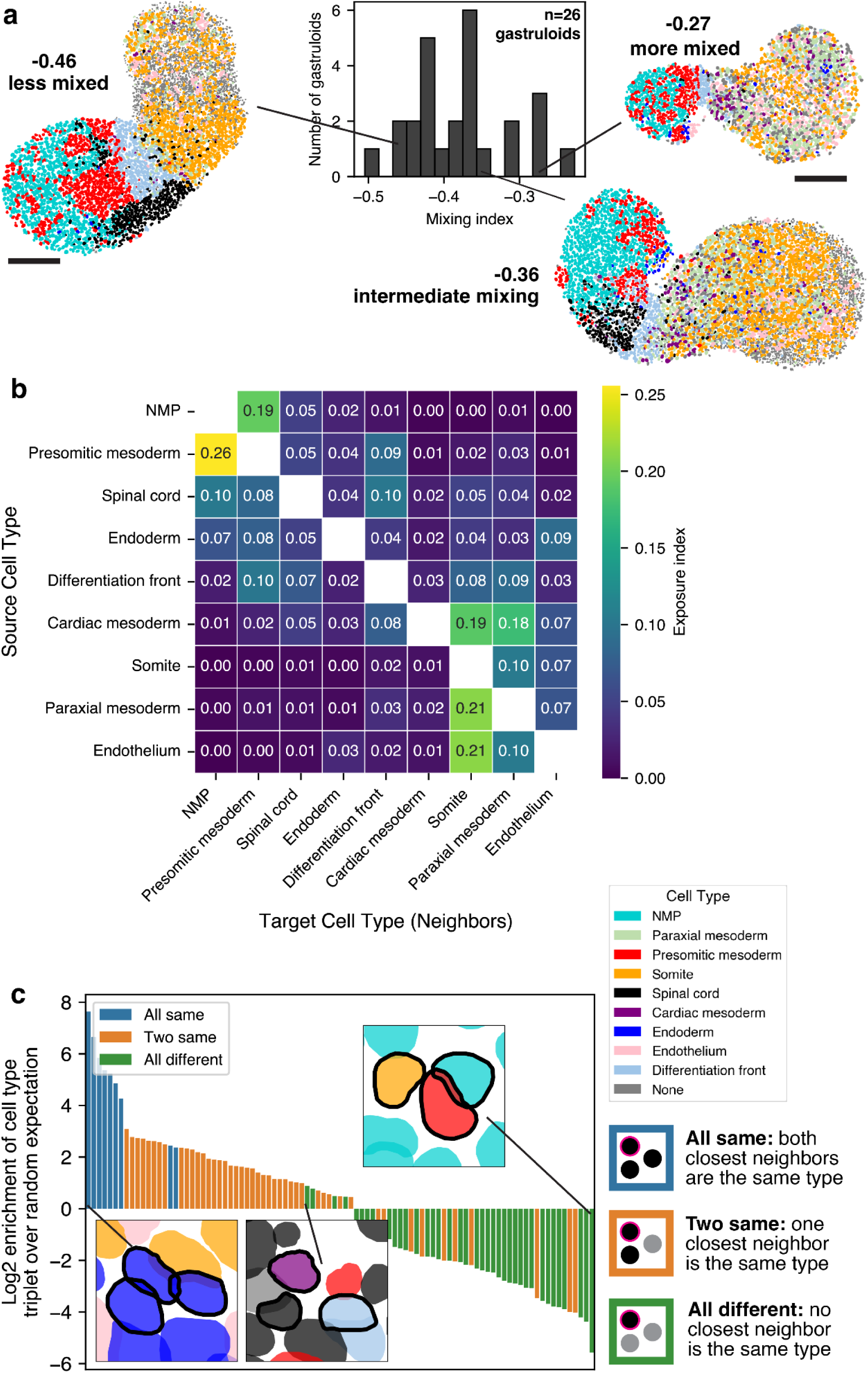
Pairwise cell type interactions quantify cell type mixing and interaction motifs. **a.** Distribution of mixing indices across gastruloids. **b.** Exposure index, which represents how frequently the source cell type (row) is found next to the target cell type (column). The maximum possible value is 1, self-interactions excluded for readability. **c.** Significantly enriched and depleted triplet combinations of cell types, ordered by effect size from left to right. Bars are colored by the composition of the triplet they represent. Only triplets found in over half of the gastruloids are shown.

### Pairwise cell type interactions suggest more cell type mixing in the gastruloid anterior than posterior

The interaction between long-range signaling gradients and short-range cell-cell interactions is thought to drive gastruloid symmetry breaking and patterning (Arias et al., 2022; Gupta et al., 2021; Simunovic and Brivanlou, 2017), and our spatial data allowed us to query both micro and meso-scale organization. Our simultaneous observation of variability in mixing index and consistent cell type organization along the AP axis led us to ask if mixing was driven largely by local topology and whether this effect was disproportionately due to the arrangement of a subset of cell types. We noted that the off-diagonal elements of the matrix we used to calculate the mixing index were informative about cell type-cell type interactions. Specifically they quantify the degree to which each cell type (source) is exposed to another cell type (neighbors). To assess the overall frequency of cell type-cell type interactions, we first pooled the data from all gastruloids together into one interaction matrix, the elements of which we refer to as the ‘exposure index’. (**Figure 2b, Figure S2.1a,b**). Higher values indicate a greater frequency of being found in close proximity (see Methods for full details). The matrix is asymmetric in that the exposure of cell type A to B may not be the same as the exposure of cell type B to A. For example, a cell type found in singlets within a larger tissue will be more frequently exposed to cells of that common tissue type, while the common type will see itself much more frequently than the intermixed cell. Thus, this asymmetry accounts for the fact that local patterning can influence the degree to which any one cell type is found next to another. For instance, somite and endothelial precursors have an asymmetric pattern: although endothelial cells are often exposed to somite cells (exposure index 0.21), somite cells are comparatively infrequently exposed to endothelial cells (0.07). This observation, coupled with a fairly strong endothelial-endothelial exposure index (**Figure S2.1b**), suggests small clusters of endothelial cells are found among somite cells, reflecting our visual impressions. In contrast, cardiac mesoderm cells were often exposed to all other anterior cell types, but their self-exposure index was low (**Figure S2.1b**), matching our observation that they are found in inconsistent and disorganized clumps or individually.

The overall interaction matrix (**Figure 2b**) also revealed that there were more inter-type exposures between the anterior cell types of the gastruloid than between the posterior types. This quantification matches the impression from visual inspection that the gastruloids contain both somitic cells and paraxial mesoderm cells that properly localize to the anterior (as per (van den Brink et al., 2020; Yaman and Ramanathan, 2023)) but are arranged in a disorganized fashion. Cells of the posterior, by contrast, were much less mixed, with individual cell types forming large, distinct clusters. These results demonstrate that our analytical methods were able to capture and quantify the degree of spatial disorder in these two compartments. The contrast between organized posterior clustering and disorganized anterior mixing matches expectations based on literature that self-organization mechanisms in gastruloids in the anterior vs. posterior are differentially sensitive to culture conditions, with somitic patterning requiring external matrix support (Beccari et al., 2018; van den Brink et al., 2014; Yamanaka et al., 2023; Yaman and Ramanathan, 2023), distinguishing it from the seemingly more autonomous organization observed in posterior cell types.

Having established the general expectations for cell type interactions, we then looked for deviation from these patterns. We quantified the variance across gastruloids in the exposure index by creating an interaction matrix for each individual gastruloid and comparing the distributions of values for each interaction. We found that for most, the gastruloid-to-gastruloid variation was not significant. However, there were several self-interactions that were significantly variable: specifically presomitic mesoderm, spinal cord, endoderm, and differentiation front (**Figure S2.1c**). This likely reflects differences in the size and shape of clusters of these cell types. There was only one pair of different cell types that significantly varied: endoderm and endothelium. We show additional evidence for the unique arrangement of these cell types in **Figures 5** and **6**.

One potential caveat to this finding is that differences in uncertainty in cell typing could be the primary driver of mixing and cell type interaction differences, both between the anterior and the posterior within an individual gastruloid, or overall between gastruloid. To control for this, we applied an entropy filter to our dataset. We filtered out cells that had entropy > 1.5 (see plot below for cutoff), which was chosen based on the distribution of entropy values for ‘none’ type cells, which effectively describe the upper limit of random transcript assignment (99.7% of ‘none’ type cells are removed with this filter, and about 50% of cardiac mesoderm cells and paraxial mesoderm cells, see **Figure S2.2a**). We first examined overall mixing; there was strong correlation between the per-gastruloid mixing indices before and after entropy filtering (Pearson r = 0.809, **Figure S2.2b**). Globally, mixing indices decreased with filtering, meaning that overall the cell types were more clustered. When we compared the absolute value of the change in mixing index pre and post filtering to the proportion of each cell type, the only significant correlation was with cardiac mesoderm (**Figure S2.2c**). From these analyses we conclude that ambiguous cardiac mesoderm cells drive some of the per-gastruloid mixing index. Exposure indices were overall quite similar after filtering, although the strength of somite-somite and somite-paraxial mesoderm interactions increased (**Figure S2.2d**).

We also examined how entropy filtering might affect the exposure index, given that the mixing index is calculated from the exposure index of across cell types. In general, the magnitude of the exposure index values increased when more uncertain cells were excluded, but the directionality and relative ordering was not affected. Although the magnitude of change in the posterior cells types was less than the anterior cell types, the cross-cell type exposure values, particularly between paraxial mesoderm/endothelium and somites, doubled. From these results we conclude that mixing in the posterior is driven mainly by NMP/presomitic mesoderm interactions, and is overall lower than mixing in the anterior, which is driven by rarer cell types like cardiac mesoderm, paraxial mesoderm, and endothelium, being interspersed within somite cells.

### Micro-scale organization is preserved across gastruloids

Our analyses focused on exposure of one cell type to another, however given that some have reported consistency in gene expression in gastruloids to the resolution of single cells (Merle et al., 2024) we were curious how hyper-local interactions might vary. To measure such interactions, we first determined all the possible combinations of three cells (triplet motifs) from the 9 cell types we had identified (with replacement). Then, for each cell in the gastruloid, we identified the two nearest neighbors and determined which triplet type it was. We calculated the enrichment of each triplet motif relative to its expected frequency based on the amount of each of the constituent cell types. **Figure 2c** shows this distribution of triplet motifs that were found in over half the gastruloids and in each were significantly enriched or depleted based on overall cell type frequency. We found that motifs formed from 3 of the same cell type were overwhelmingly the most significantly enriched, and all were found in over half of the gastruloids. The enriched 2+1 (two same) motifs were composed of all posterior cell types or all anterior cell types. 3/4 of the most common significantly enriched motifs that were of three different cell types consisted of cardiac mesoderm with two other anterior cell types or differentiation front, again suggesting that the cardiac mesoderm cells are uniquely spread out as individuals.

We also identified motifs that were highly underrepresented compared to what would be expected given the frequency of occurrence of each of the individual cell types: nearly all of the significantly underrepresented motifs were of 3 different cell types, with the rarest being somite+NMP+presomitic mesoderm. This observation was initially surprising given that these cells are all thought (in gastruloids) to derive from the same lineage; however, the early mesodermal precursors are spatially segregated from the anterior by the differentiation front, which is where somite determination happens (van den Brink et al., 2020; Veenvliet et al., 2020; Yamanaka et al., 2023). The relative paucity of this triplet indicates that the spatial organization of the differentiation process occurs with very high fidelity, and that less differentiated precursors rarely make it into the anterior, where they would be exposed to somite and paraxial mesoderm cells.

### Variation in cell type abundance and organization is structured and concentrated in specific cell types

We have demonstrated that some aspects of gastruloid composition and spatial organization are consistent across gastruloids, while others are more variable. Consistent features include proportions for NMP, presomitic mesoderm, somite, and paraxial mesoderm, whose coefficients of variation were lower than other cell types (**Figure 1d**). Organizationally, all cell types across gastruloids are more physically clustered than random (**Figure 2a**), and the order in which cell types are found along the AP axis has statistically significant high agreement between gastruloids as measured by Kendall’s W (**Figure S1.5c**). At the local neighborhood scale, we found that most cell type interactions were conserved across gastruloids (**Figure S2.1c**). At the local scale, across individual gastruloids, we found many motifs of three cells that were statistically enriched over random, suggesting a conserved local order (**Figure 2c**). While the normalized distance along the AP-axis of all cell types significantly varied compared to a bootstrapped null (**Figure S1.5a**), the effect size was small, and decreased in almost all cases when normalized to gene expression (of T) in addition to morphology (**Figure S1.5b**).

However, there were also variable features. The proportion of cardiac mesoderm, endoderm, and spinal cord had the highest coefficient of variation between gastruloids (**Figure 1d**). Because proportions must sum to one, a change in the proportion of one cell type is necessarily linked to changes in others; we performed centered log transformation and looked for statistically significant covariation. Of all possible pairings, the following proportions had a significantly negative correlation across samples: endoderm/differentiation front, NMP/endoderm, presomitic mesoderm/endoderm, none/endothelial, and spinal cord/endothelium. This result shows that the proportions of these cell types predictably co-vary between samples, potentially suggesting some kind of biological trade-off in cell type specification or organization (**Figure S1.4d**).

Across gastruloids, intra-cell type interactions (degree of clustering) of spinal cord, endoderm, and differentiation front vary (**Figure S2.1b**). This variation suggests that these cell types may be patterned differently between gastruloids. For example, the local motif of 3 endoderm cells found next to one another was statistically enriched within some but not all individual gastruloids, and by definition is completely absent from gastruloids lacking endoderm (**Figure 2c**). We interpret this contrast to mean that when endoderm is found in a gastruloid, it is consistently patterned at a local level, but may vary more at a global level. This interpretation is concordant with the findings from (Farag et al., 2023), which demonstrate several distinct classes of endoderm organization in gastruloids.

To summarize, while changes in the amount of individual cell types can vary, these changes are in most cases explained by variations in morphology and molecular characteristics (such as anterior:posterior ratio and the expression of morphogens like T). For patterning, we found that, in most cases, global patterns were conserved, but there were variations in local patterning that may lead to variable meso-scale organization of specific cell types, particularly those found in the middle of the anterior-posterior axis.

### Coexpression of cell type marker genes delineates progressive NMP differentiation in gastruloids

One of the central goals of studying developmental processes is to determine the means by which cells make fate decisions and when these decisions are driven by the physical location of the cell as opposed to cell-intrinsic programs. A large part of gastruloid development is thought to be driven by the differentiation, growth, and migration of NMPs and their progeny, which are fated towards either a mesodermal or posterior neural (spinal cord) fate (Bolondi et al., 2024; Braccioli et al., 2022; Veenvliet et al., 2020). In embryos, this process is thought to happen in a continuous fashion; NMP progeny simultaneously move anteriorly and acquire either a mesodermal or spinal cord fate (Garriock et al., 2015; Gouti et al., 2017; Henrique et al., 2015; Jin et al., 2025). The differently fated cells are driven to different parts of the developing spinal column; lineage tracing evidence suggests that fate commitment is at least partially driven by physical location and the local signaling environment (Garriock et al., 2015), and is progressive in nature (Jin et al., 2025). We wondered how this coupling between space and differentiation was resolved in gastruloids, and whether the continuous transition from progenitor to more differentiated cell type was reflected in spatially continuous patterns of gene expression.

Our data represent a single slice of time during which NMPs, presomitic mesoderm, and spinal cord precursors exist simultaneously; if this process happened continuously, we would also expect to see a subset of cells somewhere on an identity continuum between these types (**Figure 3a**). For simplicity, when initially assigning cell types, we used differential expression of panels of marker genes to assign identity. However, many of the cells in the posterior had non-zero cell type scores in all three categories (**Figure S3.1a**). This observation motivated us to leverage the advantages of our dataset to delineate where individual genes were expressed in space with single-cell resolution and determine whether that additional level of resolution gave additional information about NMP differentiation trajectories.

**Figure 3:**
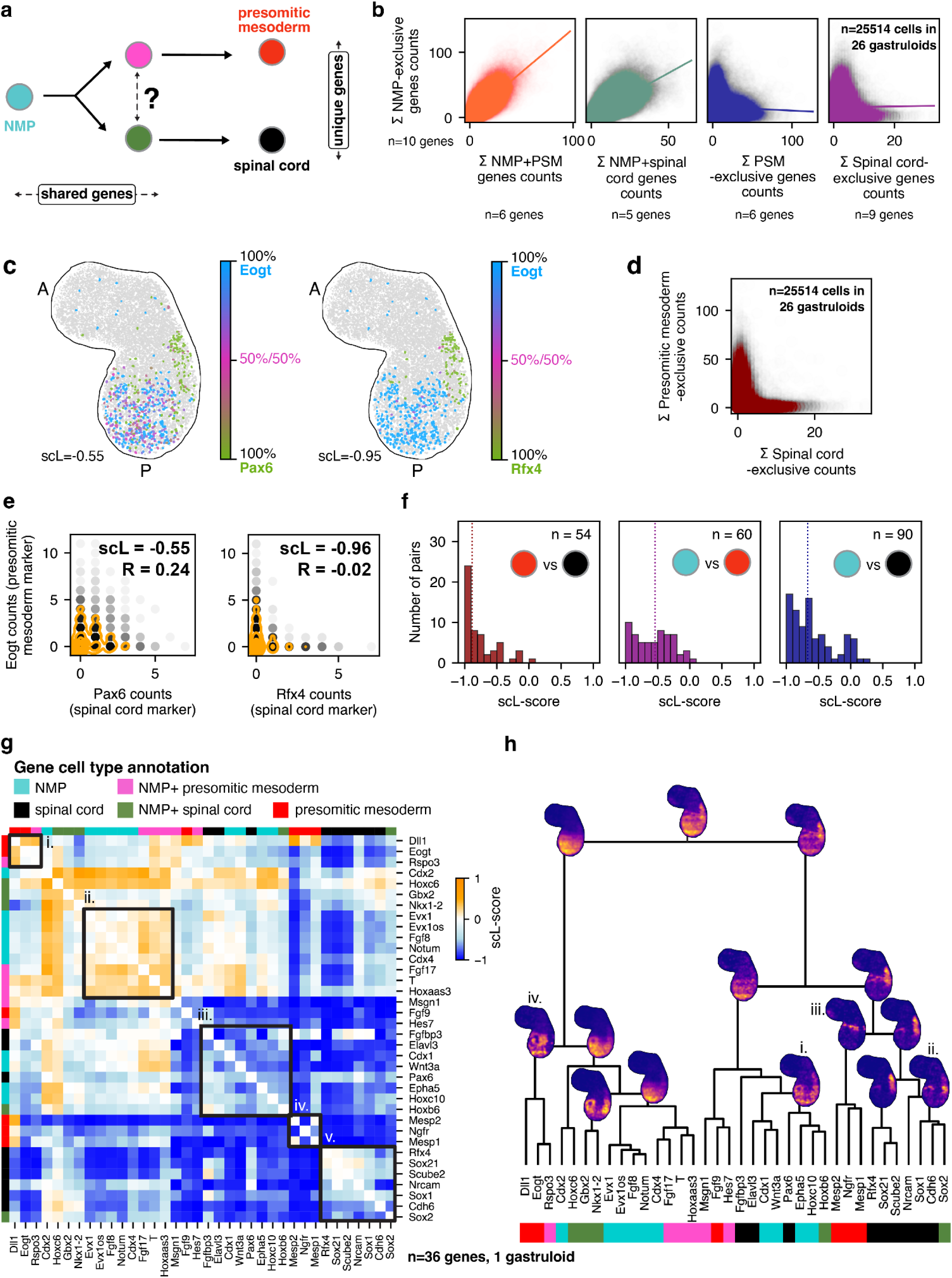
Progressive NMP differentiation is revealed by the single cell L-score. **a.** Illustration of NMP differentiation with predictions for where along the trajectory gene expression is shared or unique. **b.** Per cell expression of all NMP exclusive genes summed versus all other gene categories summed, one plot per type. n=25514 cells typed as NMP, presomitic mesoderm, or spinal cord in n=26 gastruloids. Pearson correlation is shown with colored lines. **c.** Expression plot showing the relative amount of Eogt vs Pax6 (left) and Eogt vs Rfx4 (right) per cell in a representative gastruloid. Pax6 and Rfx4 are both annotated as spinal cord-associated genes. **d.** Per cell expression of all spinal cord exclusive genes summed versus expression of all presomitic mesoderm genes summed. n=25514 cells typed as NMP, presomitic mesoderm, or spinal cord in n=26 gastruloids. **e.** Per cell expression scatterplots of the two pairs of genes shown in b). The y axis of each is the per-cell expression of Eogt. The x-axis is the per-cell expression of Pax6 (left) or Rfx4 (right). R is Pearson’s r, scL is scL-score. Count data is shown in black; smoothed 2D densities are shown in orange. **f.** Distribution of scL-score values for pairs of spinal cord and presomitic mesoderm genes (left), NMP and presomitic mesoderm genes (center), and NMP and spinal cord genes (right). **g.** Hierarchical clustering heatmap of scL-score vectors for NMP, presomitic mesoderm, spinal cord, and combined category genes. scL-score difference vectors were used for clustering; scL-score values are shown in the heatmap. Blocks highlighted are discussed in the text. **h.** Tree of the hierarchical relationships between genes resulting from the clustering shown in e). Color indicating the gene type is shown at the bottom, legend is the same as in **g.** The density plots shown are the summed, averaged gene expression of all genes in the leaves up until that node, smoothed with a 2D density kernel estimate (see Methods for details about smoothing).

Since both presomitic mesoderm and spinal cord cells derive from NMPs, we reasoned that it was possible, especially if the cells were still in the process of differentiating, for cells typed as presomitic mesoderm or spinal cord to residually express NMP genes. Indeed, there were genes in our panel that were shared between each type and the precursor (referred to here as NMP+presomitic mesoderm genes and NMP+spinal cord genes or “shared genes”). **Figure 3b** shows the total expression of each gene group versus the NMP genes for all cells in all gastruloids that we typed as NMP, presomitic mesoderm, or spinal cord. As expected, there was a clear correlation between the mixed categories and NMP genes, supporting the notion of a continuous differentiation process.

In an effort to locate cells that had fully committed to one lineage or the other, we looked at the expression of genes thought to be *exclusively* associated with presomitic mesoderm or spinal cord. Our expectation was that the expression of these genes would be mutually exclusive, and several genes did display this pattern (**Figure 3c**, right). However, we were surprised to find many examples where these supposed marker genes for different cell types were expressed in the same cell (**Figure 3c**, left). Spot assignment has the potential to mis-assign transcripts, which could, in principle, lead to artifactual mixed states. However, we don’t believe misassignment explained our observations. We used stringent nuclear dilation (small enough that ∼30% of transcripts were not assigned to a nucleus). Moreover, when we calculated the exposure index, we found very little exposure between presomitic mesoderm and spinal cord cells (0.08 and 0.05, **Figure 2b**), meaning the majority of the time these cells were not nearest neighbors, and thus had little potential for overlap and transcript mis-assignment.

Was this trend of coexpression true for the two groups of marker genes as a whole? When we compared the total per-cell expression of the genes in both categories (spinal cord and presomitic mesoderm), we noted that while they did show more mutually exclusive (i.e. L-shaped) expression than when compared to the NMP genes, there were clearly many cells that had substantial counts from both PSM and spinal cord gene categories (**Figure 3d**). One possible explanation for the coexpression of genes associated with both arms of the differentiation trajectory is that these cells had not yet fully committed to either lineage. It is also possible that a subset of the genes were better (or at least more exclusive) markers than others (as our observation of the heterogeneity in expression patterns suggested), and that there may in fact be a continuum of markers, some of which may distinguish certain cell types and not others. To determine if any of these possibilities were true, we needed a way to identify genes that were expressed in a mutually exclusive manner.

### The L-score quantifies mutually exclusive gene expression

Marker genes for distinct lineages are often assumed to be mutually exclusive in their expression. However, the above results show that such assumptions may not hold. We were curious whether there were genes that were truly exclusively associated with each cell type, and if such genes in general would prove to be more effective marker genes than those that were merely associated with the cell type.

To this end, we developed a pairwise measure between genes that reported the degree of mutually exclusive expression. It is calculated by rank ordering cells by the expression of one gene and measuring the degree to which the expression of the other gene is anti-rank-ordered (see Methods for details and **Figure S3.2** for a visual explanation of how the measure is calculated). We call this measure the “single-cell L-score” (scL-score) because when the per-cell expression of mutually exclusive genes is plotted against one another, the data make an L shape (**Figure 3e**, right). A value of -1 represents perfectly mutually exclusive expression, which only happens when the genes are never found in the same cell. Higher values indicate more coexpression. Genes that are ubiquitously expressed without a strong correlative relationship between them will have a score of ∼0. The maximum possible value is 1, which is obtained when two genes are expressed only in a shared subset of cells and are never found apart. We refer to this as ‘perfect coexpression’. Our expectation is that ubiquitously expressed genes like cell cycle and housekeeping genes will, due to the rank-ordered nature of the L-score calculation, have L-scores consistently close to 0 no matter which genes they are compared with, whereas genes that are specifically associated with a single cell type will have an scL-score value close to -1 when compared with genes specific to other types, but higher values when compared with genes associated with the same cell type. The results of our simulations confirmed these hypotheses (**Figure S3.3a**).

To benchmark this measure against existing exclusivity or coexpression measures, we calculated the Exclusively Expressed Index (EEI) (Nakajima et al., 2021) and Coefficient of Expression (COEX) (Galfrè et al., 2021) for the same simulated datasets (**Figure S3.3b**) and a subset of NMP/presomitic mesoderm/spinal cord genes (**Figure S3.4a**). All three methods were able, to some extent, to distinguish mutual exclusivity from coexpression, but the scL-score provided clearer separation between these different relationship types; a more detailed description of the analysis is included with **Figure S3.3** and **Figure S3.4**.

We compared the distribution of L-score values between spinal cord and presomitic mesoderm genes; as our previous results suggested, there were many genes that had very strong mutually exclusive expression, while others were expressed in the same cells. For example, Pax6 (spinal cord) and Eogt (presomitic mesoderm) are often expressed in the same cells; they have a mild positive correlation as measured by Pearson’s R (0.24), and an scL-score of -0.55 (**Figure 3e**, left). However, Rfx4 (spinal cord) and Eogt (presomitic mesoderm) were expressed mostly in distinct cells; their expression was nearly uncorrelated when measured by Pearson’s R (-0.02), but they have an scL-score of -0.96 (**Figure 3e**, right). Thus, Rfx4 appears to be a good marker for differentiated spinal cord cells, while Pax6 might mark cells that are neurally fated but not yet committed. We also note that it is not possible to distinguish from these analyses whether all differentiated spinal cord cells express Rfx4, merely that it is exclusively associated with this type.

Overall, the distribution of spinal cord and presomitic mesoderm gene scL-score values was strongly skewed towards -1, with at least 20 pairs of genes strongly distinguishing between the two types (**Figure 3f**, left). From the 54 possible pairs of spinal cord and presomitic mesoderm genes, we found that 24 (44.4%) had scL-score values < -0.9, likely indicating that they mark different populations of cells.

We predicted that there would be more coexpression (i.e., higher scL-score) between NMP genes and the other two categories (presomitic mesoderm and spinal cord), since these two types of cells derive from NMPs. When we compared the distribution of scL-score values of NMP and spinal cord or NMP and presomitic mesoderm gene pairs, we did in fact see a shift of the distribution to higher values compared to the presomitic mesoderm and spinal cord pairs, indicating more coexpression (**Figure 3f**, right).

Despite this shift to higher values, some pairs still seemed to delineate the cell types. For example, Fgf9 (presomitic mesoderm) and Wnt3a (NMP) have a very low scL scores (-0.84). There were even more low-scL pairs when comparing NMP and spinal cord genes, such as Sox21 and Wnt3a (L=-0.98) and Scube2 (spinal cord) and Cdx1 (NMP) (scL=-0.91). Notably, the average scL-score of presomitic mesoderm and NMP gene pairs was much higher than spinal cord and NMP gene pairs, which suggests that perhaps commitment to a neural pathway involves a more systematic reorganization of gene expression.

Analysis using this measure allowed us to distinguish genes such as Eogt and Rfx4 that clearly mark distinct subsets of cells, from those like Eogt and Pax6 that partially overlap and may mark intermediates as well as more differentiated cells. We also observed a unique pattern when we compared the expression of Rfx4, which we identified as a good marker of spinal cord genes, with Nkx1-2, a shared gene annotated as being associated with both NMPs and spinal cord cells. The expression of Nkx1-2 was strongest in the posterior, but extended into the region marked by Rfx4. It seemed to be progressively downregulated in this region from posterior to anterior, producing a graded expression pattern with the anterior-most cells only expressing Rfx4 (**Figure S3.1b, c**). This observation suggests that shared genes can mark cells on their way to becoming spinal cord cells, and at least some are turned off as the cells differentiate, suggesting that the continuous differentiation pathway observed in embryos is also found in gastruloids.

These graded expression patterns led us to be more broadly interested in the patterns of spatial expression and scL-score values of genes that were annotated as belonging to more than one cell type. Up until now, we had excluded such genes from our analyses, but we reasoned that their distribution and coexpression might reveal more about the path that NMPs take on their way to becoming presomitic mesoderm or spinal cord cells. Shared NMP+presomitic mesoderm genes included T, Fgf17, and Hoxaas3, while shared NMP+spinal cord genes included Sox2, Gbx2, and Nkx1-2. When we compared both shared categories to their terminal cell type, we found on average that the distributions of L-score values were positively skewed compared to the NMP and terminal cell type distributions, indicating that, as expected, there was even more coexpression among these categories (**Figure S3.1d**). Interestingly, we found that both distributions were bimodal: a subset of genes were clustered around 0, indicating high coexpression with the terminal cell type genes (this included Dll1 (presomitic mesoderm) and T (NMP+presomitic mesoderm), and Cdh6 (spinal cord) and Nkx1-2 (spinal cord+NMP). Others were closer to -1, meaning the gene in the shared category is either more associated with the parental type (NMPs) or potentially is specific to intermediates. Examples of these pairs include Mesp2 (presomitic mesoderm) and Msgn1 (presomitic mesoderm+NMP), and Rfx4 (spinal cord) and Hoxb6 (spinal cord+NMP).

We wondered whether we could use the scL-scores to refine our marker gene panel. We reasoned that ‘good’ marker genes would have very low scL-scores with most other genes *and* have high scL-scores with a small subset of genes (presumably those belonging to the same type). However, filtering on these criteria did not markedly improve our cell type scoring, likely because the genes in our panel were hand-selected to be good type representatives, so most fit this criterion (**Figure S3.5a, b** and Methods). However, we found that scL-score analysis could be used to quantify how related cell types were in terms of marker gene expression (**Figure S3.5c**), and moreover was extremely effective at assessing the outcome of unsupervised clustering, including quantifying overlap in gene expression between clusters and identifying gene pairs that were mutually exclusively expressed between clusters, even when they were highly related (**Figure S3.6**).

This analysis demonstrates that we could use the scL-score to delineate, in an automated fashion, whether genes reported in the literature as associated with multiple clusters in single-cell data were truly associated with multiple cell types, without parameter tuning and using only the cell-by-gene table.

### The scL-score captures the spatial distribution of gene expression despite being calculated without spatial information

The amount of information about individual gene pairs we were able to extract from this relatively simple measure led us to wonder whether it could also be used to generate a global picture of the relatedness of genes associated with NMPs and their derivatives.

For each gene annotated as belonging to any of the three cell types (NMP, PSM, or spinal cord), we calculated a vector of scL-score values with all other genes. Two genes that play similar regulatory or functional roles would be expected to have similar patterns of coexpression and exclusivity across the full gene panel and thus similar L-score vectors. We reasoned that the Euclidean distance between these vectors could be used for clustering, as it represents the degree to which A and B have a similar scL-score to all other genes considered and satisfies the requirements of a distance measure. We performed hierarchical clustering; a heatmap of the resulting relationships (with the pairwise scL-score values between individual genes displayed for clarity) is shown in **Figure 3g**. Broadly speaking, the genes associated with each cell type clustered together, and the shared category genes generally clustered closely to the NMP genes. Interestingly, although both the NMP and spinal-cord exclusive genes were each found in one large type-specific cluster (**Figure 3g ii., v.**), the presomitic mesoderm exclusive genes were split into two groups (**Figure 3g i., iv**.). There was also a subset of NMP-exclusive and spinal cord-exclusive genes that clustered together and separately from the rest of their group (**Figure 3g iii**.).

To determine what could be driving these differences, we plotted where the genes were expressed in space. Since the clustering was hierarchical, we first made a tree representing this hierarchical relationship, and then at each node of the tree, we made a summed density plot of all the genes contained below that node. The density plots at the top of the tree show broad distributions of expression, but as we followed the branches down closer to the leaves, we saw spatial patterns begin to emerge. We were surprised to find that the scL-score, which doesn’t inherently contain any spatial information, was able to capture a spatial tree of gene expression.

Most of the NMP-exclusive genes were found very close to the posterior and were diffusely expressed. However, in the mixed cluster, which contained both NMP-exclusive and spinal cord-exclusive genes, we saw a different pattern — here the NMP genes were more patchily expressed and extended far more anteriorly (**Figure 3h i**.), suggesting that genes in this group may mark cells destined to become part of the spinal cord. Indeed, one such gene was Epha5, an ephrin receptor differentially expressed in a cluster labeled as NMPs by single-cell sequencing (van den Brink et al., 2020) but which has separately been annotated to be involved in neural development (Akaneya et al., 2010). We also saw a clear division between more anteriorly and more posteriorly expressed spinal cord genes (**Figure 3h ii.**) Thus, clustering L-score values allowed us to delineate differences in behavior between cell type-associated genes: one cluster of spinal cord and NMP genes appears to mark neurally fated but uncommitted cells (**Figure 3h i**.), while another marks committed spinal-cord fated cells (**Figure 3h ii**.).

We also looked more deeply into the split presomitic mesoderm exclusive genes. Not only were they expressed in different parts of the gastruloid (compare **Figure 3h iii., iv**.), we realized that these genes were associated with presomitic mesoderm cells in different states: Mesp2 is turned on in a specific and spatially constrained location at the onset of somitogenesis (Veenvliet et al., 2020), a process which has only just started in this gastruloid, while Dll1 more generally marks the presomitic mesoderm. Thus, using only differences in scL-score values, we were able to identify both general presomitic mesoderm versus presomitic mesoderm that was starting to undergo somitogenesis.

These analyses demonstrate that information contained within the hierarchical relationships between genes, determined by scL-score can reveal novel information about cell states within cell types, although we acknowledge that since cell type is determined by a limited panel of marker genes, results should be further functionally verified. scL-score analysis can also identify distinct spatial locations of cells in this cell state, all without explicit encoding of spatial information, but rather quantifying and clustering the degree to which genes are mutually exclusively expressed with one another.

### Clustering scL-score vectors clearly resolves cell types and reveals novel genetic interactions

Given the amount of spatial and state information that was encoded in the scL-score heatmap for a subset of our gene panel, we expanded our analyses to all genes, hoping to discover new genetic interactions or refine existing ones. We first calculated the scL-score for all genes in all gastruloids, then averaged across gastruloids and clustered the resulting interaction vectors (see Methods for details). The heatmap is shown in **Figure 4a** (heatmap including cell cycle genes is shown in **Figure S4.1a**). We noted that just as when we clustered genes associated with NMPs and their direct descendants, genes associated with cell types tended to cluster together. Specifically, NMP, spinal cord, endoderm, and endothelial genes clustered very strongly together, while presomitic mesoderm genes again were split into two groups, one of which was more closely associated with genes involved in early somitogenesis. We quantified how well cell type specific genes clustered compared to a random null by first calculating the dispersion of cell types within the tree topology using cophenetic distance (see Methods), and then permuting the leaves of the tree to create a null distribution of the dispersion expected by random. The results produced by hierarchical clustering on scL-score vectors were significantly (p=0.0001) more clustered than would be expected by chance (**Figure S4.2a,b**). To assess cluster stability, we randomly selected subsets of the panel and repeated the clustering. Regardless of panel size, the tree produced by clustering on scL-score vectors was always significantly less dispersed than permuted nulls (Figure **S4.2c**). Although our method of calculating dispersion can only be compared between trees clustered on the same gene set, we noted that as we increased the number of genes, the gap between the dispersion of the real tree and the dispersion of the permuted trees increased (**Figure S4.2d**), indicating that, as would be expected, better clustering was achieved when more genes were considered.

**Figure 4:**
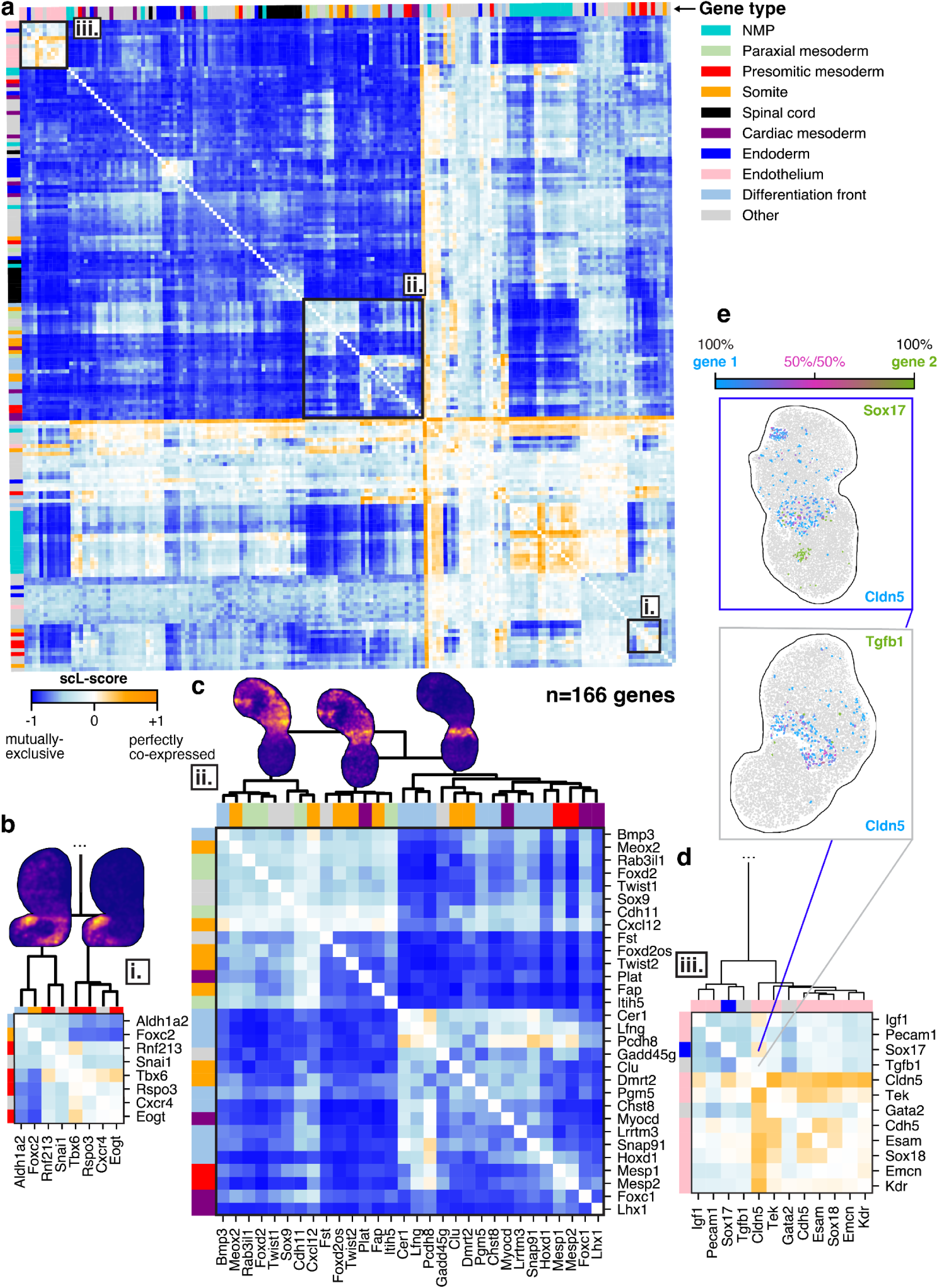
Clustering L-score vectors clearly resolves cell types and reveals novel genetic interactions. **a.** Heatmap of all scL-score values for all genes in the panel (excluding poorly detected and cell cycle genes, see Methods, n=166). Colored bars on the top and right hand sides indicate if a gene is associated with a particular cell type. Heatmap was hierarchically clustered by row. **b.** Expansion of a presomitic mesoderm cluster (**i.**). Clustering relationships are indicated with the dendrogram, and summed densities for all genes in an example gastruloid show where the genes in each cluster are expressed spatially. Clustering relationships determined from the full gene set shown in a. **c.** Expansion of the posterior cell type cluster (**ii.**). Clustering relationships are indicated with the dendrogram, and summed densities for all genes in an example gastruloid show where the genes in each cluster are expressed spatially. Clustering relationships determined from the full gene set shown in a. **d.** Expansion of the endothelial cluster (**iii.**). Clustering relationships are indicated with the dendrogram, and summed densities for all genes in an example gastruloid show where the genes in each cluster are expressed spatially. Clustering relationships determined from the full gene set shown in a. **e.** Expression of two example pairs of genes from the endothelial cluster: Cldn5 and Tgfb1, and Cldn5 and Sox17.

**Figure 5:**
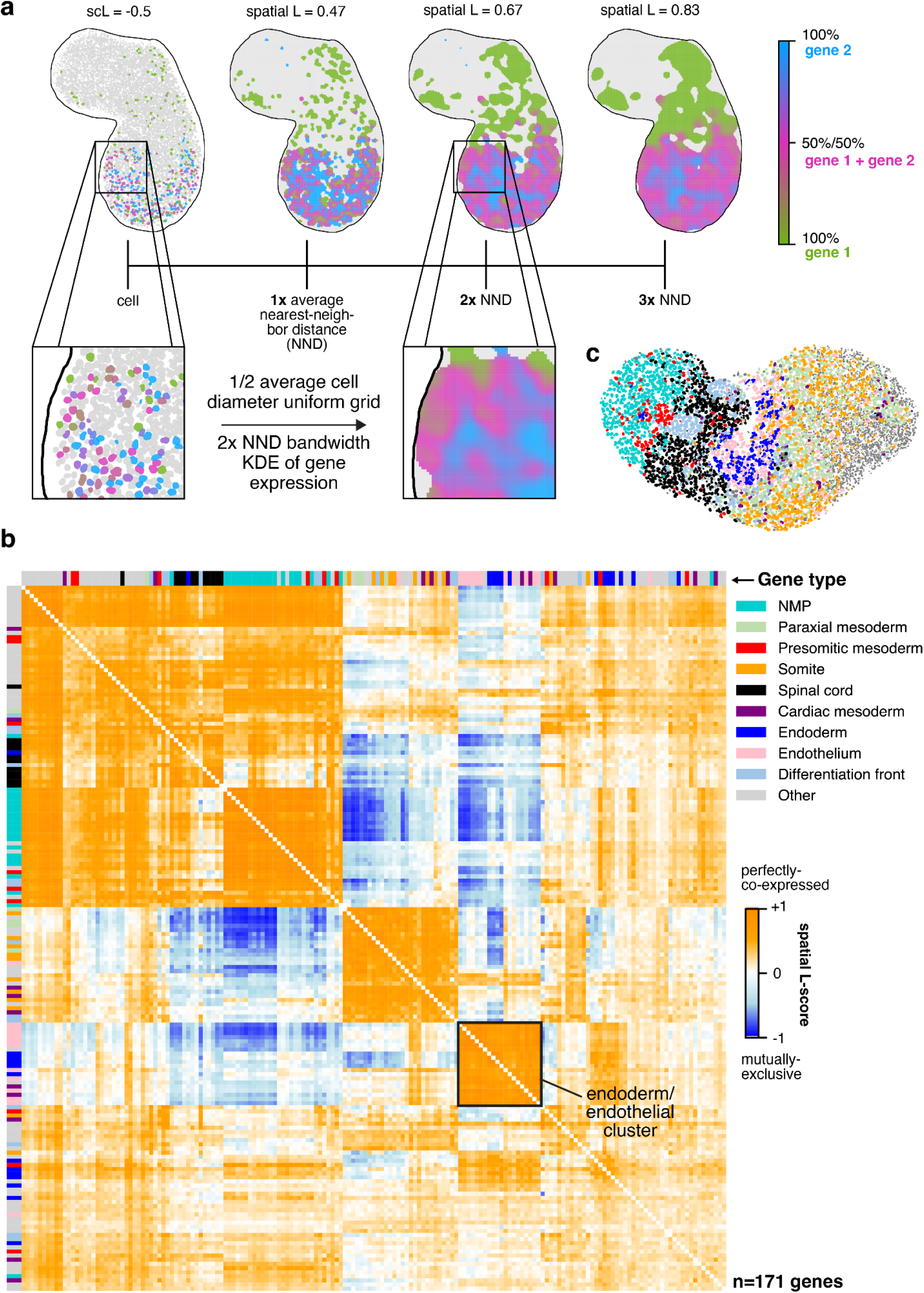
Spatial L-score reveals tissue-level patterns of gene expression. **a.** Illustration of how the density-based L-score (spatial L-score) is calculated, and how the value changes as a function of how much the density estimate is smoothed. **b.** Clustered heatmap of the spatial L-score for a representative gastruloid. Purple box indicates a mixed cluster of endothelial and endoderm genes. **c.** An image of the gastruloid used to generate the heatmap in **b**. Cell types are indicated by color, legend is the same as in **b**.

**Figure 6:**
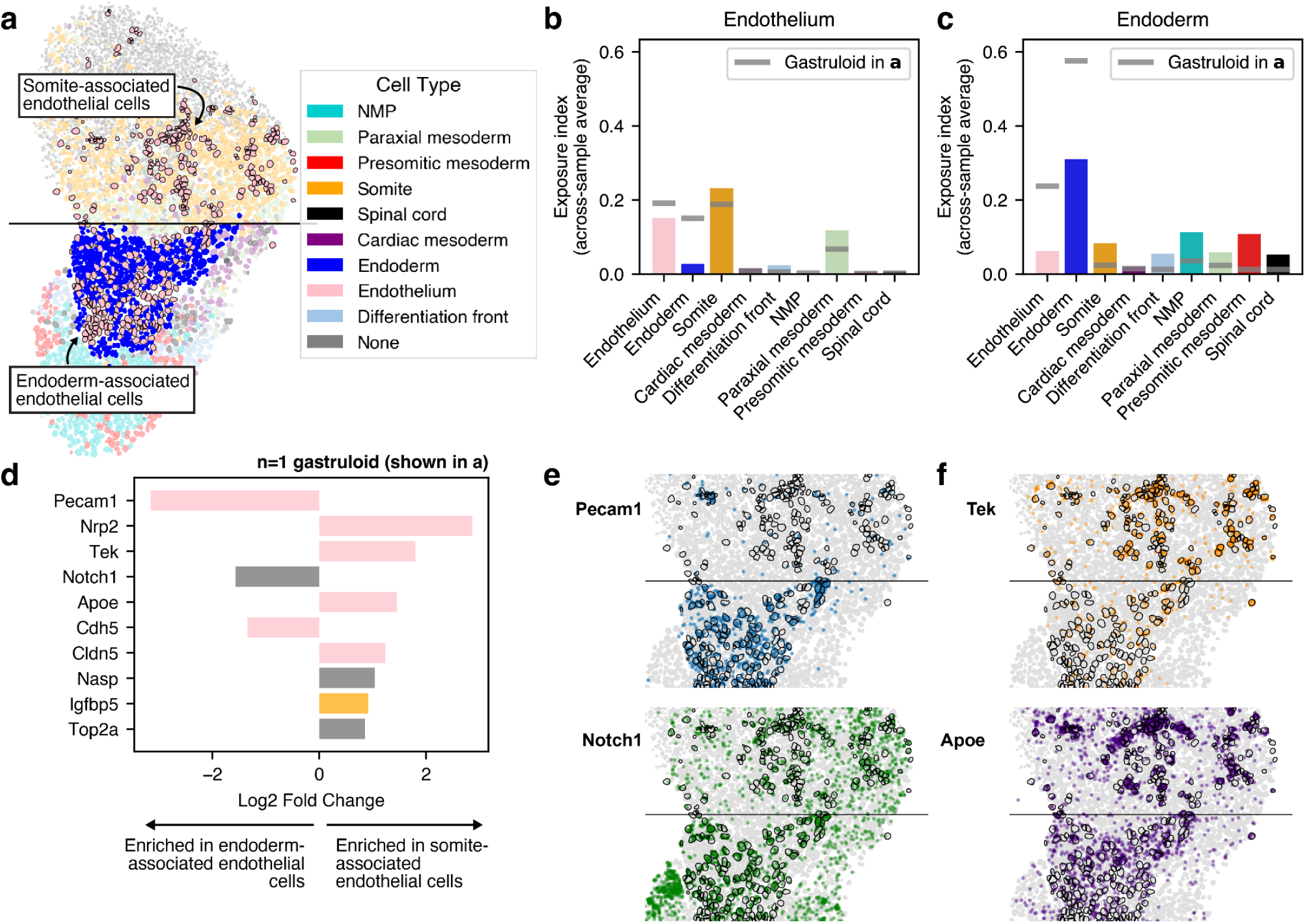
Endothelial precursors show unique organization and distinct, spatially-dependent cell states. **a.** Representative gastruloid showing two distinct morphologies of endothelial cells: somite-associated (anterior) and endoderm-associated (posterior). **b.** Exposure index for endothelial cells compared to all other cell types. Dataset-wide average (n=26 gastruloids) shown in the colored bars, grey lines indicate the values for the gastruloid shown in **a**. **c.** Exposure index for endoderm cells compared to all other cell types. Dataset-wide average (n=26 gastruloids) shown in the colored bars, grey lines indicate the values for the gastruloid shown in **a**. **d.** Genes differentially expressed in somite-associated or endoderm-associated endothelial cells. Bar color represents the cell type associated with the gene (if any); genes that were not previously known to be associated with a cell type in gastruloids are shown in gray. Bars are ordered by significance (adjusted p value) from greatest (Pecam1, adjusted p=8.49e-22) to least (Top2a, adjusted p=7.65e-3). **e.** Spatial distribution of expression for example endoderm-associated genes shown in d). Each plot shows one gene, each dot is a single transcript, and cells typed as endothelial are outlined in black. **f.** Spatial distribution of expression for example somite-associated genes shown in d). Each plot shows one gene, each dot is a single transcript, and cells typed as endothelial are outlined in black.

To generate the heatmap shown in **Figure 4a** and **S4.1a,** we used the same clustering method as described earlier with the Euclidean distance between scL-score vectors. However, we also tried clustering directly on the scL-scores themselves, by transforming each scL-score from the original [-1,1] scale to a [0,1] distance-like scale using a (1-scL)/2 mapping (**Figure S4.3a,b**). The results were largely consistent, however the cophenetic distance scale was relatively compressed when the transformed values were used (**Figure S4.3a,b**). This shallow structure implies that many branches are separated by only modest distances, so fine-scale ordering within the dendrogram should be interpreted more cautiously than the larger-scale cell type block structure. We chose to continue using Euclidean distance-based hierarchical clustering of scL-score profiles, where cell type grouping is observed alongside larger cophenetic separations between clusters.

We examined the largest cluster of presomitic mesoderm genes (**Figure 4a, i.**). Just as before, the clustering reflected distinct spatial distributions; a highlight of one diagonal block with two subclusters is shown in **Figure 4b**, along with the expression patterns in a representative gastruloid. The right subcluster has most of the presomitic mesoderm genes, and is particularly enriched in the left side of this gastruloid, despite the fact that there is a cluster of presomitic mesoderm genes on both sides of the central axis (**Figure S1.3a, xii**). The other subcluster has genes associated with differentiation into somites, and contains Snai1, which is a gene associated with EMT and cells in a more migratory state, consistent with how somitogenesis is thought to occur in gastruloids (van den Brink et al., 2020; Veenvliet et al., 2020; Yamanaka et al., 2023). These genes cluster together because both sets are strongly expressed in a central region of presomitic mesoderm, one is more posteriorly biased, while the other is more anteriorly biased.

The genes associated with more anterior cell types, like somite, paraxial mesoderm, and the differentiation front (which marks the boundary between posterior and anterior), also clustered together in one large block, although the genes associated with the different types tended to be mixed (**Figure 4c**). When we plotted their spatial distributions, the reason for this separation became clear — one block was genes more strongly expressed in the most anterior of the gastruloid, while the other, including almost all of the differentiation front genes, was expressed more posteriorly; these genes were associated with the onset of somitogenesis. We saw similar spatial patterning in the cluster of NMP genes. One sub-cluster, which contained genes like Wnt3a, Hoxc10, and Evx1, was very specifically far-posteriorly localized, while the other, which contained genes like Fgf8, Cdx4, and some of the shared NMP+presomitic mesoderm genes, extended further into the anterior (**Figure S4.1b**).

We also observed interesting differences between the subclusters of endothelial genes (**Figure 4d**). Cldn5 had the strongest average association with other endothelial genes and strong negative associations with genes associated with other cell types, indicating that it is likely the best marker in our panel for generically marking all endothelial precursors. The cluster that contained Cldn5 showed strong intra-cluster associations (high scL-score values), and although the other cluster showed relatively high association with Cldn5, the average association within the cluster and with the genes in the other cluster was weaker. However, this cluster contained some interesting genes: Sox17, an endoderm marker, and Tgfb1 (Tgfβ), which is associated with reduction in inflammation, pro-growth signals, and differentiation (Massagué, 2012). Plotting the expression of these genes in representative gastruloids confirmed their coexpression with Cldn5, a consistent marker for endothelial cells (**Figure 4e**).

Tgfβ appeared to be exclusively associated with Cldn5-expressing cells, but only in some gastruloids (it was extremely lowly expressed in others and not clearly associated with any specific cell type). Tgfβ loss causes embryonic lethality due to hematopoiesis and vascular development failures (Dickson et al., 1995), specifically in extraembryonic tissues. Extraembryonic endoderm has been inconsistently reported in gastruloids; some studies find evidence of this tissue (van den Brink et al., 2020), while others do not (Rosen et al., 2022). Moreover, Tgfβ can drive the transition from endothelial precursors to hematopoietic stem cells (Monteiro et al., 2016), so it is particularly exciting to see it specifically expressed in endothelial tissue. While we note that the regions expressing Tgfβ have unique morphology (**Figure S1.3a, xv.**), further work is needed to determine the role this gene plays in the development of these tissues in gastruloids. These results demonstrate that the scL-score can be used to identify novel cell states in subsets of cell types.

To validate the clustering produced by the scL-score, we compared our results to a state-of-the-art method for identifying gene programs in an unbiased fashion from single-cell data: consensus non-negative matrix factorization (cNMF) (Kotliar et al., 2019). We pooled nuclei from all individual gastruloids and ran cNMF. We found that many of the resulting clusters (**Figure S4.4a,b**) corresponded to the clusters identified when the scL-score tree was truncated to produce exactly the same number of clusters (**Figure S4.4c**). The similarities were even greater when the scL-score clusters were hand-selected based on visual inspection of the tree and density of marker genes (**Figure S4.4d**). To quantify the overlap between clusters, we calculated both the Jaccard Index and the Adjusted Rand Index (ARI) between each scL-score cluster (**Figure S4.4c**) and the most similar cNMF cluster. These distributions are shown in Figure **S4.4e** (blue). We compared to a bootstrapped null where we permuted the genes found in the scL-score clusters, and found that permuted clusters were far less similar to the cNMF clusters than those derived from the real scL-score tree (**Figure S4.4e**). From these observations, we conclude that the two methods are capable of producing similar results at a high-level, but are different in their application. Individual cells receive component scores for cNMF gene programs, yielding more per-cell information, while the tree produced by L-score clustering reveals hierarchical information about gene programs, which quantifies their similarity in expression on a global scale.

Thus, by quantifying the similarity between genes based on the mutual exclusivity of their expression with all other genes, we were able to extract similarity in the expression of cell type genes without explicitly specifying them. Moreover, the spatial relationship between genes, including among genes associated with the same cell type, was encoded in the tree of relationships produced by clustering. Finally, although the strong blocks we find in the heatmaps in **Figures 4a** and **S4.1a** largely reflect cell types, as is consistent with our panel design, we discovered some novel functions of genes in the panel through their location in the scL-score tree: although Tgfβ was initially included in our panel to generally detect inflammatory and growth signaling, clustering by expression patterns revealed its unique association with endothelial precursors. Together, these data demonstrate the power of mutually exclusive expression relationships to uniquely quantify relationships between genes and the cells that express them.

### scL-score analysis reveals cell type groupings and new transcription factor associations in a single-cell RNAseq dataset

To test the generality of scL-score analysis, we analyzed a previously-published dataset from (van den Brink et al., 2020) where individual gastruloids (at the same stage as those used in this study) were pooled and subjected to single-cell RNA-seq analysis. After filtering for cell quality and common gene detection, we calculated the scL-score values using a cell-by-gene table of 14304 cells x 19075 genes.

To first test whether we could reproduce the results from this study, we performed hierarchical clustering on the scL-score difference vector (as previously described) on the set of 207 well-detected genes that were also present in our seqFISH gene panel. The resulting tree showed clustered cell types, similar to the tree produced with the expression data in this study (compare **Figure S4.5a** to **Figure S4.4d**). The cell types were clustered significantly more than expected by chance (**Figure S4.5b**).

Then, to test whether scL-score analysis would be effective for analyzing the entire dataset, we performed hierarchical clustering on all 19075 genes. We then examined the resulting heatmap (**Figure S4.5c**) for clusters of interest. We observed a cluster enriched for endothelial genes (**Figure S4.5d**), which contained some genes in our panel but many others which were not; this finding demonstrates that the clustering in **Figure 4a** is not solely due to the selection of genes in our seqFISH panel. We also observed a large cluster that contained genes associated with pluripotency or primordial germ cell fate (**Figure S4.5e**). Although some of the genes in this cluster were in our seqFISH panel, when we performed scL-score analysis we did not see them cluster with each other or with any other cell type genes. This lack of clustering implies that in our dataset cells that co-express these genes may be rare or too poorly detected to cluster strongly; however, the same analysis performed with more cells and genes showed association. This result demonstrates that clustering scL-score difference vectors can identify known cell-type associated genes, even within transcriptome-scale data. Finally, we also found a small cluster showing strong coexpression of the transcription factor Gata4, a crucial regulator of the development of visceral and parietal endoderm, and two other genes: a predicted gene of unknown function (Gm43715) and Troponin C (Tnnc1) (**Figure S4.5f**). Intriguingly, Gata4 has been implicated in heart development (albeit in an indirect manner) (Watt et al., 2004), and troponin C is important for cardiac muscle cell contraction and has been implicated in cardiomyopathy, although at a much later stage of development than that modeled by gastruloids (Li and Hwang, 2015).

Together, these results demonstrate that scL-score analysis is reproducible across datasets, even when different numbers of genes are compared. It effectively clusters genes associated with cell types, and can reveal developmental transitions. Moreover, increasing the number of cells and genes can reveal new clusters, some of which may predict novel regulatory interactions or spatial co-occurrence not previously observed.

### Calculating the L-score on smoothed gene expression spatial profiles reveals higher-order tissue organization and unique cell type arrangement in certain gastruloids

Our analysis of mutually exclusive gene expression on a per-cell basis both yielded the tools to identify highly specific marker genes and revealed gene expression states that were characteristic of subsets of our annotated cell types. However, there are examples where graded gene expression in tissue, independent of the individual cells expressing the gene, is biologically meaningful: zonation in tissues such as intestine (Moor et al., 2018), liver (Ben-Moshe and Itzkovitz, 2019) and kidney (Lindgren et al., 2017), and hot and cold areas of tumors (Li et al., 2018). Furthermore, many spatial analysis techniques are not resolved to a single cell. Therefore, we wondered whether we could adapt the scL-score to identify patterns of gene expression at different length scales independent of cellular assignment.

To do so, we first noted that in a spatially-resolved dataset, the assignment of spots to nuclei is essentially a non-linear partitioning of genes in space — by being assigned to a nucleus, an individual expression spot loses its specific spatial location and is binned into a region defined by the cell boundary. An example of this binning is shown on the left side of **Figure 5a**. We reasoned that we could instead capture relationships between the purely spatial expression of genes by mapping each transcript onto an evenly spaced grid and then smoothing the resulting 2-dimensional distribution using kernel density estimation. The bandwidth of smoothing can be set in any units, but we found the average (per gastruloid) nearest-neighbor distance to be an intuitive and useful length scale to use. **Figure 5a** shows an example of how choosing different distances for smoothing captures different scales of patterning—any amount of smoothing that causes gene expression in adjacent cells to be averaged would increase the L-score by increasing the degree of coexpression, and more smoothing yields a further increase in L. We chose to use density estimations with a distance value of 2 times the nearest-neighbor average distance, as this spacing seemed likely to capture meso-scale organizational patterns in gastruloids while preserving finer structures of spatial variance (**Figure 5a**).

The scL-score is calculated by comparing gene expression on a cell-by-cell basis. To adapt this measurement to spatial data, we calculated the L-score using the density estimation at each spatial bin across the entire gastruloid (i.e., each bin is treated the way a cell is treated when the scL-score is calculated). We denote this value as the “spatial L-score”.

We calculated the spatial L-score for all the consistently-detected genes in our panel across all gastruloids (202/220 genes, see Methods). When we averaged across gastruloids and clustered the values, the resulting heatmap (**Figure S5.1a**) had many properties similar to the scL-score heatmap shown in **Figure 4a**: For the most part, cell types clustered together; however, we noted a few unique patterns. The order of clustering seemed to more closely reflect overall position in the gastruloid: the more posterior cell types (NMP, presomitic mesoderm, spinal cord) were found close together, then endoderm, differentiation front, and a mixed cluster of paraxial mesoderm, somite, and endothelial genes. This ordering suggests that when the spatially smoothed relationships (spatial L-scores) of all genes in the panel are clustered, the resulting relationships reflect the overall spatial organization of the underlying cell types within the larger tissue structure. Individual cell types appeared to cluster more strongly when using scL-score, but the order and mixing of cell type genes with the spatial L-score reflects tissue-level organization—NMP and presomitic mesoderm genes were close together and mixed, reflecting the mixing we see of the cell types associated with these genes.

The other difference of note was that, unlike in the scL-score heatmap, the endothelial genes were much less strongly clustered, likely reflecting the diversity of locations they were found in within individual gastruloids (**Figure S1.2**). We looked at their patterns in the heatmaps for individual gastruloids and noticed that there was strong co-clustering of endothelial and endoderm genes in some of the gastruloids. This was hinted at earlier when we calculated significant differences in the interactions between these cell types between gastruloids (**Figure S2.1c**). An example is shown in **Figure 5b**, which is the clustered heatmap for an individual gastruloid (cell types and morphology shown in **Figure 5c**). A strong block of interaction on the diagonal (“endoderm/endothelium cluster”, outlined) indicates a strong association of these genes on a length scale beyond individual cells. When we looked at the cell type arrangement of this gastruloid, there were two clear populations of endothelial cells: those that were located in the anterior and those that were more posteriorly located and strongly intermixed with the endoderm tissue. Thus, the unique clustering behavior of genes in this gastruloid is reflective of a unique, tissue-level pattern that is only found in some of our samples. When we calculate the scL-score heatmap for the same gastruloid, although endothelial and endoderm genes cluster close together, they form two distinct diagonal clusters, compared to the single, high-L block found in the spatial-L heatmap (compare the outlined clusters in **Figure 5b** and **Figure S5.2a**).

These comparisons suggested that the spatial L-score could capture patterns on a tunable length scale. When we directly compared the intra-cell type L-score values calculated using both methods (**Figure S5.1b**), we saw that while the values for some cell types, like NMP, were quite similar, others, such as spinal cord, showed a dramatic shift towards +1 when calculated using a smoothing of 2 nearest neighbors. This shift is consistent with our observation of consistent expression of NMP genes and a more “salt-and-pepper” expression of spinal cord genes, and suggests that by calculating the spatial L-score over many length scales, one could learn the effective length scale of interaction for specific gene pairs.

### Endothelial precursors show unique organization and distinct, spatially-dependent cell states

We observed that in 5 out of the 26 gastruloids, there was a large central patch of endoderm cells intermixed with endothelial precursors; these samples also had unique spatial L-score clustering of endothelial and endoderm genes (**Figure 5b**). An example of one such gastruloid is shown in **Figure 6a**. Migration to and association with the endoderm is also a hallmark of endothelial development (Dyer and Patterson, 2010; Sendra et al., 2024), and we were curious whether there were differences between these cells and the cells we observed forming anterior, somite-associated clusters. When we computed the cell type exposure index for just this gastruloid, we found that, consistent with our visual observations, in this particular sample endothelial and endoderm cells were much more frequently found next to one another than on average (**Figure 6b,c**). To determine whether these spatial and organizational differences reflected gene expression differences, we divided the gastruloid normal to the anterior-posterior axis to separate the endothelial cells into endoderm-associated and somite-associated and looked for differentially expressed genes between the two groups in this gastruloid. To ensure we were focused on genes that truly varied in expression in endothelial cells and were not merely a reflection of spillover from surrounding cells, we pre-filtered genes on expression, so only genes that were present in at least 50% of the cells in either group at a greater than 2 count per cell level were considered. The significantly differentially expressed genes after filtering are shown in **Figure 6d**. As an additional check on the degree to which transcript mis-assignment affected our analysis of gene expression in these cells in particular, we varied the nuclear dilation in this gastruloid specifically, and calculated cell type score entropy as a function of nuclear dilation (**Figure S6.1a**). Because cell type score entropy of a cell reflects the degree to which that cell specificity expresses genes associated with a single cell type, our expectation was that if spillover between endoderm and endothelial cells was a significant issue, then decreasing the nuclear dilation should greatly decrease the entropy scores for both groups. Although we saw a slight increase in the spread of the distribution as nuclear dilation increased, the median cell type entropy stayed extremely low for both groups (**Figure S6.1a**). From this analysis we conclude that the genes we identify as differentially expressed are not due to spillover from surrounding cells, but instead are due to spatially-dependent differences in endothelial cell biology.

The genes with the highest fold-change in expression in endoderm-associated endothelial genes are shown on the left hand side of **Figure 6d**. Two are endothelial genes: Pecam1 and Cdh5, both of which are associated with angiogenesis. Spatial expression of these genes is shown in **Figure 6e** (larger version in **Figure S6.1b)**. Notch1 is more expressed in endoderm-associated endothelial cells, and this could reflect an increase in Notch signaling in the posterior of the gastruloid. (Chan et al., 2017) demonstrated that Notch signaling can be sensitive to shear stress, raising the possibility that the differences in cell state we observe may be driven by differences in mechanical forces in the anterior and posterior. Although most endothelial cells are thought to be of mesodermal origin, some evidence suggests that, in the organogenesis of specific tissues like the liver, the endoderm can give rise to endothelial cells (Goldman et al., 2014). Furthermore, in (Rossi et al., 2022) the authors show that in a gastruloid-like model specifically designed to model blood development, there is strong spatial adjacency between endothelial and endoderm cells. They hypothesize that these may be a subset of endothelial cells, specifically hemogenic endothelial cells (which have the potential to become blood progenitors). Our data demonstrate a molecularly driven organization distinct from the clustering we observed in the anterior and suggest that multiple mechanisms of endothelial specification could be modeled in gastruloids, even simultaneously within the same structure, although further characterization is needed to determine exactly what processes these unique endodermal/endothelial structures model.

Several other endothelial genes are instead differentially expressed in somite-associated endothelial cells: Nrp2, Tek, Apoe, and Cldn5. Spatial distributions of these genes are shown in **Figure 6f** (larger version in **Figure S6.1b**). Although these genes have less obvious functional distinctions than the endoderm-associated genes, Nrp2 enables semaphorin receptor activity, including nervous system development and ventral trunk neural crest cell migration and Tek negatively regulates endothelial cell apoptotic process and response to retinoic acid (RA), which is known to be higher in the gastruloid anterior. Furthermore, a specialized population of endothelial precursors associated with somites was also observed in trunk-like structures, which show more tissue-like organization than gastruloids (Veenvliet et al., 2020).

Although endothelial cells have consistently been observed in single-cell measurements of gastruloids, their relative rarity has precluded in-depth analysis of subtypes or inference of spatial location. Our results strongly suggest that endothelial precursor formation, migration, and organization may all be modeled in 3D gastruloids, even without treatment with additional factors as in (Rossi et al., 2022, 2021); recent advances in 2D gastruloids have allowed modeling of cardiac and hepatic vascularization (Abilez et al., 2025), and our data suggest that 3D gastruloids may similarly be adapted to model more specific aspects of hematopoiesis and vascularization. Early specification from a pool of mesodermal precursors is a hallmark of the endothelial lineage (Sendra et al., 2024); given the consistency with which we observe endothelial precursors, we speculate that this behavior is recapitulated in gastruloids, but further epigenetic measurements are required to validate this hypothesis.

## Discussion

A central goal of developmental biology is to map how the identity and arrangement of cells are determined and coordinated. Although precise scaling of gene expression in gastruloids had been previously reported (Merle et al., 2024), the relationship between this scaling and the proportions of specific cell types or more complex tissue organization was not known. Towards that end, we have performed detailed measurements and analyses of the expression of hundreds of genes in tens of individual gastruloids, revealing how cell types are organized within and across gastruloids. Our results highlight that many aspects of embryonic development, such as the arrangement and differentiation trajectories of posterior cell types, are faithfully recreated on a per-gastruloid basis. Others, however, can vary, and conditions may need to be tuned to increase reproducibility if a specific aspect of their development needs to be modeled. For example, we reproduced observed variation in endodermal structures (Farag et al., 2023) and reported both the distinct organization of endothelial cells and variation in that organization between gastruloids. Although we do not fully reproduce the level of per-gastruloid cell type proportion variation reported in a previous single-cell sequencing dataset (Rosen et al., 2022), we do find evidence of a tradeoff between neural and mesoderm tissues; differences in the number of cells analyzed per gastruloid could potentially explain this discrepancy. Moreover, we observe many of the same cell types as in this study, and show that their organization is quite consistent.

We introduce the L-score, which quantifies mutually exclusive gene expression, and highlight two ways of using it that can be employed on different types of data. The cell-based L-score, scL-score, can be applied to any technique that generates a cell by gene table. Hierarchical clustering analysis of the resulting vectors can reveal how similar each gene is to others in terms of how it is co-expressed with all genes, and may reveal spatial information if the sample was collected from a spatially-organized tissue. We were able to verify the spatial information because our data were spatial, but the scL-score is calculated only using the cell-by-gene table, a standard output for any single-cell analysis technique.

The spatial L-score, which operates spatial bin-wise rather than cell-wise, can be used on spatial data even when cell assignment or nuclear segmentation isn’t possible — in principle, it can reveal which genes are patterned similarly at many different length scales, with the fundamental resolution limit set by the measurement technique. It was able to identify coexpression, even amongst lowly-expressed genes, such as the spinal cord markers, that had a more ‘salt-and-pepper’ expression pattern and were only strong markers when considered in aggregate. It is possible that some genes with bursty gene expression may appear not to express together, but spatial smoothing may allow for the identification of coexpression of these genes, which may be useful for cell type identification.

Although the identification of cell types has utility for high-level analyses of organization, our in-depth dissection using the L-score revealed that, particularly for cells in the process of differentiating, a gene-centric view may yield a more holistic picture of tissue organization. It is particularly exciting to imagine these analyses performed in a time-resolved fashion, as dynamic changes in gene expression across a tissue in response to physical rearrangement and broad signaling patterns could reveal mechanisms or spatial aspects of differentiation not visible when cells are merely binned into predetermined categories.

---

**Figure S1.1:**
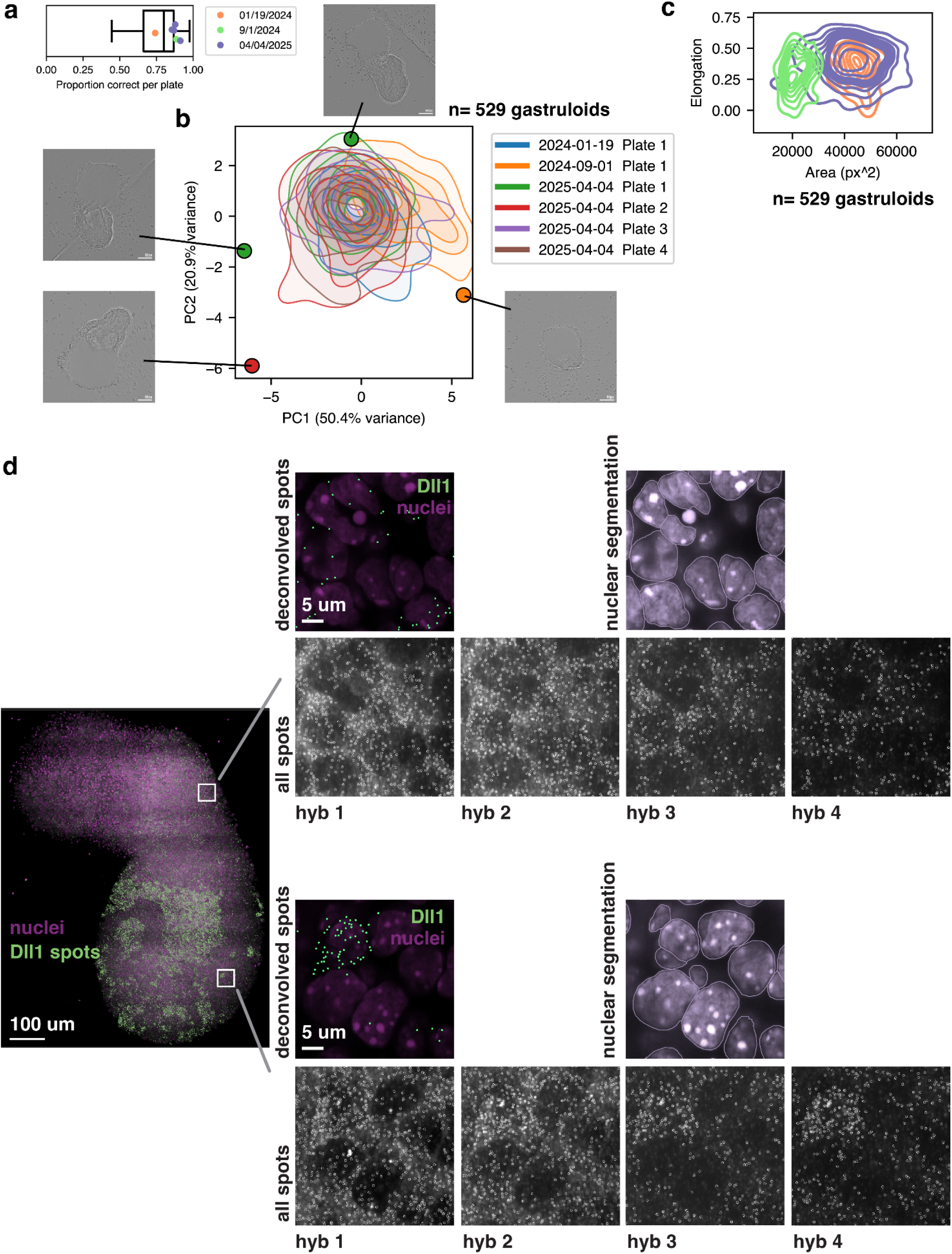
Gastruloid shape analysis and representative raw seqFISH images, spot detection, and gene deconvolution. **a.** Proportion of each plate of gastruloids that, at 120 hours, were scored as ‘correct’. The correct phenotype was elongated with one axis and a clear anterior and posterior domain. The experiments used to generate the samples used in this paper are shown with colored dots, with colors indicating the day. One of the datasets used multiple plates pooled together, but the plates are shown individually for clarity. **b.** PCA projection of the morphological characteristics of all gastruloids generated in the 3 separate experiments analyzed in this study. n = 529 total gastruloids, 84-90 gastruloids per plate. Highlighted points are the extreme values in each dimension; images of the relevant gastruloid are shown in insets. **c.** Area and elongation of all gastruloids generated for this study (n=529), colored by date. **d.** Representative images showing nuclear staining, spot detection by hybridization (hyb) and deconvolution of Dll1.

**Figure S1.2:**
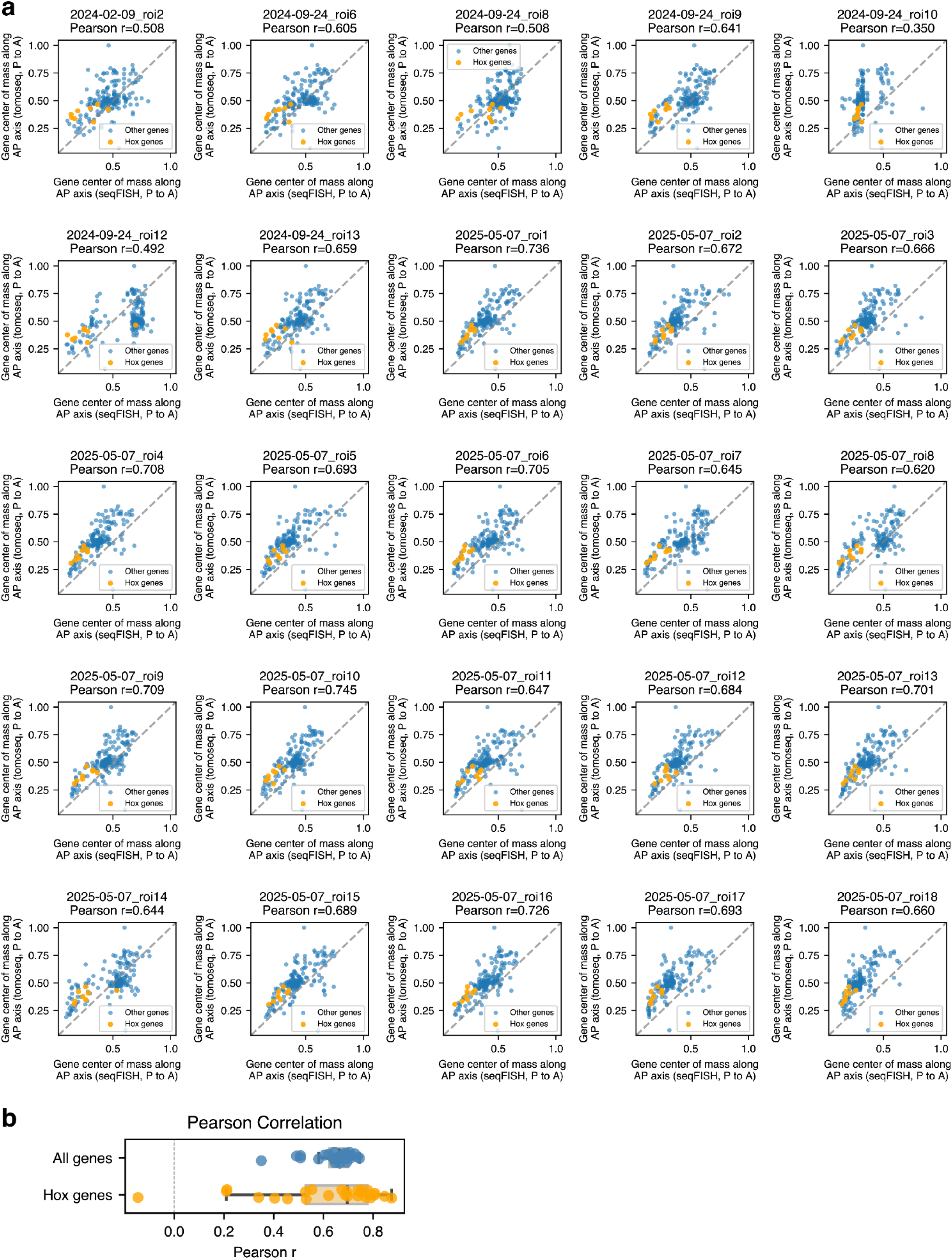
Comparison of AP-axis gene expression centers of mass between individual gastruloids reported in this work and previously-reported tomoSeq performed on gastruloids. **a.** Center of mass of gene expression in all gastruloids compared to the center of mass of genes expression in one gastruloid analyzed by tomo-seq as reported in (van den Brink et al., 2020). The number of genes in each plot is shown in the title; numbers vary by gastruloid as some of the genes in the seqFISH panel were not consistently detected across gastruloids (see Methods). Orange points are the Hox genes included in the panel. **b.** Summary of the Pearson correlation of all samples shown in a) for all genes (blue) and Hox genes in the seqFISH panel (orange).

**Figure S1.3:**
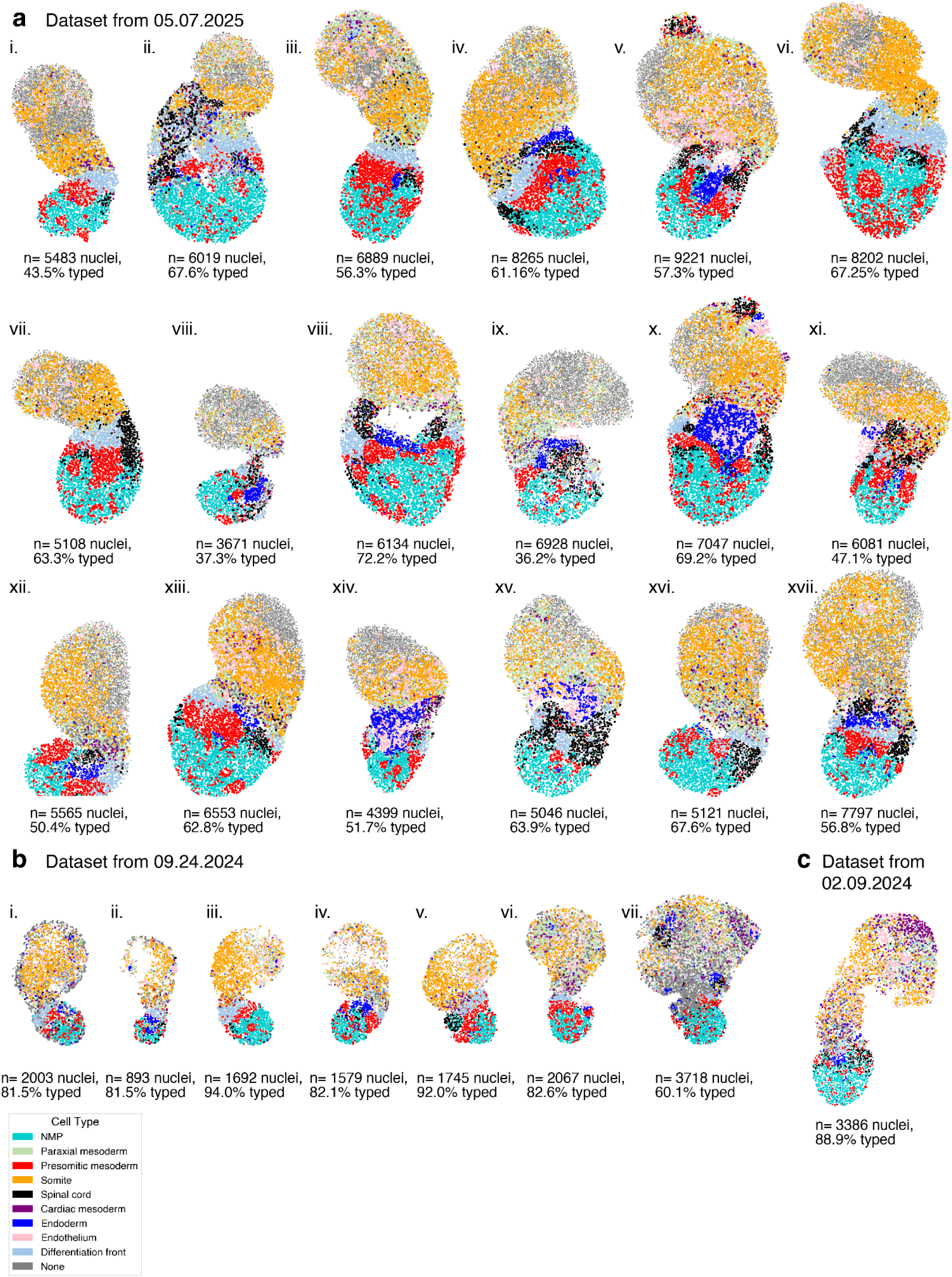
Gallery of individual gastruloids colored by cell type. **a.** Gastruloids from the experiment conducted on 05/07/2025; color of each nucleus indicates cell type (legend on bottom left). **b.** Gastruloids from the experiment conducted on 09/24/2024; color of each nucleus indicates cell type (legend on bottom left). **c.** Gastruloids from the experiment conducted on 02/09/2024; color of each nucleus indicates cell type (legend on bottom left).

**Figure S1.4:**
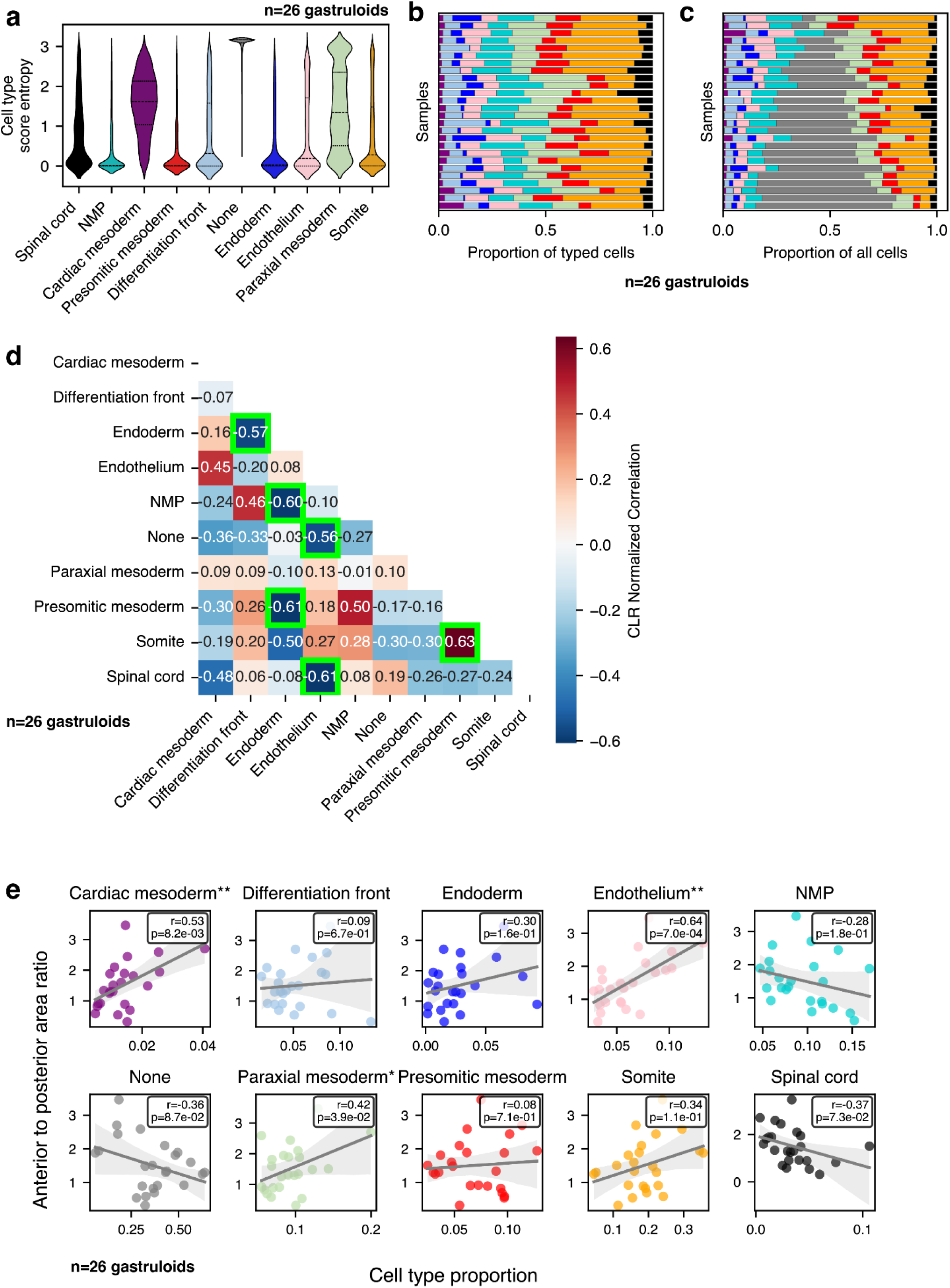
Cell type entropy scores, cell type proportion covariation, and correlation between individual cell type proportions and morphological characteristics. **a.** Cell type score entropy distributions for each cell type. Each violin shows all the values for that cell type across all gastruloids. n=79607 nuclei across 26 gastruloids. **b.** Per-gastruloid cell type proportions (summarized in Main Figure b). **c.** Per-gastruloid cell type proportions including untyped cells (’None’). **d.** Co-variation in proportion (normalized by Centered Log-Ratio (CLR) transformation) between cell types. Green boxes indicate significant hits (adjusted p value < 0.05 at a 5% FDR). **e.** Correlation between the per-gastruloid mixing index with the proportion of the labeled cell type in that gastruloid. Pearson’s r and the significance (p) for each relationship is shown in the black box. ** = p < 0.01, * = p < 0.05.

**Figure S1.5:**
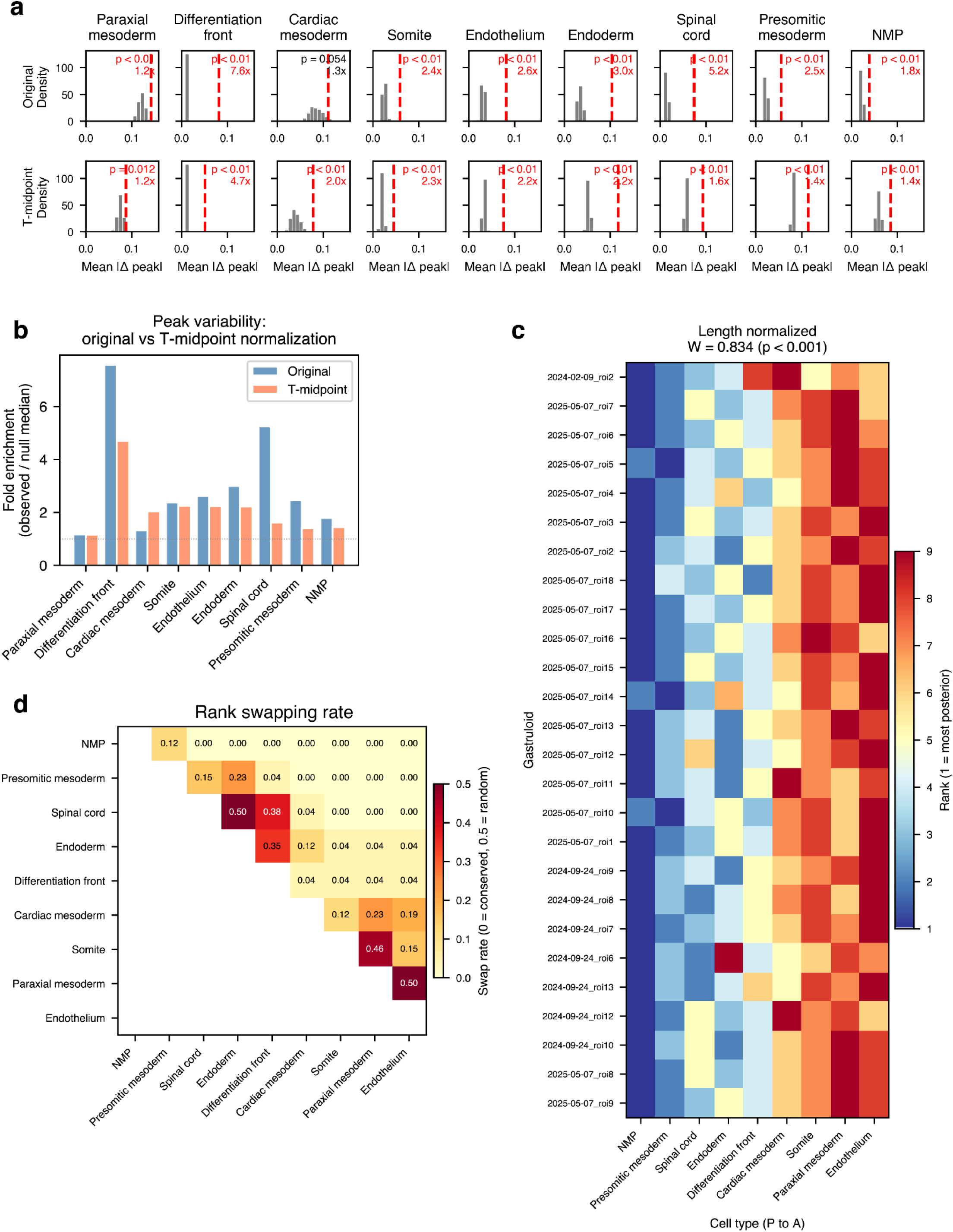
Analysis of the AP-axis location and rank order of cell types. **a.** Bootstrapped null distributions for the difference in peak location to the overall mean for each cell type along the AP axis length normalized (top row) or normalized to length with the midpoint determined by T expression (bottom row). Red lines show the average value for the observed peak differences in the 26 samples shown in Figure S1.3a,b,c. The fold change from the mean is shown in the inset, as is the p value for the difference. **b.** Comparison between the fold changes reported in the top and bottom row of a). **c.** Rank heatmap for the peak location of each cell type for each gastruloid (length normalized). **d.** Rate of rank swapping between cell types; 0=conserved, 0.5=equal likelihood of occurring at two adjacent ranks.

**Figure S2.1:**
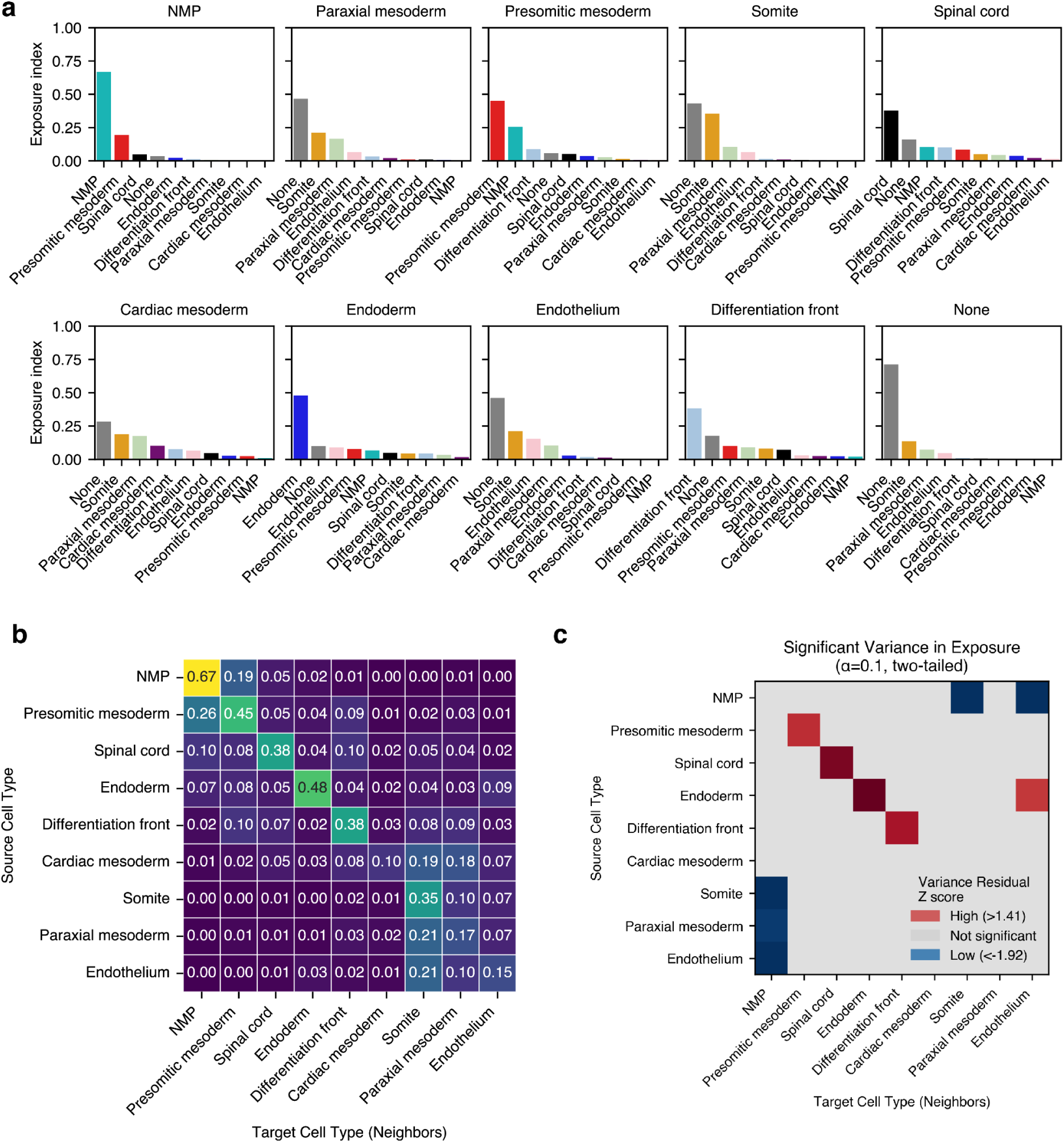
Exposure indices triplet motifs. **a.** Exposure indices for all cell types shown individually as bar plots. **b.** Significantly varying (red) and conserved (blue) neighbor interactions across gastruloids (n=26). **c.** Exposure index matrix with self-interactions included.

**Figure S2.2:**
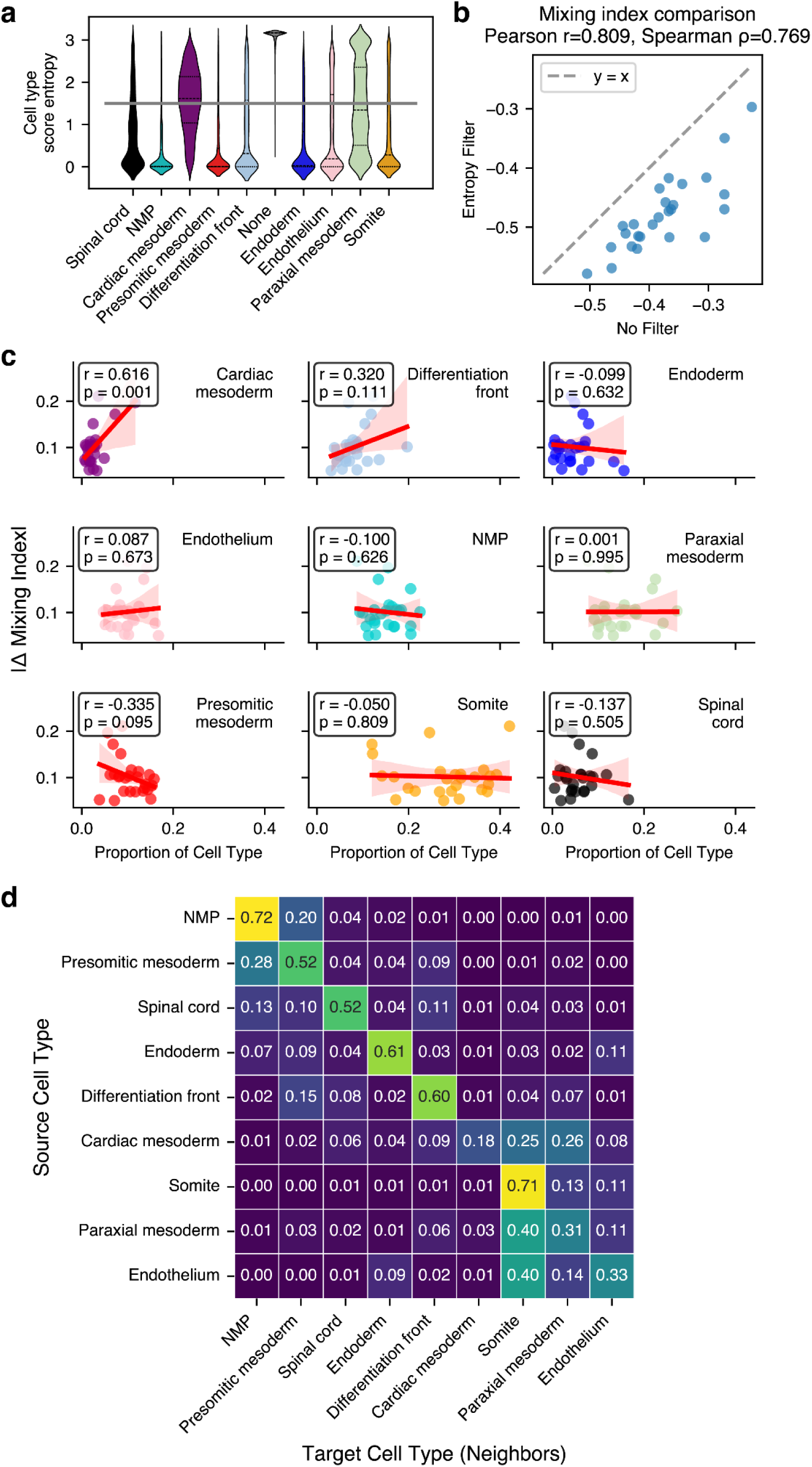
Cell type scoring robustness. **a.** Entropy cutoff used to generate a subset of high-confidence cells. At this cutoff (entropy <= 1.5) 99.7% of ‘none’ type cells are removed, as well as ∼50% of cardiac mesoderm and paraxial mesoderm cells. **b.** Correlation between the overall mixing index per-gastruloid, calculated before and after entropy filtering (n=26 gastruloids). **c.** Correlation between the proportion of each cell type and the change in mixing index before and after entropy filtering. Pearson’s r and the associated p value are shown in the inset. **d.** Full exposure index matrix recalculated after entropy filtering.

**Figure S3.1:**
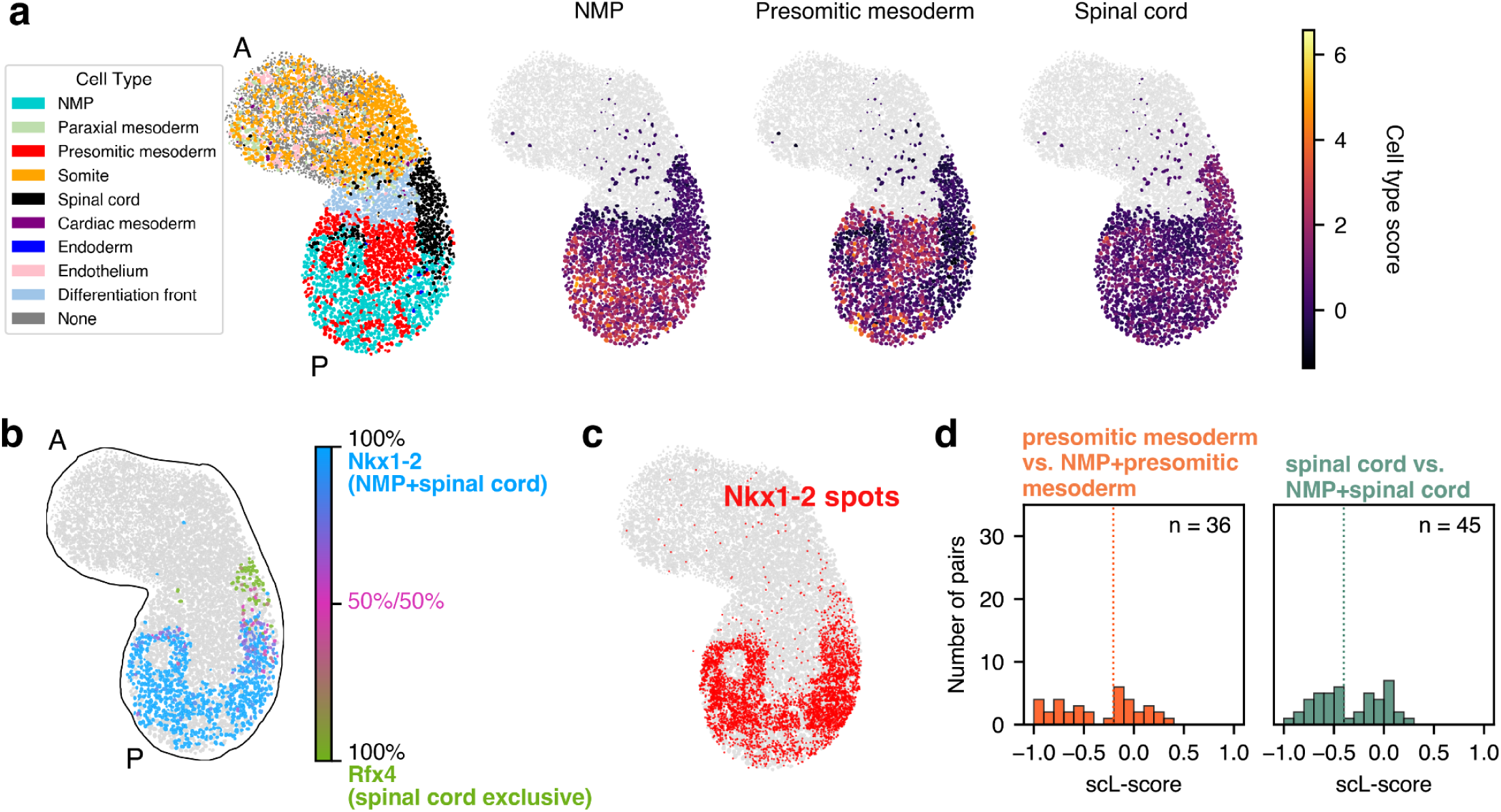
Single cell L-score of NMP, presomitic mesoderm, and spinal cord genes. **a.** Left: cell type scores for all nuclei in the gastruloid used in this figure. Right: cell type scores for all nuclei typed as NMP, presomitic mesoderm, or spinal cord in each of those categories. **b.** Spatial distribution of the expression of Nkx1-2 and Rfx4 in an example gastruloid. **c.** Nkx1-2 transcripts in the same gastruloid as in **c.** **d.** Distribution of scL-score values between pairs of terminal cell type genes and genes annotated as belonging to that cell type and NMP genes.

**Figure S3.2:**
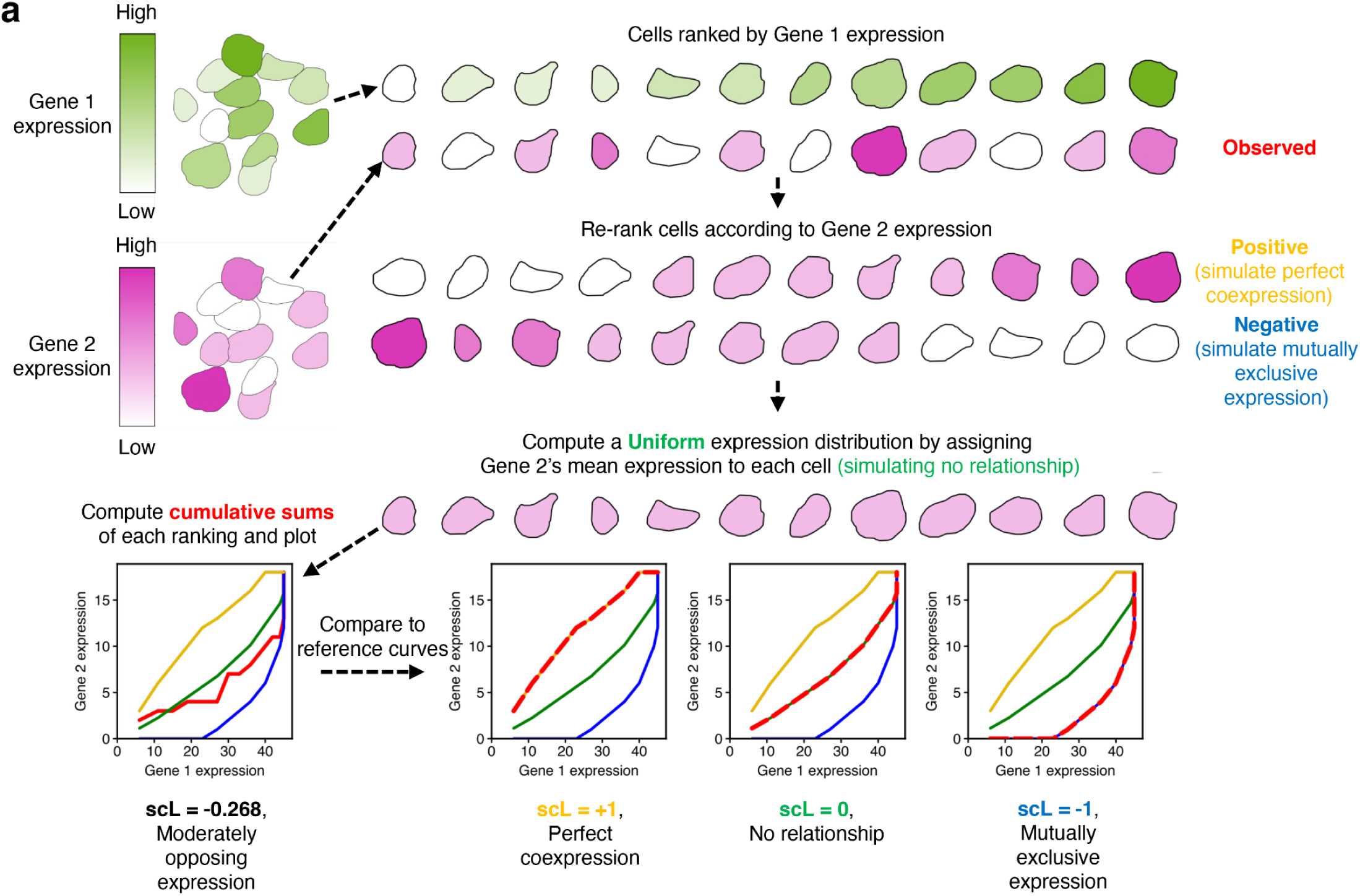
Illustration of L-score calculation. **a.** Illustration of the calculation of the scL-score.

**Figure S3.3:**
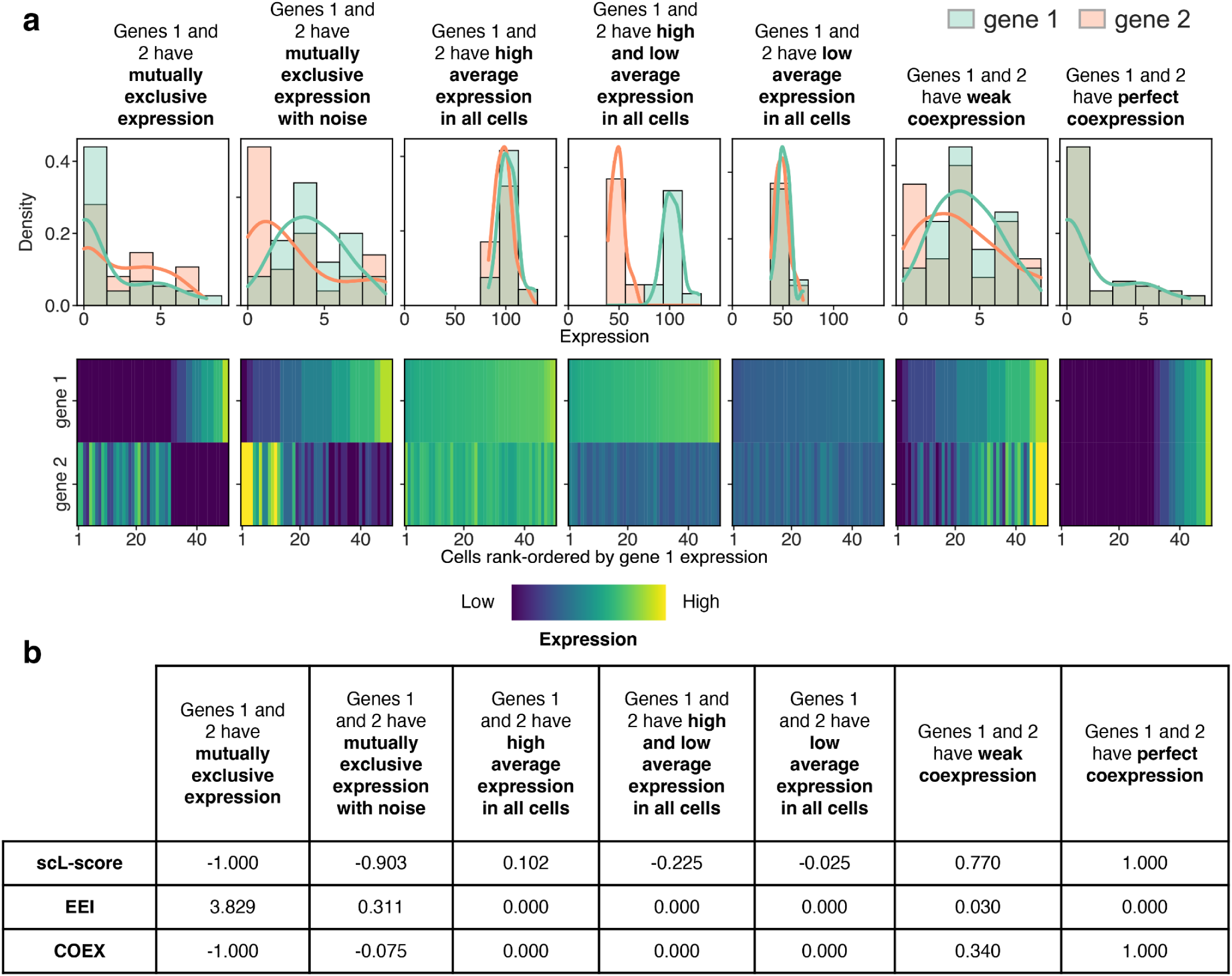
Comparison between scL-score, Exclusive Expression Index (EEI), and coefficient of coexpression (COEX) calculated from simulated data representing common expression scenarios. **a.** Simulations of several common gene expression scenarios. The top row shows the simulated per-cell expression distributions for each scenario, and the bottom row shows the expression values rank-ordered by the expression of gene 1. This rank-ordering is used to calculate the L-score. **b.** scL-score, EEI, and COEX (from the COexpression Table ANalysis (COTAN) framework) for the scenarios shown in **a**. We compared the scL-score to two existing measures of exclusivity. The Exclusively Expressed Index (EEI) (Nakajima et al., 2021) is bounded below by 0 and increases with mutual exclusivity. EEI is computed from binary zero/non-zero quantification and is designed specifically to measure exclusivity but not coexpression. The coefficient of coexpression (COEX) from the COexpression Table ANalysis (COTAN) framework (Galfrè et al., 2021) can also be used to quantify relationships between genes: positive values indicate coexpression, negative values indicate mutual exclusivity, and values near 0 indicate little structured relationship. In the two mutually exclusive simulations, all three methods detected exclusivity. In the three simulations of genes independently expressed in all cells (but with varying relative expression levels), EEI and COEX were 0, while the scL-score remained close to 0 (0.102, -0.225, and -0.025, respectively), consistent with little structured relationship. In the weak coexpression simulation, the scL-score was positive (0.770), EEI remained close to 0, and COEX was positive (0.340), indicating detectable but modest coexpression. In the perfect coexpression simulation, the scL-score reached 1.000, EEI was 0, and COEX was strongly positive (1.000). Together, these simulations show that all three methods detect strong mutual exclusivity, and both scL-score and COEX distinguish positive coexpression from exclusivity and from unstructured expression.

**Figure S3.4:**
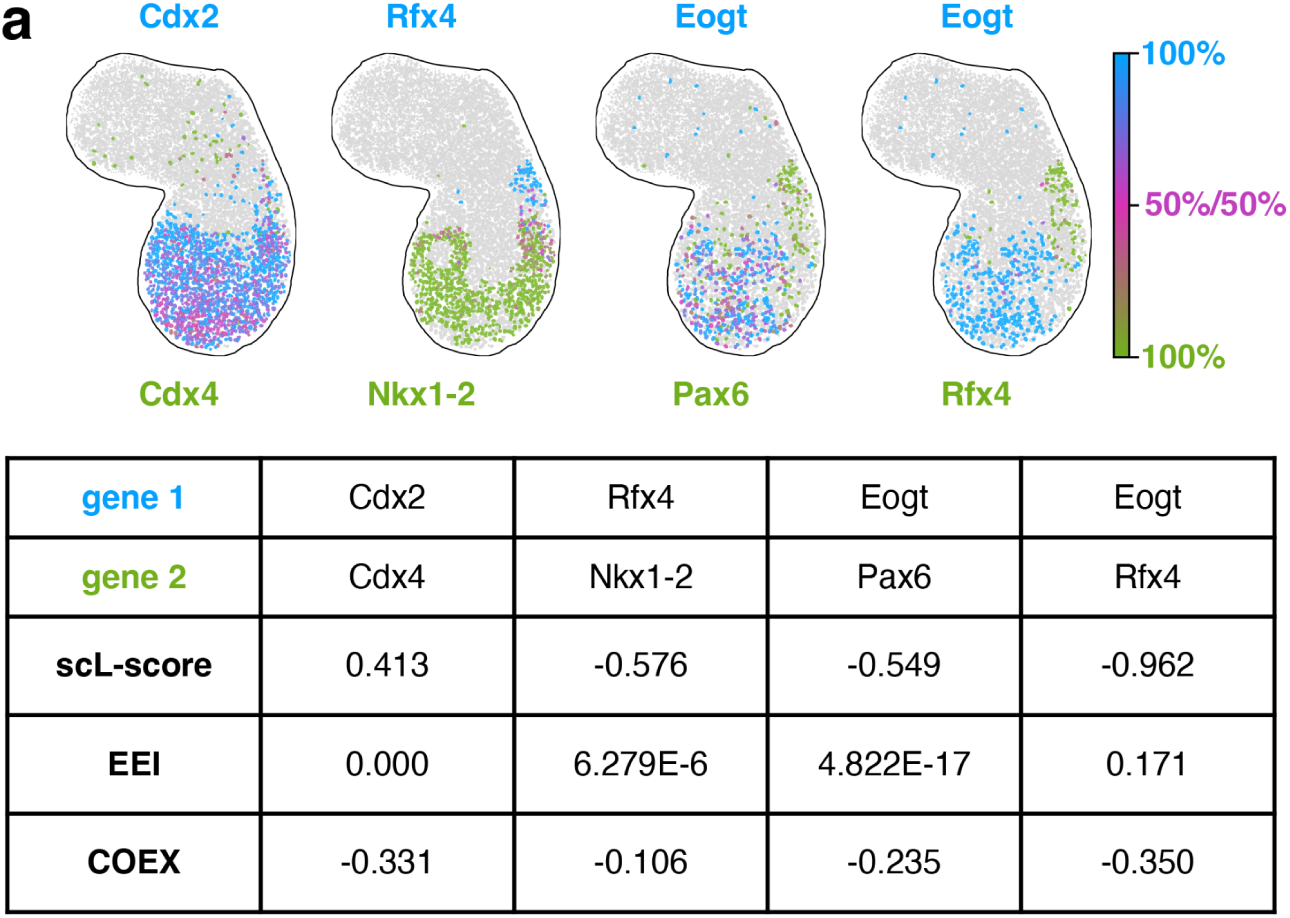
Comparison between scL-score, Exclusive Expression Index (EEI), and coefficient of coexpression (COEX) for 4 example gene pairs. **a.** Expression of 4 example gene pairs in a single gastruloid (top) and their calculated scL-score, EEI, and COEX values (bottom). We calculated the scL-score, EEI, and COEX for four gene pairs from one gastruloid sample (2025-05-07_roi2). The scL-score consistently delineated gene pairs possessing opposing expression profiles, while EEI was not always able to measure those exclusivity patterns (Pax6-Eogt: scL-score=-0.572 and -0.526, EEI=0; Rfx4-Eogt: scL-score=-0.970 and -0.955, EEI=0.171; Nkx1-2-Rfx4: scL-score=-0.537 and -0.615, EEI∼0). Only the scL-score was able to detect the coexpression pattern present between the positively associated expression profiles of Cdx4 and Cdx2 (scL-score=0.413, EEI=0, COEX -0.331) (shown visually in **Figure S3.4a**). Thus, while EEI was informative for measuring gene expression relationships characterized by mutual exclusivity, the scL-score more clearly separated positive, random, and mutually exclusive relationships on a single signed bounded scale. The COEX value trended in the opposite direction than expected, but this may be due to the fact that it cannot be calculated on single gene pairs and necessarily uses information from the entire count table, which here only consisted of 6 genes. These comparisons combined with the simulations in **Figure S3.3**, clarify a conceptual advantage of the L-score over the EEI and COTAN frameworks. In contrast, by leveraging the ranked structure of transcript counts across cells, the L-score framework does not binarize expression and does not require fitting a parametric distribution. It can be calculated on single gene pairs, and is more sensitive to mutual exclusivity.

**Figure S3.5:**
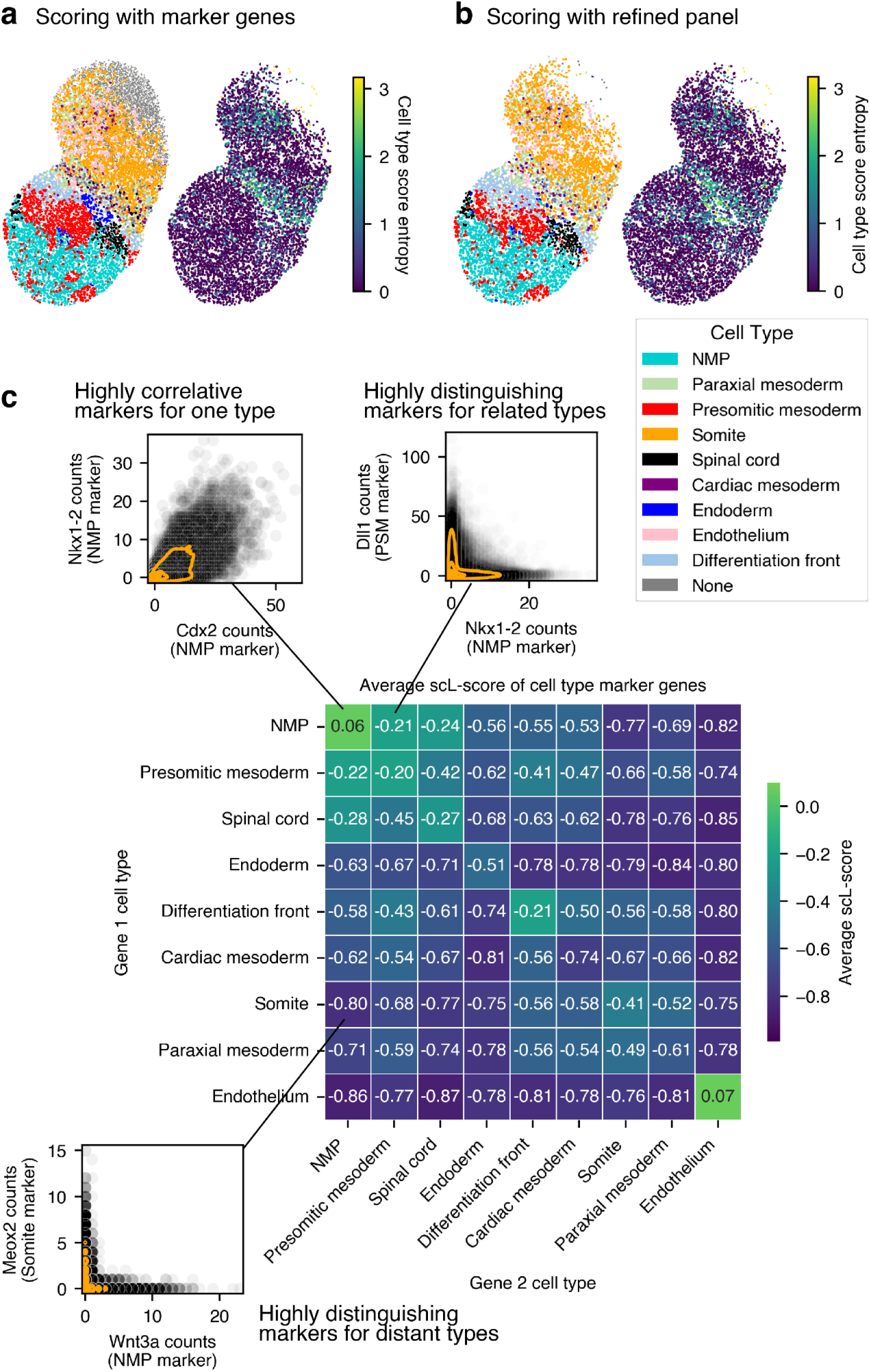
Refining and assessing marker gene panels with scL-score values. **a.** A representative gastruloid with cell types as scored by the full marker gene panel (left) and entropy values of the score distributions of each cell (right). **b.** The same gastruloid scored with a reduced panel of marker genes filtered by the properties of their scL-score values with all other genes. **c.** Heatmap showing the average scL-score values between sets of marker genes from the hand-curated panel used in this paper. Panels were filtered by average per-cell expression, see Methods for details.

**Figure S3.6:**
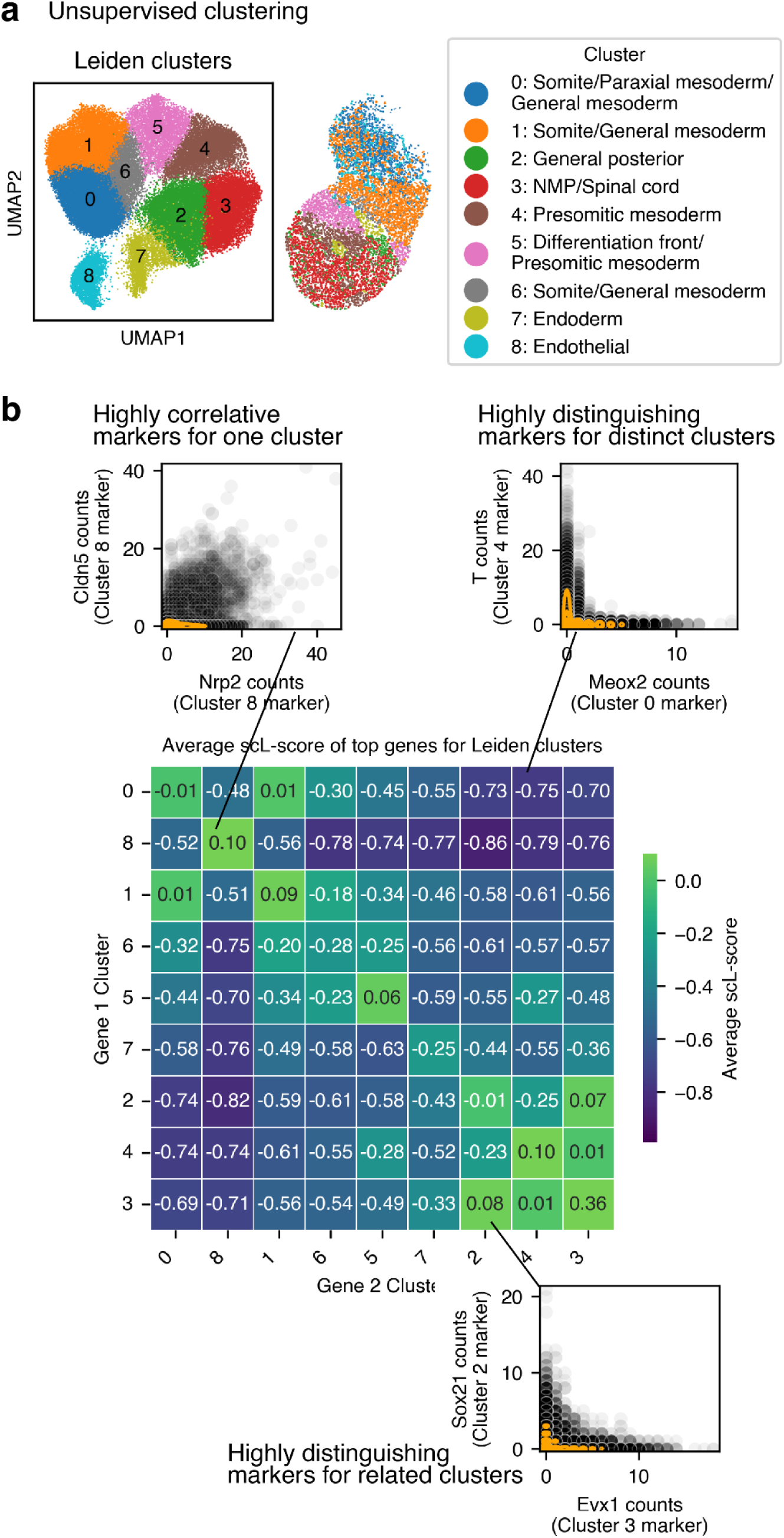
Using the scL-score to assess unsupervised clustering and identify marker genes. **a.** Left: UMAP of all gastruloid nuclei from the dataset taken on 05/07/2025 with > 40 counts projected onto a UMAP and colored according to Leiden clusters. Right: Cluster assignment for a representative gastruloid projected into spatial coordinate. **b.** Heatmap showing the average scL-score values for the top 10 genes associated with each cluster shown in **a**. We reasoned that since the L-score quantifies mutually-exclusive gene expression, a property of good marker genes, it might be possible to use a gene’s scL-score values to refine our marker gene panel. We calculated the scL-score vectors, and then filtered for only genes which had at least 10% of their L-score values < -0.8, and at least 12% > -0.3. This did shift the balance of differentiation front and presomitic mesoderm genes, however it slightly increased the entropy scores (**Figure S3.5a,b**), indicating that for small, hand-chosen marker gene panels, removing genes decreases scoring confidence, albeit only slightly. However, we also used the average scL-score of genes associated with one type compared to another type to quantify type similarity (in terms of per-cell gene expression), and found marker pairs that distinguished cell types or were concordantly expressed in the same cell types (**Figure S3.5c**). This method can also be used to post-facto analyze similarity of clusters produced by Leiden or other clustering methods. To demonstrate this we pseudo-bulked all the nuclei for our entire dataset and clustered using a standard scanpy workflow (**Figure S3.6a**). We then took the top genes for each cluster, and calculated the average scL-score between the groups (**Figure S3.6b**). This analysis demonstrated that the clusters produced by unsupervised clustering, while they could be qualitatively mapped onto the cell types we expected to see (**Figure S3.5a**), were much more similar in terms of gene expression. Just using the top genes from this analysis would not be sufficient to produce marker genes. However, we used scL-score values to find genes that were divergently expressed, even among similar clusters (**Figure S3.6b**).

**Figure S4.1:**
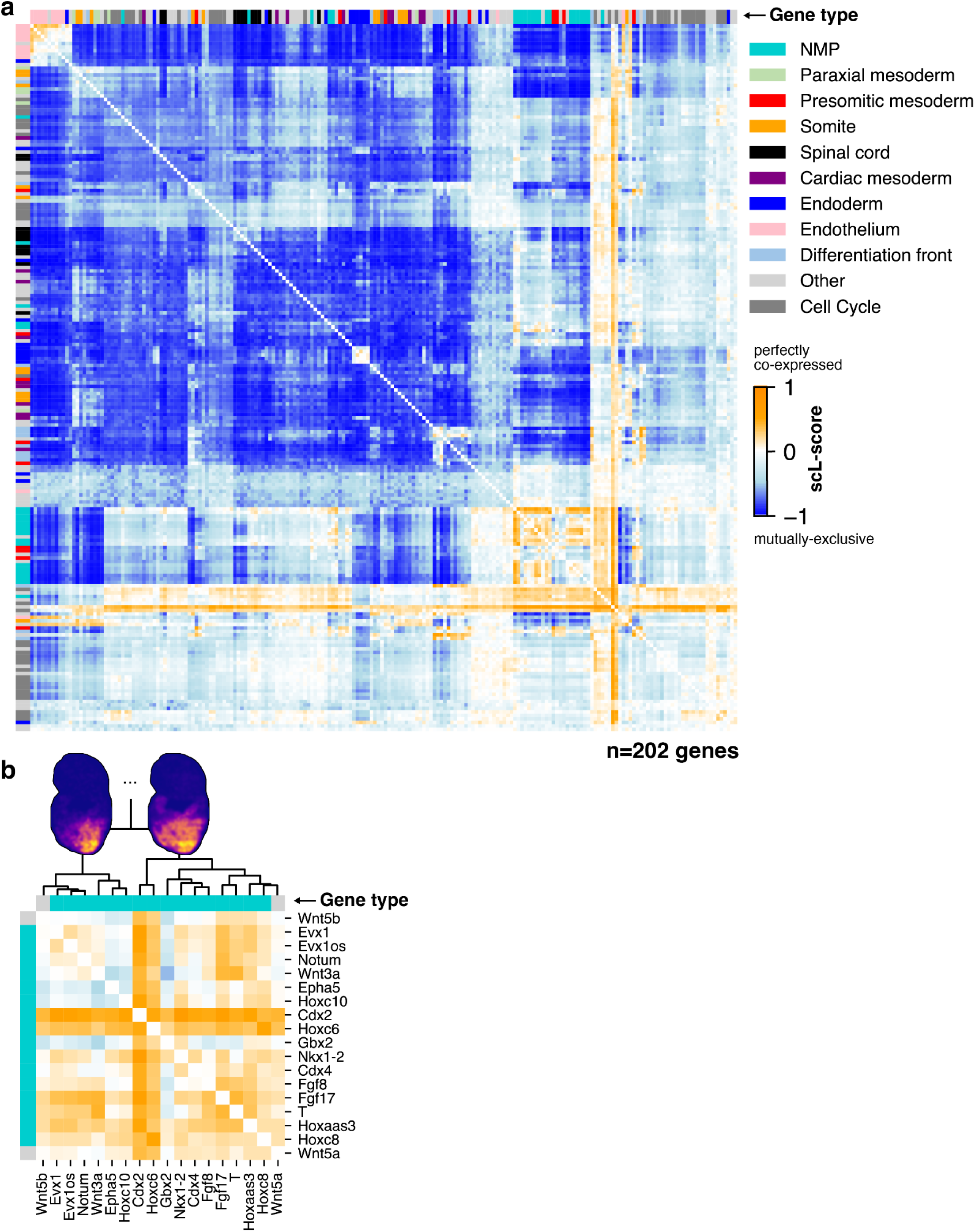
Heatmap of L-score values including cell cycle genes and NMP subcluster. **a.** Heatmap of all scL-score values for all genes in the panel (excluding poorly detected but including cell cycle genes n=202 genes). Colored bars on the top and right hand sides indicate if a gene is associated with a particular cell type. Heatmap was hierarchically clustered by row. **b.** Highlight of the NMP cluster from Figure 4a. Clustering relationships are indicated with the dendrogram, and summed densities for all genes in an example gastruloid show where the genes in each cluster are expressed spatially. Clustering relationships determined from the full gene set shown in Figure 4a.

**Figure S4.2:**
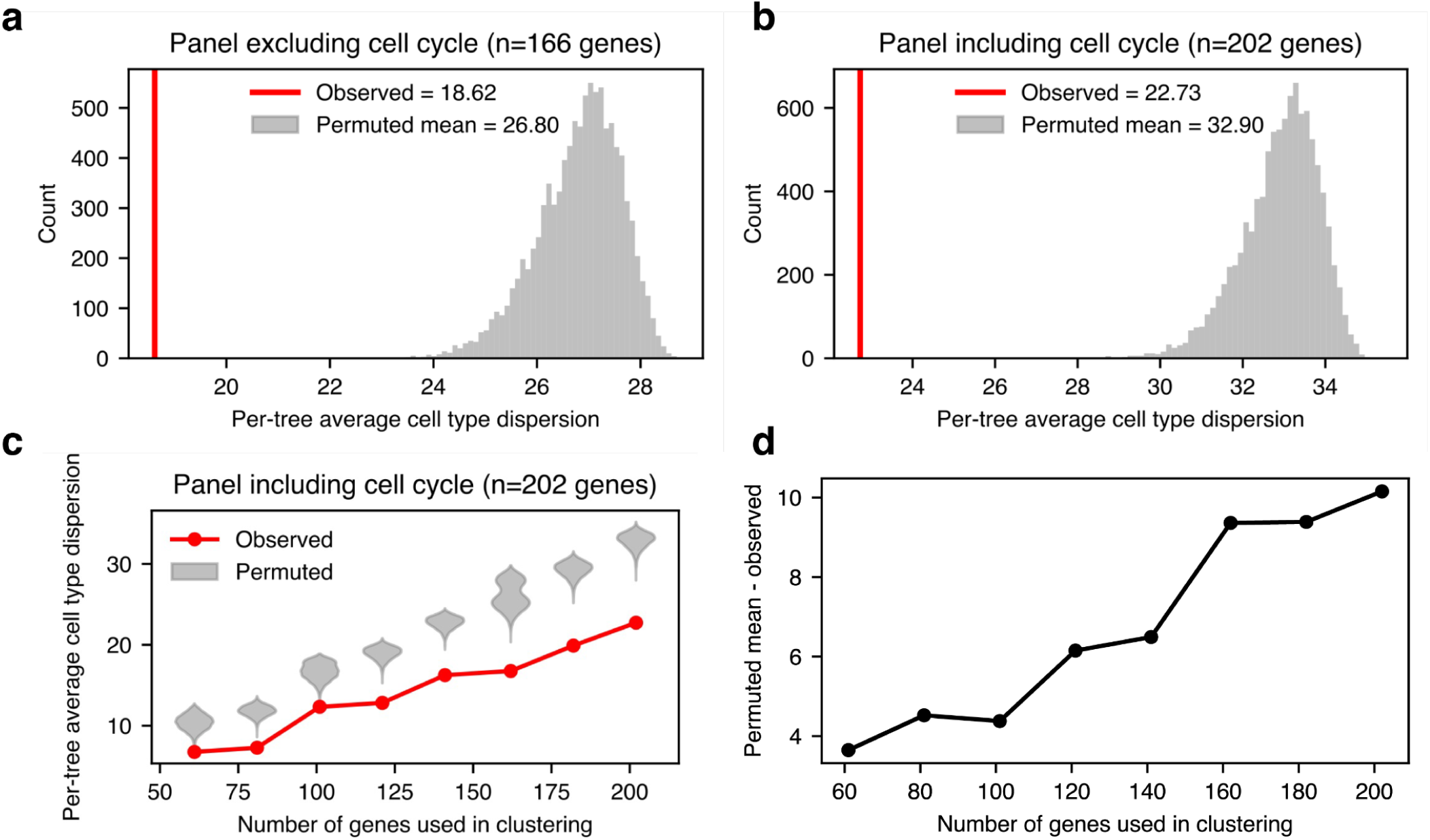
Quantification of cell type clustering by scL-score analysis and robustness of clustering to number of genes included. **a.** Average intra-cell type mean pairwise branch length (cell type dispersion) for the tree produced by scL-score clustering on 166 well-detected genes (excluding cell cycle genes) for real (red line) and permuted (gray distribution) leaf identities. **b.** Cell type dispersion of the tree produced by scL-score clustering on 202 well-detected genes (including cell cycle genes) for real (red line) and permuted (gray distribution) leaf identities. **c.** Cell type dispersion of observed (red) and permuted (gray) leaf identities as a function of the number of genes used to produce the underlying tree. **d.** Gap between the mean permuted cell type dispersion and the real cell type dispersion as a function of the number of genes used to produce the underlying tree.

**Figure S4.3:**
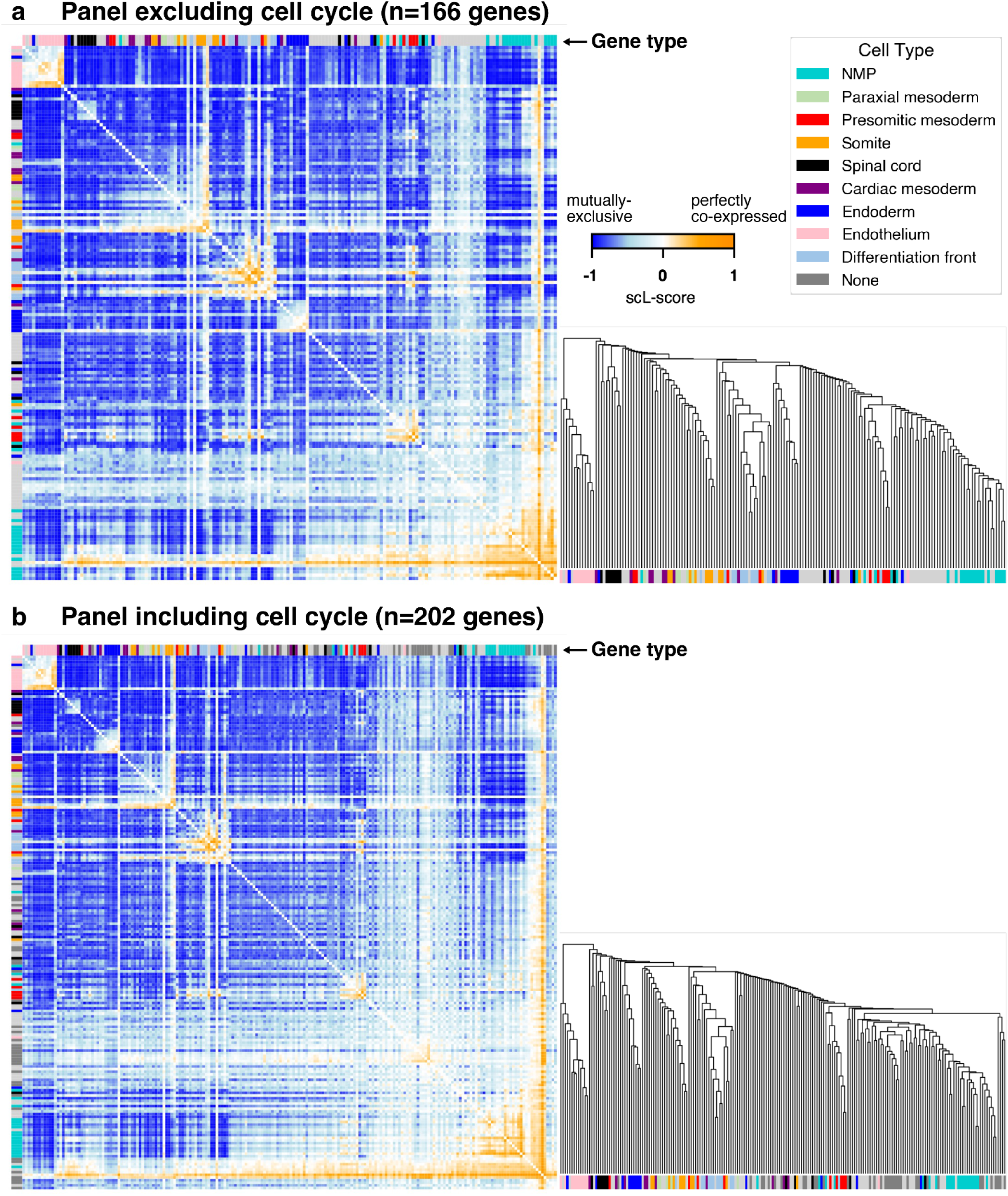
Clustering on transformed scL-scores. **a.** scL score values for all pairs of well-detected non-cell cycle genes (n=166). Row order is determined by hierarchical clustering on transformed scL-score values: (1-scL)/2. Values in the heatmap on the left are the untransformed values, the dendrogram on the right shows the clustering relationships. Column order was set to be the same as the row order. **b.** scL score values for all pairs of well-detected genes (n=202). Row order is determined by hierarchical clustering on transformed scL-score values: (1-scL)/2. Values in the heatmap on the left are the untransformed values, the dendrogram on the right shows the clustering relationships. Column order was set to be the same as the row order.

**Figure S4.4:**
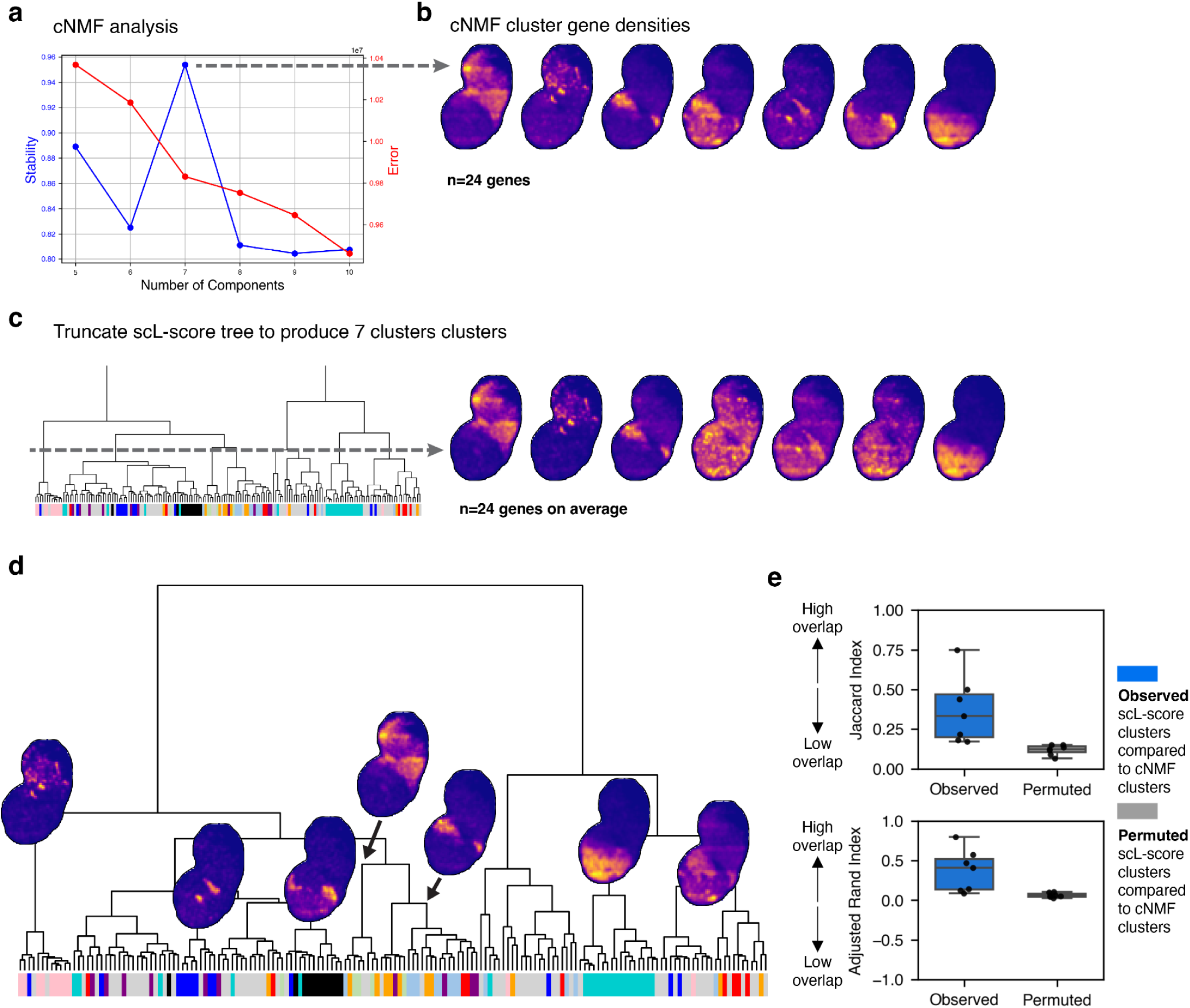
Comparison of scL-score clusters to cNMF clusters. **a.** cNMF stability analysis for varying numbers of components. cNMF was run with standard parameters on all nuclei from the dataset taken on 05/07/2025. **b.** Top 24 genes in each cluster averaged in space and summed together for visualization. 24 genes were chosen as this was the average number of genes per cluster when the L-score tree was truncated to produce 7 clusters for comparison (see **c**) **c.** Truncation of the scL-score tree to produce 7 clusters. The genes in each cluster were then visualized as in **b**. **d.** Qualitative clusters on the scL-score tree that have many genes associated with a single cell type. **e.** Quantification of the overlap between scL-score clusters and cNMF clusters. Overlap between each scL-score cluster and each cNMF cluster was quantified by Jaccard Index or Adjusted Rand Index. The scL-score clusters were then shuffled and the same values were quantified. The value of the overlap between each scL-score cluster (blue) or permuted scL-score cluster (gray) and the most similar cNMF cluster is shown. We ran cNMF (Kotliar et al., 2019) on our data to identify gene programs. Stability analysis indicated that 7 was the optimal number of components. To compare how the genes identified by cNMF compared to those which are related by scL-score similarity, we truncated the L-score tree to produce 7 clusters. We visualized the genes for each method in space. There was clear correlation between the clusters produced by both methods. The cNMF clusters looked ’cleaner’ (because the top 24 genes in each program were plotted, and thus many genes that did not meet this threshold in any program weren’t visualized at all), while the L-score clusters had a hierarchy of relatedness which was absent from the cNMF programs. Furthermore, we could select tree nodes qualitatively as being enriched for cell type marker genes, and found that these clusters also closely matched the cNMF clusters.

**Figure S4.5:**
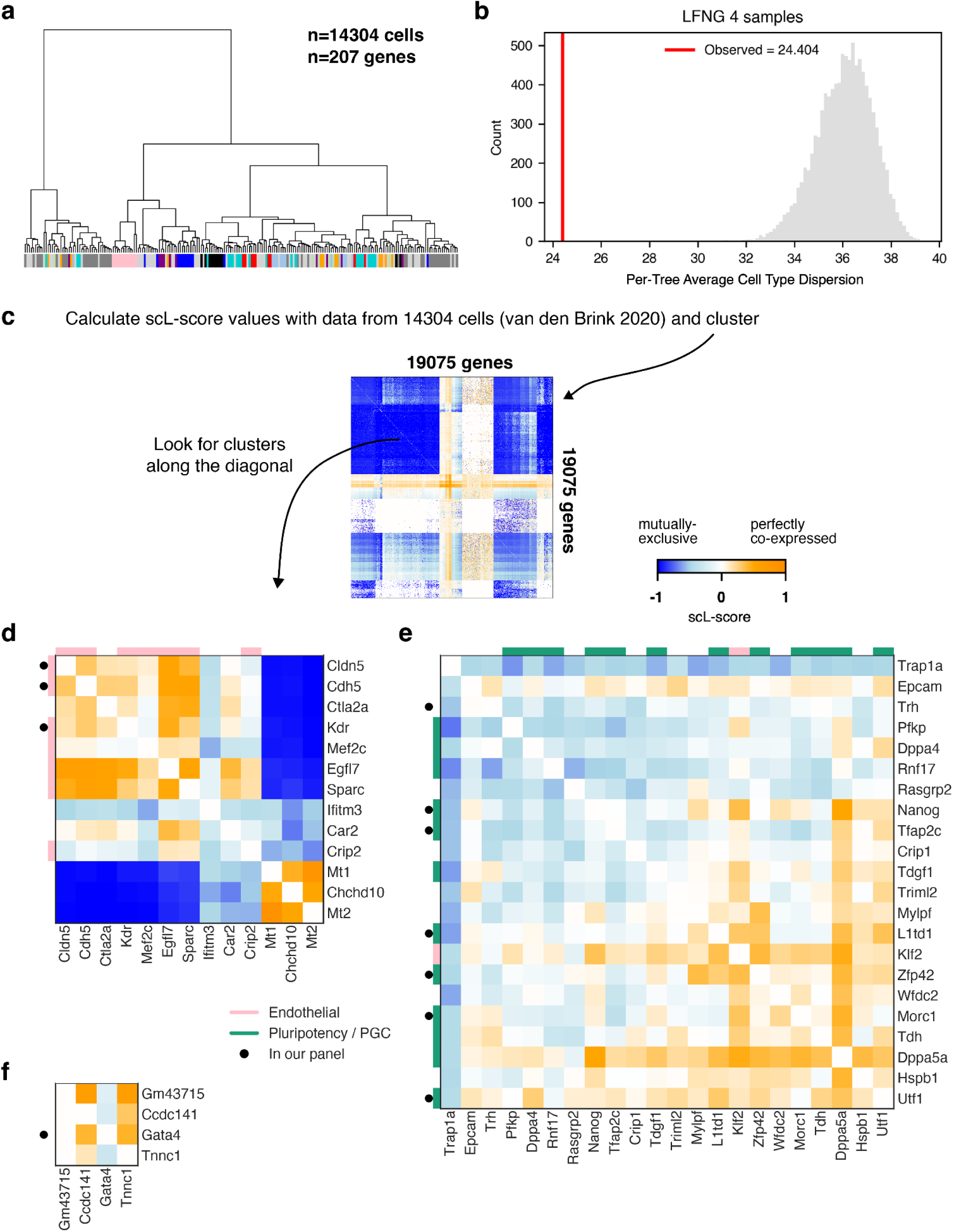
scL-score analysis of scRNA-seq gastruloid dataset. **a.** Dendrogram produced by hierarchical clustering the scl-scores of the 207 genes in the panel in the work as measured by sc-RNAseq in (van den Brink et al., 2020): 4 samples of 120 hour gastruloids were pooled; after filtering for read quality 14304 cells remained. scL-score difference vectors were used as the distance measure for clustering as in Figure 4. **b.** Cell type dispersion (average pairwise branch length per cell type) of the real (red line) and permuted (gray distribution) tree shown in **a**. **c.** scL score heatmap of values calculated from (van den Brink et al., 2020); the same quality control, filtering, and clustering method was performed as is shown in a), but all well-detected genes (19075) were used to perform clustering. The scL-score value, rather than the distance, is shown for each gene pair. **d.** Example cluster found on the diagonal of the heatmap shown in **c**. **e.** Example cluster found on the diagonal of the heatmap shown in **c**. **f.** Example cluster found on the diagonal of the heatmap shown in **c**.

**Figure S5.1:**
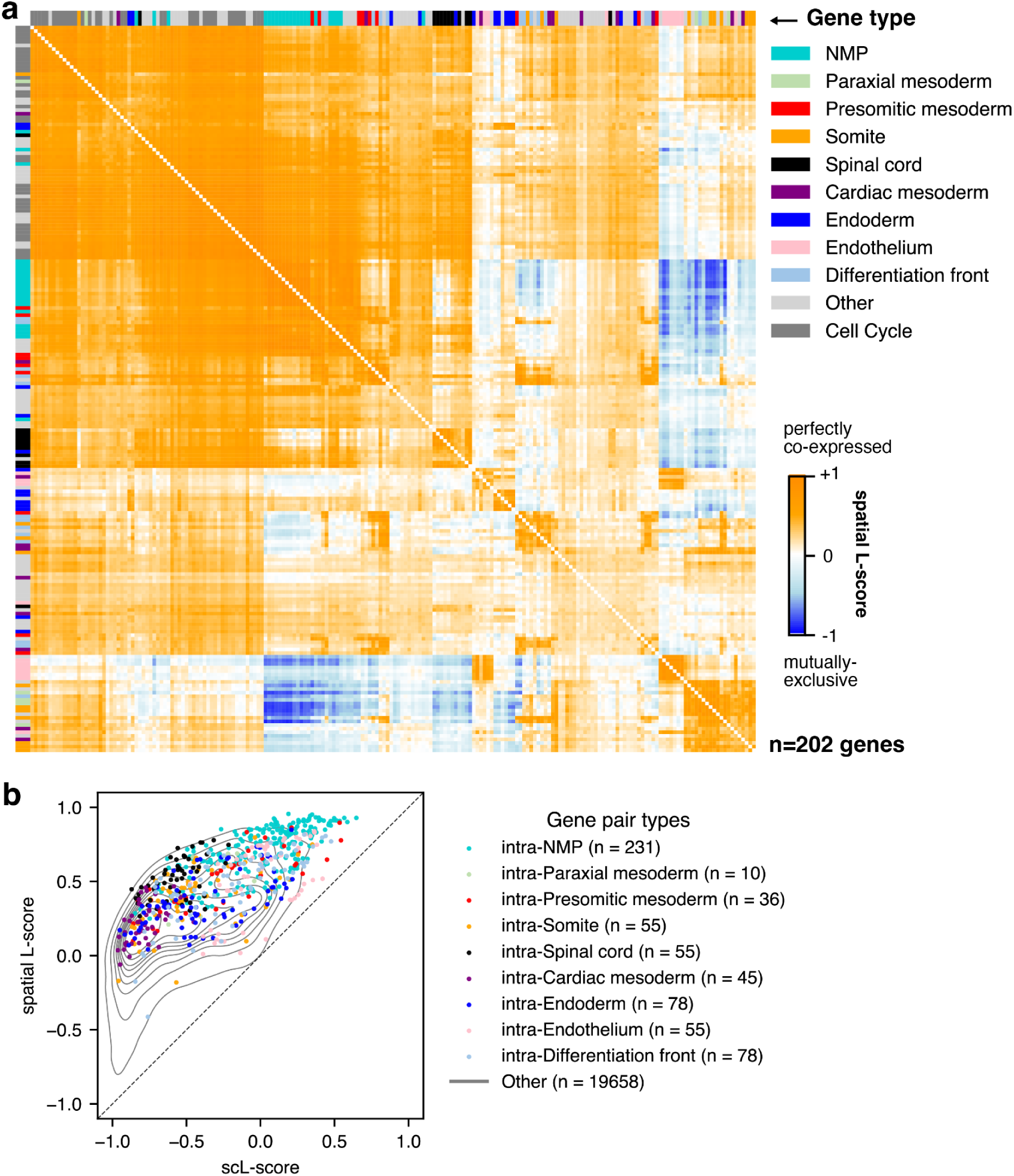
Spatial L-score heatmap of values averaged across all samples and direct comparison of scL-score and spatial L-score values for all gene pairs. **a.** Heatmap of clustered spatial L-score values for all genes averaged across all gastruloids. n=202 genes. **b.** Comparison of scL-score and spatial L-score values for cell type markers. Colored dots represent intra-type pairs, with the color specifying the cell type. The gray trace is the smoothed density estimate for all pairs, including inter-type pairs and pairs including cell cycle or non-marker genes.

**Figure S5.2:**
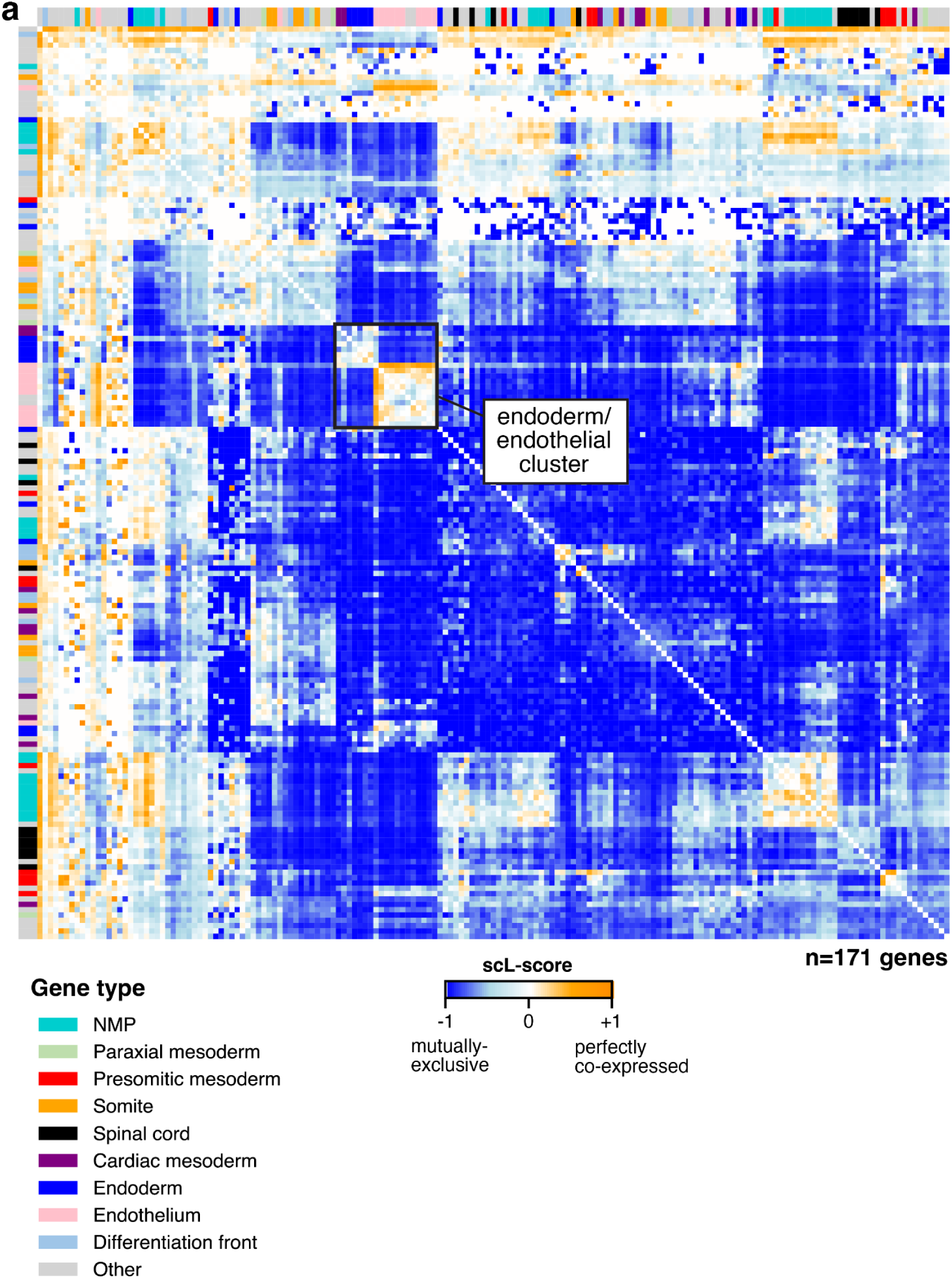
Clustered heatmap of the scL-score for all genes in an example gastruloid. **a.** Heatmap of clustered scL-score values for all genes for the representative gastruloid shown in the main figure. Purple rectangle highlights endoderm and endothelial gene clusters.

**Figure S6.1:**
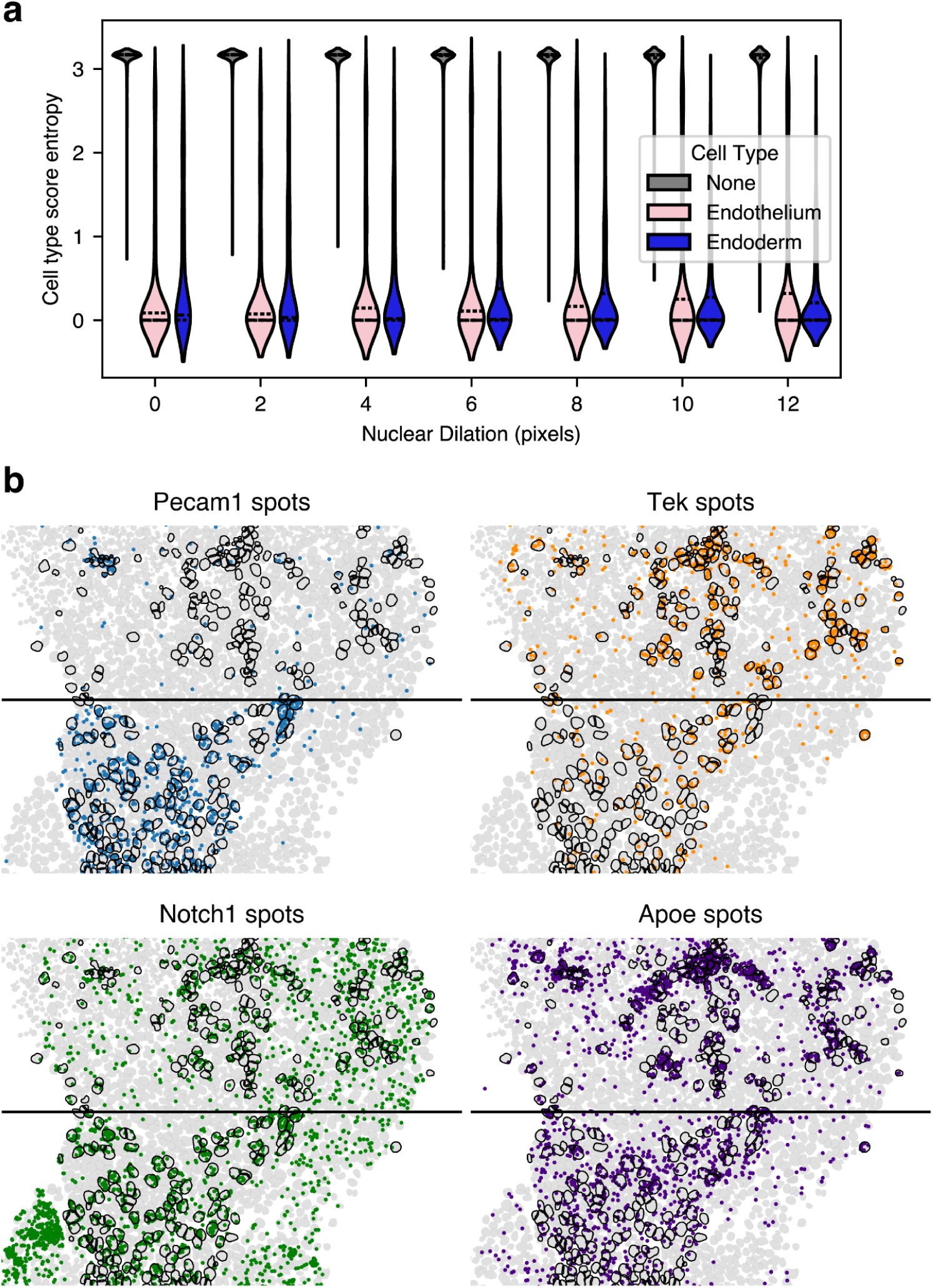
Estimation of the effect of spot mis-assignment. **a.** Cell type entropy score distributions as a function of nuclear dilation (higher = more pixels added to the original segmentation). Nuclei from the gastruloid in Figure 6a are shown. **b.** Expanded images from Figure 6e**, f**.

**Figure S6.2:**
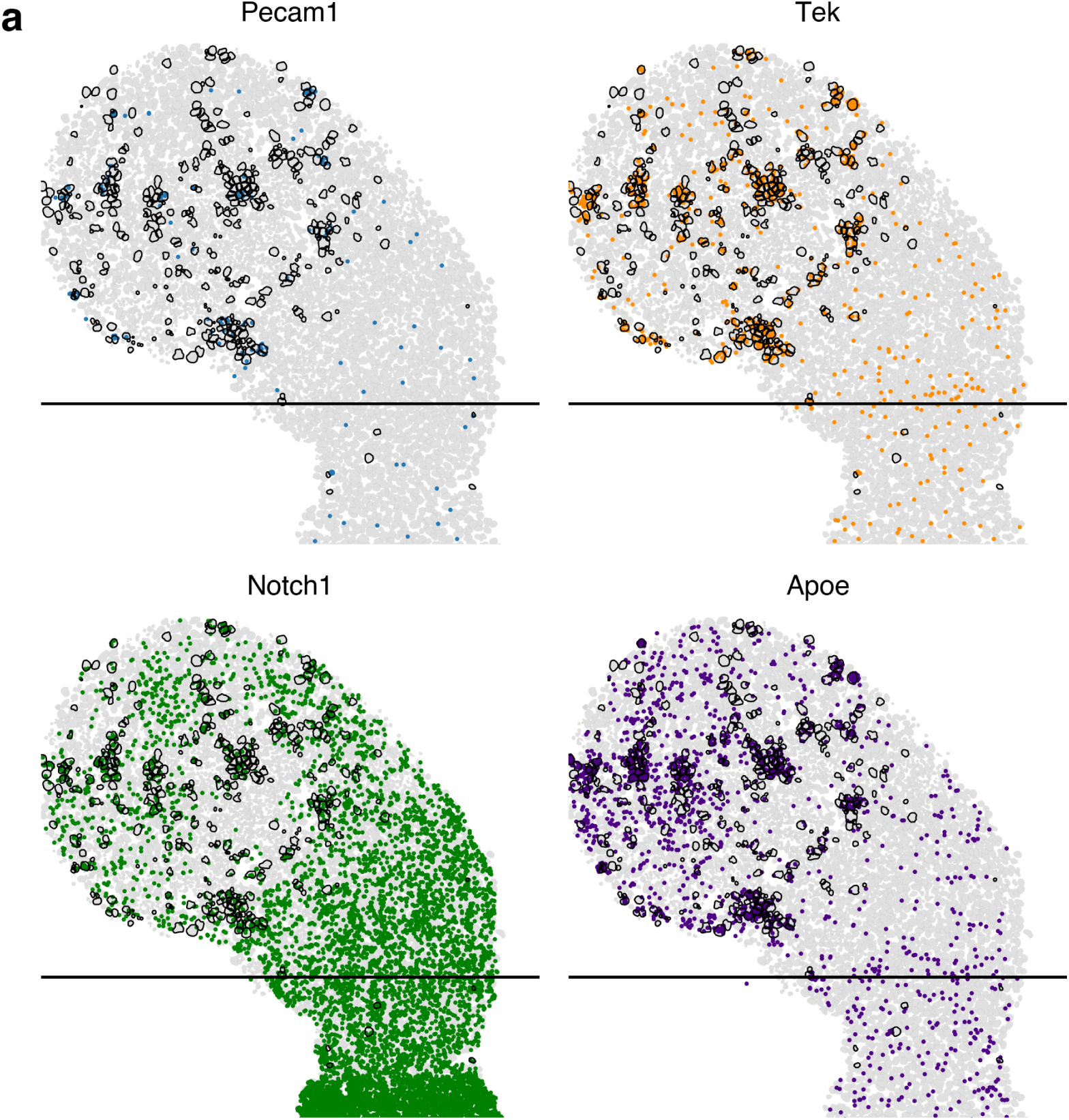
Expression of differentially expressed endothelial genes in a gastruloid with only one endothelial population. **a.** Expanded images showing the expression of the same genes highlighted in Figure 6e**, f** in the gastruloid shown in Figure **S1.3a iii**.

## Methods

### Cell culture

All gastruloids were generated from E14TG2a cells (a generous gift from the Toettcher lab). Cells were cultured in 2i+LIF media at 37 °C, 5% CO_2_ and grown on plates coated in 0.1% gelatin (Millipore/Sigma ES006B). 2i+LIF media consisted of GMEM (Gibco 11710-035), 10% stem-cell qualified FBS (R&D Systems S10250), 1x GlutaMAX (Gibco 35050–061), 1x MEM non-essential amino acids (Gibco 11140050), 1 mM sodium pyruvate (Gibco 11360–070), 100 μM 2-mercaptoethanol (Gibco 471 21985–023), and 100 units/mL penicillin/streptomycin (Gibco 15140–122), 1000 units/mL LIF (Millipore Sigma ESG1107), 2 μM PD0325901 (Tocris 4192), and 3 μM CHIR99021 (Tocris 4423). Cells were passaged every 2 days: cells were trypsinized (Life Technologies 12563011), counted using an automated cell counter (Invitrogen Countess II), and 5x10^-5^ — 8x10^-5^ cells were transferred to a 25cm^2^ dish; cells were passaged at least once after thawing before gastruloid aggregation.

### Gastruloid aggregation

The aggregation protocol was as described in (van den Brink et al., 2020) (<u>protocol here</u>). Briefly, cells were trypsinized, rinsed twice with 1X PBS to remove media, and then resuspended in N2B27 medium. N2B27 consisted of equal parts neurobasal medium (Gibco 21103049) and DMEM/F12 (Gibco 11320033) supplemented with 1X N2 (Gibco 17502048) and B27 (Gibco 17504-044), 100 units/mL penicillin/streptomycin (Gibco 15140–122), and 100 μM 2-mercaptoethanol. Cells were resuspended at a concentration of 2500-7500 cells/mL; 40 μL of this mixture were placed in each well of round-bottom ultra non-adherent U-bottom 96-well plates (Costar 7007), yielding 100-300 cells/well. The first two seqFISH runs used 300-cell gastruloids (n=8 total) and the last used 100-cell gastruloids (n=18). Cells were allowed to grow at 37 °C, 5% CO_2_ for 48 hours and then supplemented with 150 μL fresh N2B27+3 μM CHIR99021 (Chiron, Wnt agonist). After 24 hours, 150 μL of media was removed and replaced with N2B27 without Chiron; this media change was repeated daily until the gastruloids reached maturity at 120 hours. All wells were scored for elongation at 120 hours just before fixation. The gastruloids selected for fixation were elongated and uniaxial; the experiments that generated them had an 80-94% success rate (see **Figure S1.1a** for full distribution).

### Sample fixation

After gastruloids had grown for 120 hours, they were harvested by pooling together all successful (i.e., elongated and uniaxial) wells in a plate, washed with 1X PBS, and then fixed for 10 minutes in 4% PFA in PBS. After fixation, gastruloids were washed with 1X PBS and stored in methanol for up to a month before preparation for seqFISH. 3 separate seqFISH runs were performed on 6 different aggregations (one experiment pooled 4 plates of gastruloids), producing a total of 26 individual samples.

### Embedding and sample preparation for seqFISH

∼30 gastruloids were placed in a Lo-Bind tube (Eppendorf 0030108523). After removing as much methanol as possible, the gastruloids were washed with 2XSSC, then incubated in 2XSSC + 1 μg/mL DAPI overnight at 4 °C. Stained gastruloids were permeabilized with 1X PBS + 0.2% Tween20, arrayed on a gelatin-coated slide (Spatial Genomics 00200001) with an eyelash tool (Electron Microscopy Sciences 11852), covered in liquid acrylamide gel, and allowed to polymerize for 2 hours. Polymerized samples were washed with 8% SDS, then treated with 8x10^-3^ units/mL Proteinase K (NEB P8107S). After this wash, the primary probes were hybridized for 16-40 hours, after which the samples were washed and loaded onto a GenePS machine (Spatial Genomics).

Detailed protocol available here.

### seqFISH and raw data handling

Sequential rounds of secondary probe addition and imaging were performed in an automated fashion by the instrument (Eng et al., 2019). The GenePS software that controlled the instrument was v1.6.0. The instrument was run with a 1 or 2 μm z-offset on 5-18 individual regions of interest, highlighting each gastruloid. The genes chosen for the seqFISH panel were based on differential expression in single-cell RNA sequencing experiments in references (Rosen et al., 2022; van den Brink et al., 2020) or by consensus in the literature. 200 of the genes were detected via a sequential barcoding scheme (see (Eng et al., 2019) for details), and 20 were serially detected as in traditional smRNA-FISH (Raj et al., 2008). The serial genes tended to vary in quality between gastruloids; genes with high background or in which expression was too high for the spot detection software to confidently assign spots were excluded. The list of excluded genes for each gastruloid is included with the raw data in a file called ‘bad-genes.txt’.

### Gastruloid annotation and spot assignment

Detected spots were decoded and exported without nuclear assignment by SG Navigator software version 0.9.0. At this stage, gastruloids without even and consistent expression of the housekeeping gene Eef2, which indicated focus or technical issues or sample variability, were discarded. 5 out of 6 gastruloids were discarded from the first run, and 3 out of 11 in the second run. None of the samples from the third run were discarded. Raw TIFF files of the DAPI images were annotated using the NimbusImage platform, which is available here: https://github.com/arjunrajlaboratory/NimbusImage/ and hosted here: https://www.nimbusimage.com/. Expression of the gene T was used to locate the posterior end and corroborated by consistent differences in nuclear morphology between the posterior and anterior. An AP axis was inferred by drawing a line between that pole and the extreme opposite end of the gastruloid along the center of the gastruloid parallel to the edges and connecting with the furthest extended point to define the anterior pole. Segmentation was performed using cellposeSAM (Pachitariu et al., 2025) on the high-resolution .tiff files of the DAPI-stained nuclei from the 3rd round of hybridization. Once the nuclear masks had been created, spots were assigned to nuclei using a custom software package called SGAnalysis (<u>available here</u>). Nuclei were dilated by 8 pixels (0.86 μm), and all spots that fell within this dilated nuclear area were assigned to that nucleus. In cases where the dilated regions overlapped, spots were assigned to the nucleus for which the edge of the dilated area was furthest away. We were able to resolve 131,112 nuclei total, with an average of 5043/gastruloid. 79,607 nuclei had enough mRNA counts to be typed (mean 3062/gastruloid). Nuclei without enough mRNA were generally small, uniformly bright (in contrast to the usual mottled appearance of mouse nuclei), and lacked expression of the highly-expressed housekeeping gene Eef2, indicating the corresponding cell was likely dead or fragmented. We primarily found these nuclei in the anterior tissues and classified them as “none”-type, excluding them from our analyses unless noted. The average reads/nucleus per gastruloid ranged from 47-80; for typed nuclei the value was 105-265.

### Morphological characterization of gastruloids

To assess gastruloid morphology, 4x images taken with an Incucyte, were segmented in Nimbus Image (https://www.nimbusimage.com/). Gastruloids were imaged directly in a 96-well U-bottom dish: 1 plate from 1/19/2024, 1 plate from 9/1/2024, and 4 plates from 4/4/2025 containing 88, 86, 93, 84, 88, and 90 gastruloids respectively. The Nimbus feature ‘Calculate blob metrics’ was used to quantify the following: "solidity”, "rectangularity", "perimeter", "elongation", "eccentricity", "convexity", "circularity", and "area" (code can be found at https://github.com/arjunrajlaboratory/NimbusImage/). These metrics were then decomposed by PCA. All images and annotations can be found at the following link: https://app.nimbusimage.com/#/project/69d3fa8f1be4701f5fab6359.

### Cell type scoring

To assign cell type, we used a panel of marker genes for 9 consensus cell types thought to exist in gastruloids at this stage of development: Neuromesodermal precursor (NMP), presomitic mesoderm (PSM), spinal cord precursor (spinal cord), endoderm, differentiation front, cardiac mesoderm, paraxial mesoderm, somitic, and endothelial precursor (endothelium). Genes were chosen based on differential expression in references (Rosen et al., 2022; van den Brink et al., 2020), or consensus in the literature. Each nucleus received a score for each cell type using the score_genes function in scanpy (Wolf et al., 2018), and the highest-scoring cell type category was assigned to that cell. If none of the scores met a universally applied cutoff (0.1), then the cell was assigned ‘none’.

### Cell type entropy score

The degree of uncertainty in cell typing was quantified by calculating the entropy of the cell type score distribution. The scores can be negative or zero, so they were first converted to probabilities using a softmax function with temperature scaling:

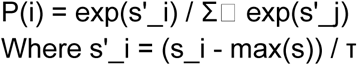

A temperature of 0.1 was used. Once the scores had been converted, the Shannon entropy of the distribution was calculated:

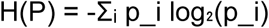

### Cell type proportion variation

Since the proportion of each cell type in a gastruloid must sum to 1, changes in one cell type will affect at least one other cell type within the same sample. To properly account for this when applying statistical tests, we calculated the center log ratio (CLR)-transformed proportion and tested all pairs of cell types for significant variation. Statistical significance of pairwise correlations between CLR-transformed cell type proportions was assessed using Pearson correlation coefficients with associated p-values. To account for multiple comparisons, p-values were adjusted using the Benjamini-Hochberg procedure to control the false discovery rate (FDR). Correlations with an adjusted p-value < 0.05 were considered statistically significant.

### Computation of AP axis location

Two methods of normalization were used to define position along the AP axis. Both use a hand-drawn annotation of the axis from posterior to anterior as inferred from morphology and T-expression. All annotations are available; see Data Availability section below. For the first method (length-based), the drawn AP axis was normalized from 0 to 1. Each nucleus was projected onto the closest point on this normalized line. For the second method, the ends of the axis were still defined as 0 and 1, but rather than uniformly binning the space, a midpoint (0.5) was defined as the 90th percentile of T expression. The total amount of T spots was used, rather than the per-nucleus amount. The length from 0 to the T-midpoint was uniformly binned, as was the length from the midpoint to 1. The same projection method was used to find the AP axis location of each nucleus.

### Quantification of Cell Type Spatial Relationships: Exposure index

To characterize the spatial organization of cell types, we computed two related measures: a pairwise exposure index matrix capturing type-specific spatial relationships, and a scalar mixing index summarizing overall spatial integration. For each cell, we identified neighbors as all cells whose centroids fell within a specified radius r of the focal cell’s centroid. We chose a value of r of 16 μm, which gave an average of 5-6 neighbors per cell. We chose this value as it captures local interactions, which was the primary goal of these analyses. For each ordered pair of cell types (s, t), we calculated the exposure index as the proportion of type s cells’ neighbors that are type t:

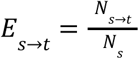

where Ns→t denotes the count of neighbor pairs in which the focal cell is type *s* and the neighbor is type *t*, and Ns denotes the total number of neighbors across all type *s* cells. Each row of the resulting exposure matrix sums to unity and represents a probability distribution over neighbor types for a given source type. To account for differences in cell type abundance, we normalized exposure indices relative to the expectation under random spatial arrangement:

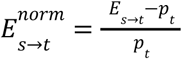

where pt is the proportion of cells that are type *t*. Normalized values of zero indicate exposure consistent with random mixing, positive values indicate spatial attraction (co-localization), and negative values indicate spatial avoidance, with a minimum of −1 representing complete exclusion.

### Quantification of Cell Type Spatial Relationships: Mixing index

To summarize overall spatial integration across all cell types, we computed a mixing index defined as the fraction of neighbor pairs involving different cell types:

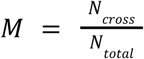

where N_cross is the number of neighbor pairs involving cells of different types and N_total is the total number of neighbor pairs. For normalization, we compared the observed mixing to the expectation under random spatial arrangement:

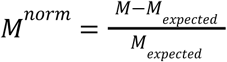

where

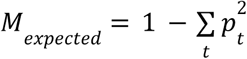

is the expected cross-type interaction rate given cell type proportions. Normalized values of zero indicate random spatial mixing, positive values indicate greater integration than expected (hyper-mixing), and negative values indicate spatial segregation.

### Exclusion of untyped cells

When computing the mixing index, cells lacking confident type assignments were optionally excluded from both the numerator and denominator, ensuring the measure reflects only spatial relationships among typed cells. These cells were retained in the exposure matrix to quantify how typed cells interact with unclassified cells.

### Statistical analysis of variance

To identify cell type pairs whose spatial relationships varied significantly across samples, we computed the variance in exposure indices across samples for each type pair. To account for the expected relationship between mean exposure and variance, we regressed log-variance against log-absolute-mean across all type pairs and computed residuals. Type pairs with residual variance exceeding the 97.5th percentile (two-tailed α = 0.05) were considered significantly variable, indicating spatial relationships that differ across samples beyond what is expected from sampling variation and composition differences.

### Triplet identification and enrichment

Spatial enrichment of cell type triplets was assessed using a computational framework based on Delaunay triangulation. Each gastruloid was processed independently to account for potential inter-sample variability in cell density and spatial organization. Spatial neighbors were identified using Delaunay triangulation, which partitions the coordinate space into triangular tessellations connecting nearest neighbors. A distance cutoff of a maximum of 200 pixels was used; neighbors with greater than this distance were not considered. All groups of 3 nearest neighbors were used to evaluate triplet frequency. Statistical significance was evaluated through a permutation testing framework. For each sample, the observed triplet frequencies were compared against null distributions generated through random reassignment of cell type labels while preserving spatial coordinates. The number of permutations was set to 1,000 per sample. The data for all gastruloids were compiled together such that the significance of each triplet in each sample, and how many samples each triplet was found in, could be assessed. Only significantly enriched or depleted triplets were analyzed; significant triplets found in at least half of the gastruloids are shown in **Figure 2c**.

### Kernel density estimation for generating smoothed spatial profiles of gene expression

We performed kernel density estimation (KDE) to generate smoothed spatial profiles of gene expression patterns. Using the nonparametric.KDEmultivariate method from the statsmodels package in Python, a given gene’s KDE of gene expression in a given gastruloid was calculated using a two-dimensional Gaussian kernel for each RNA spot fitted to a bandwidth equal to the average nearest neighbor distance between cell centroids in that gastruloid multiplied by 2 (Table S1) in both dimensions. To produce an expression profile for a gene from its KDE on a given gastruloid, we first partitioned the gastruloid’s annotated hull to produce a grid containing square elements, each with side length equal to half of the average cell diameter, and then evaluated the KDE on the center of each grid element. Visualizations of multiple genes’ combined expression in a given gastruloid were obtained by summing individual genes’ normalized KDE values for each grid element.

### Calculating the single-cell L-score

The single-cell L-score (scL-score) was computed for each ordered gene pair (gene A, gene B) within a single gastruloid. The cell-by-gene expression matrix was filtered to retain only cells with at least 2 detected transcripts for both genes. Cells were sorted in descending order by gene A’s expression values; gene A thus serves as the reference distribution, and the score is asymmetric with respect to gene order.

Three reference distributions were constructed for gene B: (1) perfect coexpression, in which gene B’s values were sorted in the same descending order as gene A; (2) perfect mutual exclusivity, in which gene B’s values were sorted in ascending order; and (3) independence, in which every cell was assigned the mean expression value of gene B.

Cumulative sums of expression values were computed for gene A, for the observed expression of gene B, and for each reference distribution. The cumulative sum of each gene B distribution (observed and reference) was then plotted against the cumulative sum of gene A. This cumulative-sum-versus-cumulative-sum representation captures how gene B’s expression accumulates relative to gene A’s: if gene B’s expression is concentrated in the same high-expressing cells as gene A, gene B’s cumulative curve rises steeply at first; if concentrated in opposite cells, the curve rises steeply at the end. The area under each curve was calculated using trapezoidal integration and normalized by the product of gene A’s and gene B’s total expression, yielding four normalized areas: A_observed (observed relationship), A_positive (perfect coexpression), A_negative (perfect mutual exclusivity), and A_uniform (independence). This normalization ensures that scores are comparable across gene pairs with different overall expression levels (**Figure S3.2a,b**). The scL-score was then defined as follows:

If A_observed > A_uniform: scL-score = (A_observed − A_uniform) / (A_positive − A_uniform), yielding values in (0, 1].

If A_observed = A_uniform: scL-score = 0.

If A_observed < A_uniform: scL-score = −(A_observed − A_uniform) / (A_negative − A_uniform), yielding values in [−1, 0).

A score of +1 indicates perfect coexpression, −1 indicates perfect mutual exclusivity, and 0 indicates independence.

The scL-score was computed for all gene pairs in each gastruloid from the 05/07/2025 dataset (n = 18 gastruloids). The other two datasets (n = 8 gastruloids) were excluded due to lower transcript detection quality. To generate an average scL-score matrix, the analysis was restricted to a common set of 202 genes well-detected across all 18 gastruloids, and per-gastruloid matrices were averaged. Unless otherwise noted, a symmetrized scL-score was used: scL-score_sym(A, B) = [scL-score(A, B) + scL-score(B, A)] / 2.

### Calculating the spatial L-score

The spatial L-score extends the scL-score to spatial regions. For each gene, a kernel density estimate (KDE) was fitted over all detected transcript spots and evaluated on a regular square grid spanning the gastruloid. Bin side length was set to twice the median nearest-neighbor distance between detected spots, calculated separately for each gastruloid. Spatial bins were ranked by KDE-derived density and processed identically to the scL-score calculation. Low-density bins were not filtered, as KDE smoothing produced non-zero density values throughout the imaging area. The spatial L-score was symmetrized as for the scL-score, except when displaying asymmetric heatmaps.

### Hierarchical clustering of L-score matrices

Gene-gene distances were defined as Euclidean distances between L-score vectors. Agglomerative hierarchical clustering was performed using Ward’s linkage criterion (scipy.cluster.hierarchy.linkage, method=’ward’, metric=’euclidean’). This approach operates on L-score vector differences rather than on pairwise L-score values directly, and therefore does not require the L-score itself to satisfy the properties of a mathematical distance metric; the Euclidean distance between L-score vectors is non-negative and symmetric by construction, satisfying the requirements of Ward’s method. Heatmaps display pairwise L-score values, not vector distances.

We applied this clustering procedure to the following gene sets:

1. A subset of NMP, presomitic mesoderm, and spinal cord marker genes in one gastruloid (n = 36 genes; **Figure 3g**).
2. All well-detected genes excluding cell cycle genes, averaged across all gastruloids (n = 166 genes; **Figure 4** and **Figure S4.3a**).
3. All well-detected genes common to all gastruloids, averaged across gastruloids (n = 202 genes; Figures S4.1a, S4.3b, S5.1a).
4. All well-detected genes excluding cell cycle genes in one example gastruloid (n = 171 genes; **Figures 5b, S5.2a**).
5. All well-detected genes common between our seqFISH panel and those that were detected in > 3 cells in scRNA-seq data from (van den Brink et al., 2020) (n=207 genes; **Figure S4.5a**).
6. All genes detected in > 3 cells in scRNA-seq data from (van den Brink et al., 2020) (n=19075 genes, **Figure S4.5c-f**).

In **Figure S4.3a,b** a transformation of the L-score values was used to cluster genes. The scL-scores were averaged across gastruloids as described above, and then each pairwise scL-score was transformed to a distance-like value with(1 - scL)/2. The matrix was then symmetrized as described previously. Agglomerative hierarchical clustering was performed directly on this transformed gene-gene distance matrix using Ward’s linkage criterion (scipy.cluster.hierarchy.linkage, method=’ward’, metric=’euclidean’).

### Cell type analysis via unsupervised clustering

We used scanpy (Wolf et al., 2018) to process the dataset from 05/07/2025. All nuclei were pooled together, nuclei with less than 40 total gene counts were filtered out, and then PCA (50 PCs) and UMAP embedding were performed using the standard pipeline. We performed Leiden clustering using scanpy.tl.leiden, and used visual inspection to choose a resolution at which to cluster. A resolution of 0.8 produced fairly well-separated clusters and roughly corresponded to the expected number of cell types. Clustering results and spatial projection are shown in **Figure S4.4a**.

### Filtering marker genes

We re-scored nuclei using a refined marker gene panel. We retained only genes that met the following criteria: more than 10% of all scL-score values < -0.8 (73% of all genes) and more than 12% of all scL-score values > -0.3 (90% of all genes). 70% of the genes in our panel met both. Scores with this filtered gene panel are shown in **Figure S3.5a**.

### Assessing marker gene panels and cluster-associated genes

To visually represent the relatedness of the genes either differentially expressed in clusters identified with unsupervised clustering, or the groups of genes associated with cell types in a marker gene panel, we calculated the average L-score of all possible combinations where the first gene in the L-score score belonged to the first type or cluster, and the second gene belonged to the second group or cluster. In the case of the marker gene panel, we first filtered genes on expression level, considering only genes that had > 2.3 counts / nucleus in nuclei where they were expressed; this filtering retained 84% of the genes in the panel.

### Using cNMF to find gene clusters

All nuclei were pooled together, and cNMF (Kotliar et al., 2019) was run on all nuclei and all non cycle-genes. The default parameters were used and 5-10 components were calculated. 7 components produced the highest stability, and the top 24 most positively differential (by ordered z-score) genes were used for visualization of qualitative clusters. To quantify the overlap between the scL-score derived clusters and cNMF clusters, we computed the pairwise overlap between every scL-score cluster and every cNMF cluster and quantified each comparison using the Jaccard Index and Adjusted Rand Index. For each scL-score cluster, we used the maximum value observed across its 7 possible cNMF cluster comparisons, thereby capturing the strongest correspondence between each scL cluster and the cNMF-defined programs for a given metric. To establish a baseline for these overlap measures, we designed a reference simulation by preserving the same cNMF clusters while defining a permuted set of scL clusters obtained by randomly assigning genes to clusters of the same number (7 clusters) and set of sizes (average 24 genes) as the scL clusters. As above, for each permuted scL cluster and each metric, we retained only the maximum overlap value across 7 possible cNMF cluster comparisons. We note that under this framework, the same cNMF cluster can serve as the highest-overlap comparison for more than one scL cluster.

### Calculating the spatial L-score

The spatial L-score was computed analogously to the scL-score except that rather than rank ordering the expression of genes in individual cells, we instead fitted a KDE over the detected spots for a given gene and evaluated it on a square grid of ’spatial bins’ that spanned the entire gastruloid (as described above), then rank-ordered these spatial bins by the resulting probability densities of expression. We treated these bins the same way we had treated cells in the previous calculation, so that the resulting L value reflected mutually exclusive expression in space rather than per-nucleus. The length scale of the interactions considered was determined by the chosen size of the spatial bin. Throughout, we report values for bins at 2 times the nearest-neighbor distance (which was calculated on a per-gastruloid basis). We did not filter low-expressing bins the way we had for cells for 2 reasons: 1) we calculated the KDE on all spots detected, not just those that fell within a nuclear area, so there were fewer areas of low expression, and 2) smoothing resulted in a non-zero density throughout the imaging area. Just as for the scL-score, we symmetrized the spatial L-score by averaging the 2 values calculated for each gene pair, except when displaying the values as a heatmap.

### Evaluation of cell type specific gene clustering in scL-score trees

To determine how well genes associated with specific cell types clustered within a given tree produced by clustering L-score vectors, we first created a measure that we called ‘cell type dispersion’. For all the genes associated with a given cell type, we calculate the average pairwise cophenetic distance within the tree, with the expectation that a tree topology that closely clusters cell type specific genes will produce a lower value than one which spreads cell type specific genes widely among the leaves. To summarize a specific tree, values were averaged across cell types to produce a single value. To test for significance, a permutation-based bootstrap test was performed with 10,000 permutations. p < 0.01 was considered significant.

### Assessment of L-score cluster robustness

To quantify how robust the clusters produced by the L-score were to which genes were used to cluster, we randomly selected subsets of the panel from 60-180 genes, and performed the same gene dispersion analysis described in the section titled ‘Evaluation of cell type specific gene clustering in scL-score trees’. The true values and distributions were compared for each subset, with the full panel of 202 genes well-detected in all gastruloids included as a reference.

### Analysis of single-cell RNA-seq data

Single-cell RNA-seq data from (van den Brink et al., 2020) of pooled 120 hour gastruloids made from the LfngT2AVenus cell line were downloaded from GEO (GSE123187). 4 individual samples were pooled together to generate a dataset of 19075 genes expressed in 14304 cells. Cells with <6000 detected genes were removed, as were genes detected in <3 cells. Cells with >5% mitochondrial counts were also discarded. After this filtering, scL-score values were calculated as previously described, and hierarchical clustering was performed using the Euclidean Distance between scL-score vectors for each gene pair as a distance measure.

This clustering was first performed only using genes common to the panel in this work (207), and then on the entire dataset. Clusters along the diagonal of the resulting matrix were chosen by hand.

### Benchmarking

To demonstrate the effects of noise and expression levels, and to compare to existing methods of calculating exclusivity and coexpression, five scenarios were simulated: (1) two ubiquitously expressed genes with similarly high mean expression, (2) two ubiquitously expressed genes with differing mean expression, (3) two ubiquitously expressed genes with similarly low mean expression, (4) two genes coexpressed across a subset of cells, and (5) two genes expressed in generally opposite subsets of cells. For scenarios 1–3, two genes were independently sampled across 50 cells from Poisson distributions with mean expression set to 100 or 50 (high or low average expression, respectively), with no expression bias toward any cell subset. For scenarios 4 and 5, gene 1 was sampled across 50 cells from a Poisson distribution with mean expression set to 4; gene 2 was then generated by sampling from cell-specific Poisson distributions parameterized to either generally match or oppose the transcript count of gene 1 in each cell.

The scL-score, Exclusive Expression Index (EEI) (Nakajima et al., 2021), and COEX (coexpression coefficient, as calculated by the COTAN package) (Galfrè et al., 2021) were determined for the gene pair in each simulation. As further comparison on real data, the three values were also calculated on 4 representative gene pairs from 2025-05-07_roi2 (Pax6-Eogt, Rfx4-Eogt, Nkx1-2-Rfx4, Cdx4-Cdx2).

### Differential gene expression in endothelial subtypes

Nuclei associated with subpopulations were defined spatially by defining a horizontal cutoff along the AP axis in one gastruloid. The cells typed as endothelial above and below this line were compared to one another. Gene expression values were normalized to Eef2 expression (with a pseudocount of 1% of the mean Eef2 expression added to avoid division by zero). Genes were retained for testing if they were detected (≥2 counts) in at least 50% of cells in one or both groups. Differential expression between groups was assessed using Welch’s t-test on the normalized expression values, and p-values were corrected for multiple comparisons using the Benjamini-Hochberg procedure to control the false discovery rate. Genes with an adjusted p-value <

0.05 were considered differentially expressed. Log2 fold changes were calculated as the ratio of mean normalized expression between groups.

## Acknowledgements

The authors acknowledge Evan Underhill, Harry McNamara, and Jared Toettcher for the generous gift of cell lines, protocols, and training, Sedona Murphy for sample prep and panel design help, Sedona Murphy, Harry McNamara, Grant Kinsler, and Vinay Ayyappan for helpful discussions, and all members of the Raj lab for emotional support and feedback on the manuscript.

C.T. is funded by a Damon Runyon Postdoctoral Fellowship (DRG-2465-22). P.S. acknowledges support from the Roy and Diana Vagelos Scholars Program in the Molecular Life Sciences and the Roy and Diana Vagelos Science Challenge Award. A.R. acknowledges support from a center grant from the Mark Foundation for Cancer Research, NIH Director’s Transformative Research Award R01 GM137425, NIH R01 CA238237, NIH R01 CA232256, and NIH 4DN U01 DK127405.

## Supplementary Tables

**Table S1.** Average nearest neighbor distances between cell centroids in each gastruloid in our dataset from 05/07/2025 (n=18 gastruloids). These values multiplied by 2 were used as bandwidths for individual two-dimensional Gaussian kernels fitted to each RNA spot to produce a smoothed spatial profile of expression for a given gene on a particular gastruloid.

| Gastruloid from 05/07/2025 dataset | Average nearest neighbor distance between cell centroids (μm) |
| --- | --- |
| roi1 | 6.46 |
| roi2 | 6.24 |
| roi3 | 5.61 |
| roi4 | 6.10 |
| roi5 | 6.35 |
| roi6 | 5.93 |
| roi7 | 5.69 |
| roi8 | 5.98 |
| roi9 | 6.58 |
| roi10 | 5.69 |
| roi11 | 5.96 |
| roi12 | 7.01 |
| roi13 | 5.54 |
| roi14 | 6.88 |
| roi15 | 6.80 |
| roi16 | 6.81 |
| roi17 | 5.71 |
| roi18 | 5.45 |

**Table S2.**
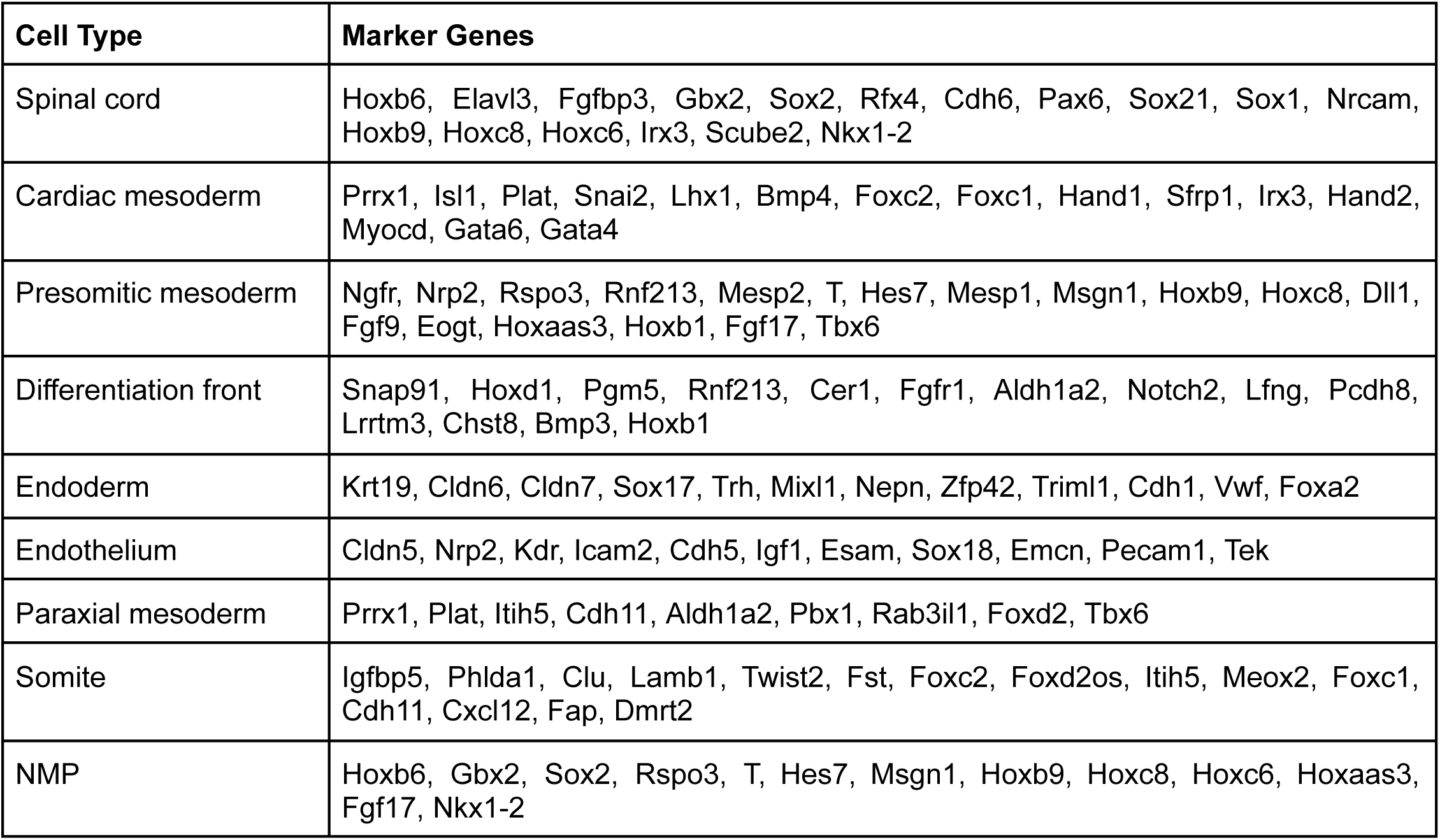
Marker genes for all scored cell types. Genes that mark more than one cell type are shown with both.

| Cell Type | Marker Genes |
| --- | --- |
| Spinal cord | Hoxb6, Elavl3, Fgfbp3, Gbx2, Sox2, Rfx4, Cdh6, Pax6, Sox21, Sox1, Nrcam, Hoxb9, Hoxc8, Hoxc6, Irx3, Scube2, Nkx1-2 |
| Cardiac mesoderm | Prrx1, Isl1, Plat, Snai2, Lhx1, Bmp4, Foxc2, Foxc1, Hand1, Sfrp1, Irx3, Hand2, Myocd, Gata6, Gata4 |
| Presomitic mesoderm | Ngfr, Nrp2, Rspo3, Rnf213, Mesp2, T, Hes7, Mesp1, Msgn1, Hoxb9, Hoxc8, Dll1, Fgf9, Eogt, Hoxaas3, Hoxb1, Fgf17, Tbx6 |
| Differentiation front | Snap91, Hoxd1, Pgm5, Rnf213, Cer1, Fgfr1, Aldh1a2, Notch2, Lfng, Pcdh8, Lrrtm3, Chst8, Bmp3, Hoxb1 |
| Endoderm | Krt19, Cldn6, Cldn7, Sox17, Trh, Mixl1, Nepn, Zfp42, Triml1, Cdh1, Vwf, Foxa2 |
| Endothelium | Cldn5, Nrp2, Kdr, Icam2, Cdh5, Igf1, Esam, Sox18, Emcn, Pecam1, Tek |
| Paraxial mesoderm | Prrx1, Plat, Itih5, Cdh11, Aldh1a2, Pbx1, Rab3il1, Foxd2, Tbx6 |
| Somite | Igfbp5, Phlda1, Clu, Lamb1, Twist2, Fst, Foxc2, Foxd2os, Itih5, Meox2, Foxc1, Cdh11, Cxcl12, Fap, Dmrt2 |
| NMP | Hoxb6, Gbx2, Sox2, Rspo3, T, Hes7, Msgn1, Hoxb9, Hoxc8, Hoxc6, Hoxaas3, Fgf17, Nkx1-2 |

## Author contributions

C.T. performed experiments, C.T. and P.S. performed analysis and prepared figures, Y.H., P.S., and A.R. developed the L-score and wrote associated code, C.T. and A.R. wrote the manuscript, and all authors read and approved the manuscript.

## Declaration of generative AI and AI-assisted technologies

During the preparation of this work, the authors used Claude to generate and improve code for analyses and to suggest improvements to the clarity and flow of the text. The authors reviewed and edited the content and take full responsibility for the content of the manuscript.

## Data and code availability

All code used to process the raw data and generate figures, as well as the raw spot counts, processed data and figures can be found at the following link : https://www.dropbox.com/scl/fo/121330490ladgamapn41b/AObaHONAA7TIwENjYmXh9EM?rlkey=kf5ye1r9p9fx7t1ndeistpc51&dl=0

Additional custom scripts used to process the raw seqFISH data can be found on github: https://github.com/arjunrajlaboratory/NimbusImage/

The code for calculating the L-score can be found on github: https://github.com/arjunrajlaboratory/l-metric

Images of all gastruloids generated for this study, as well as single channel seqFISH images with segmentation and annotations are available here: https://app.nimbusimage.com/#/project/69d3fa8f1be4701f5fab6359

Raw seqFISH images are available upon request.

